# Strategic Modulation of Polarity and Viscosity Sensitivity of Bimane Molecular Rotor-Based Fluorophores for Imaging α-Synuclein

**DOI:** 10.1101/2025.01.07.631748

**Authors:** Yarra Venkatesh, Karthik B. Narayan, Tobias Baumgart, E. James Petersson

**Author notes:** E-mails: Y.V., E.J.P.

## Abstract

Molecular rotor-based fluorophores (RBFs) that are target-selective and sensitive to both polarity and viscosity are valuable for diverse biological applications. Here, we have designed next-generation RBFs based on the underexplored bimane fluorophore through either changing in aryl substitution or varying π-linkages between the rotatable electron donors and acceptors to produce red-shifted fluorescence emissions with large Stokes shifts. RBFs exhibit a twisted intramolecular charge transfer mechanism that enables control of polarity and viscosity sensitivity, as well as target selectivity. These features enable their application in: (1) turn-on fluorescent detection of α-synuclein (αS) fibrils, a hallmark of Parkinson’s disease (PD), including amplified fibrils from patient samples; (2) monitoring early misfolding and oligomer formation during αS aggregation; and (3) selective imaging of αS condensates formed by liquid-liquid phase separation (LLPS). In all three cases, we show that our probes have high levels of selectivity for αS versus other aggregating proteins. These properties enable one to study the interplay of αS and tau in amyloid aggregation and the mechanisms underlying neurodegenerative disorders.

## Introduction

Fluorescence imaging is a crucial tool for studying complex biological pathways, diagnosing diseases, and gaining detailed insights into molecular and cellular processes.^1-3^ This technique requires precise labeling of fluorescent probes to visualize biological entities and track interactions in real time. Over the past two decades, several labeling strategies have been developed that include fluorescent proteins like GFP and genetically encodable tags (e.g., HaloTag, SnapTag, Spinach).^4-6^ However, these often have limitations due to their large size that can interfere with the native function of the biological molecules that are intended to study.^7^ Small organic dyes can be directly attached to proteins or DNA/RNA through chemical conjugation methods, but these methods are either restricted to *in vitro* or require manipulation of cellular machinery through methods like genetic code expansion, making them hard to apply in native biological samples.^1^ To overcome these limitations and enable direct fluorescence imaging without need for extensive labeling strategies, there is a need to develop fluorescent probes that are highly target selective.

Molecular rotor-based fluorophores (RBFs) using the twisted intramolecular charge transfer (TICT) mechanism find applications in biological imaging, membrane chemistry, and material science.^8-10^ Typically, RBFs feature an electron donor, an electron acceptor, and a linkage that facilitates charge transfer. In solvents that allow rotation between the donor and acceptor, these fluorophores adopt twisted conformations in excited states, leading to rapid intramolecular charge transfer (ICT) and effective fluorescence quenching. Restricted rotation enhances fluorescence by reducing non-radiative decay, making RBFs valuable as biosensors in viscous environments.

Recently, small molecule-based RBFs have emerged as powerful tools to study protein misfolding and aggregation, which can occur due to genetic mutations, post-translational modifications, exogenous toxins, and cellular stresses. Protein misfolding and amyloid fibril formation has been associated with Alzheimer’s disease (AD), Parkinson’s disease (PD), and many other neurodegenerative disorders.^11^ Thioflavin-T (ThT), an exemplar RBF, is widely used for detection of a variety of protein aggregates.^12, 13^ However, ThT and many other RBFs primarily turn on in the presence of mature fibrils, and are weakly fluorescent with early aggregation species such as protein condensates or non-fibrillar oligomers.^14-17^ Detecting these early species is critical for understanding the full scope of protein aggregation pathology.

In a recent development, Zhang and co-workers introduced RBFs derived from ThT with extended π-rich linkages (**Figure 1a**).^18^ This innovation enabled the control of viscosity sensitivity by changing the rotational energy barrier (*E*_a_) thereby inducing fluorescence turn-on through a restricted TICT mechanism, enabling selective imaging of protein oligomers and aggregates using their AggTag method (where Halo-tag is fused to a protein of interest and labeled with Halo ligand modified attached to the fluorophore).^19^

**Figure 1.**
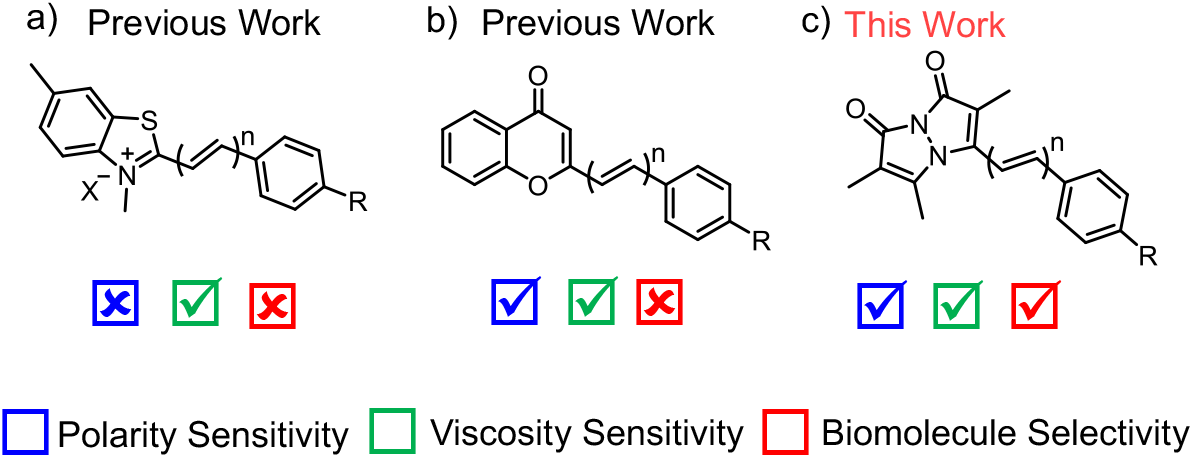
Molecular rotor-based fluorophores (RBFs). (a) Benzothiazolium-based RBFs with viscosity sensitivity. (b) Chromone-based RBFs with dual sensitivity to polarity and viscosity. Here “n” denotes π-linker length between donor and acceptor moieties, n = 0, 1, and 2 and R = NMe_2_. (c) Bimane-based RBFs exhibit dual sensitivity to polarity and viscosity with high biomolecule selectivity. Here n = 1 or 2 and R = NH_2_, NHMe, NMe_2_, azetidine, and morpholine

**Scheme 1.**
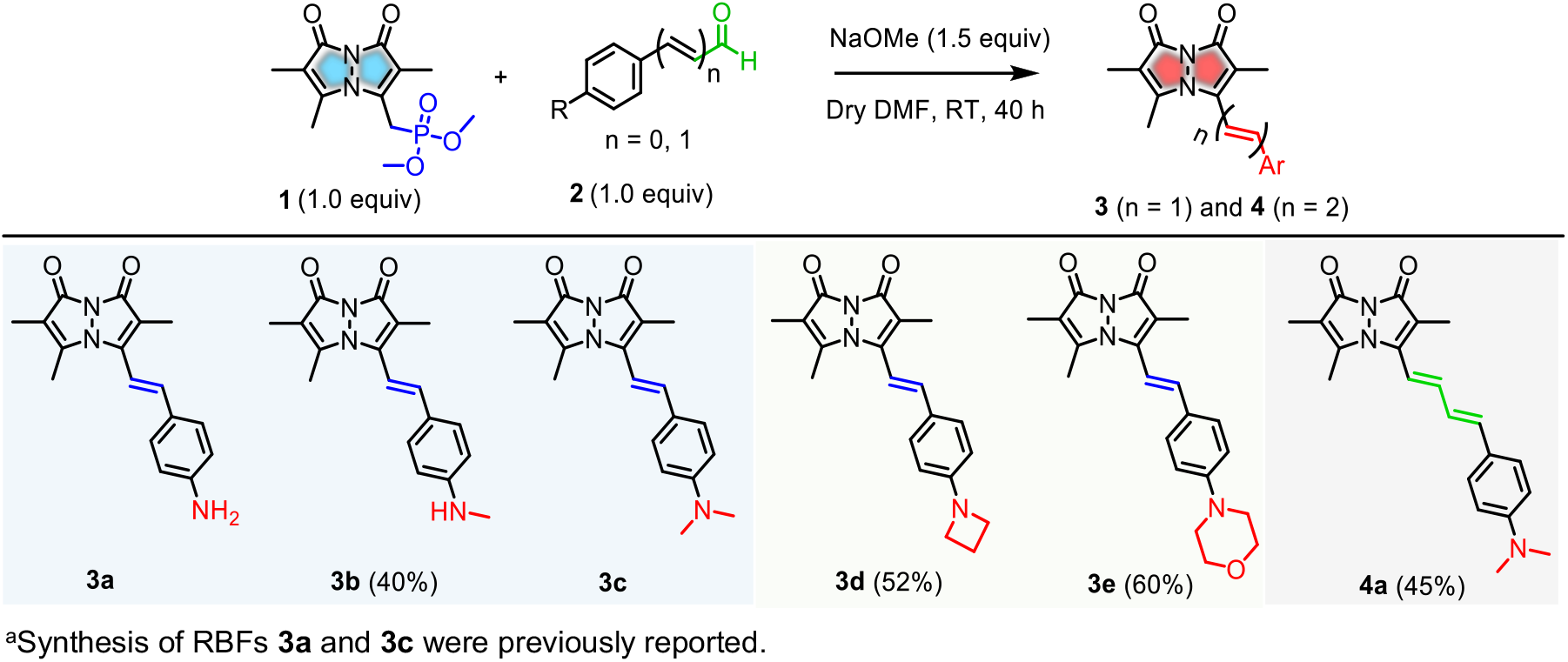
Design and Synthesis of RBFs 3a-e and 4a with Either Varied Aryl Substituents or Extended π-Rich Linkages^a^.

Subsequently, Liu’s laboratory demonstrated that polarity sensitivity could be regulated by extending the conjugation length derived from a chromone acceptor moiety (**Figure 1b**).^20^ This approach controlled both polarity and viscosity sensitivity through change in ICT and *E*_a_, respectively. The resulting probes offered insights into the polarity of aggregated proteins *via* AggTag labeling in live cells. Despite the advancements of the AggTag method, the large size of Halo-Tag has the potential to disrupt disordered amyloidogenic proteins like α-synuclein (αS) and tau.^21, 22^ A key challenge in studying αS aggregation is the lack of tools to study early species, such as those involved in misfolding and liquid-liquid phase separation (LLPS),^23-25^ limiting their application in early disease diagnosis. The sensitivity of RBFs developed by Zhang’s lab is restricted to oligomeric species and fibrils. To detect low-viscosity species like early aggregates or LLPS, novel fluorescent probes with adjustable sensitivity are needed. Additionally, the lack of intrinsic protein selectivity requires the use of Halo-tag and limits studying unmodified proteins in neurodegenerative diseases that involve co-aggregation of multiple amyloidogenic proteins.

In this study, we report the chemical modification of RBFs based on the underexplored bimane fluorophore by either changing the aryl substituent or installing π-rich bridges between the two rotational moieties (**Figure 1c**). These chemical modifications demonstrate new mechanisms to control RBF polarity sensitivity through enhanced ICT character and to control viscosity sensitivity by altering *E*_a_ for the transition between the fluorescent planar excited state and the dark twisted state. The modifications also enhance photophysical properties, such as red-shifting absorption in the visible region. These features enable use as “turn-on” fluorescent probes to detect αS fibrillar aggregates, a hallmark of PD.^26^ Structural variations in RBFs improve selectivity for αS fibrils versus other amyloidogenic proteins. This selectivity enables detecting αS fibrils amplified from patient samples. Finally, the sensitivity of our RBFs to low viscosity early aggregation species is used in fluorescence imaging of αS protein LLPS condensates.

## Results and Discussion

### Rational Design of RBFs

RBFs **3a** (R = NH_2_, n = 1) and **3c** (R = NMe_2_, n = 1) are stilbene-type analogs that feature a methine group between the electron-donating aryl group and the electron-accepting bimane fluorophore (D-π-A system). Recently, they have been demonstrated as selective fluorogenic probes for detecting amyloid fibrils.^27^ To further investigate the bimane-based D-π-A system, we modified either the aryl substituent (R) or the π-linker length (n) between the donor and acceptor moieties. To generalize this system, we introduced substituents with varying electron-donating capacities, such as **3b** (R = NHMe, n = 1), along with heterocyclic amines like **3d** (R = azetidine, n = 1) and **3e** (R = morpholine, n = 1). Next, we anticipated that extending the methine bridge, as in **4a** (R = NMe_2_, n = 2), would result in a second-generation bimane probe with improved photophysical properties within the same D-π-A framework. RBFs **3b, 3d, 3e**, and **4a** were synthesized via the Horner–Wadsworth–Emmons (HWE) reaction between phosphonate **1** and commercially available aryl aldehydes (**2b, 2d, 2e**, and **2c**), following a previously described procedure, in 40-60% yields (**Scheme 1**). These compounds were characterized by ^1^H, ^13^C NMR, and high-resolution mass spectrometry (HRMS) analysis (**Figures S1-S4**).

After synthesizing RBFs with varied aryl substituents or linker lengths, we performed photophysical characterization using absorption and fluorescence spectroscopy, comparing them to **3a** and **3c** in a 1:1 mixture of acetonitrile (ACN) and phosphate buffered saline (PBS). As shown in **Figure 2a-b** and **Table 1**, a bathochromic shift in absorption maximum (λ_abs_) and emission maximum (λ_em_) was observed with increased electron donation: **3a** (R = NH_2_, λ_abs_ = 385 nm, λ_em_ = 583 nm), **3b** (R = NHMe, λ_abs_ = 399 nm, λ_em_ = 591 nm), and **3c** (R = NH_2_, λ_abs_ = 418 nm, λ_em_ = 604 nm). These experimental λ_abs_ and λ_em_ values align well with density functional theory (DFT) and time-dependent DFT (TD-DFT) calculations for the ground and excited states, respectively. As shown in **Table 1** (and **Figure 2c**), the HOMO–LUMO energy gap (*E*) decreases between **3a** (R = NH_2_, 3.429 eV) to **3c** (R = NMe_2_, 3.172 eV), along with a decrease in fluorescence emission energy from **3a** (2.40 eV) to **3c** (2.32 eV). Interestingly, RBF **3d** (R = Az, λ_abs_ = 403 nm) showed a hypsochromic shift in λ_abs_ compared to **3c** (R = NMe_2_, λ_abs_ = 418 nm), and an even greater blue-shift seen for **3e** (R = Mor, λ_abs_ = 386 nm), can be explained by the decreased electronic conjugation caused by the rigid structures of the azetidine (Az) and morpholine (Mor) groups (**Figure S5**).

**Table 1.**
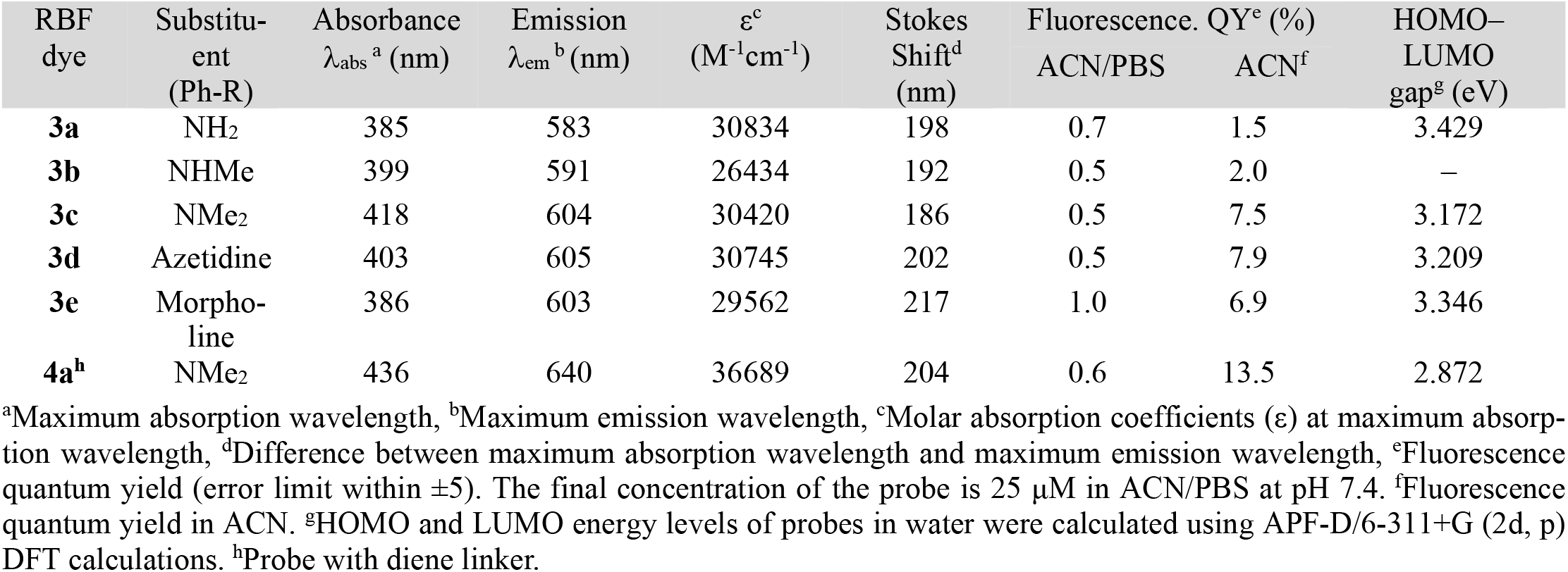
Photophysical properties of RBFs **3a**-**3e** and **4a**.

**Figure 2.**
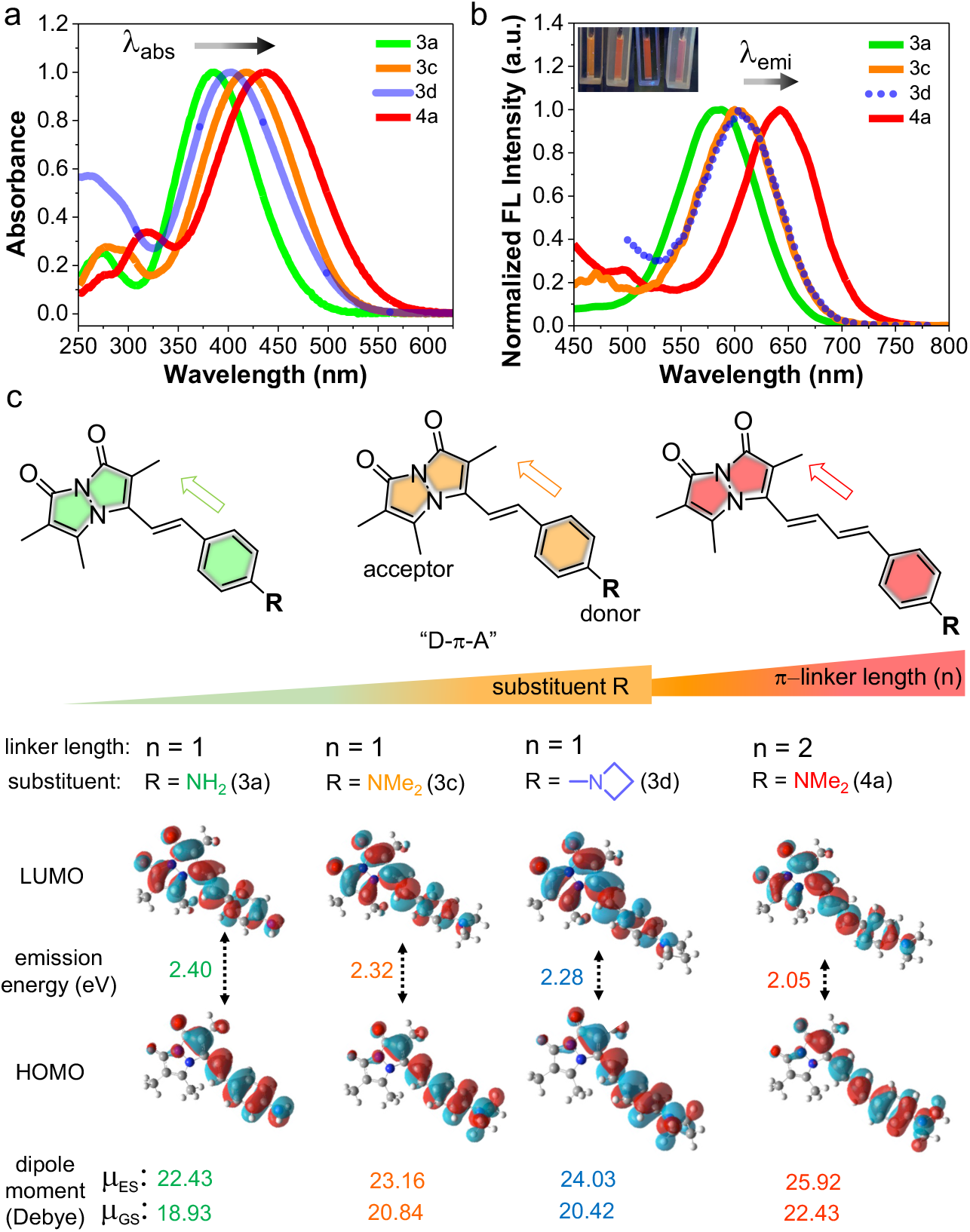
Photophysical properties of RBFs with variations in the aryl substitution (R) or π-linker length (n). (a) Normalized absorption spectra and (b) Normalized fluorescence spectra of **3a, 3c, 3d** and **4a**, measured with excitation at their optimal max absorption. Inset: photographs of probes (25 μM) under a UV lamp at 365 nm in an ACN/PBS buffer (50:50 v/v) at pH 7.4. (c) Top: schematic representation of the bimane probes showing “push–pull” D–π–A type systems; D: donor, π: spacer, and A: acceptor. Bottom: TD-DFT calculated fluorescence emission energy from the excited S_1_ state (ES) back to the ground S_0_ state (GS) of **3a, 3c, 3d**, and **4a** in water and their calculated dipole moments at their GS and ES in water, supports the enhanced ICT character from **3a** to **3c-d** to **4a**.

The reduced conjugation in the cyclic substituents results in a larger HOMO-LUMO energy gap for **3d** (*E* = 3.209 eV) and **3e** (*E* = 3.346 eV) than **3c** (R = NMe_2_, *E* = 3.172 eV). Despite the reduced conjugation affecting the ground-state electronic structure, **3d** (R = Az) showed a red-shifted λ_em_ at 605 nm compared to **3c** (R = NMe_2_, 604 nm). This is further supported by TD-DFT calculations, which show a lower emission energy for **3d** (*E*_em_ = 2.28 eV) than for **3c** (*E*_em_ = 2.32 eV). Similarly, **3e** (R = Mor) also exhibited a red-shifted λ_em_ at 603 nm, reflecting the impact of the structural rigidity on the excited-state properties.^28, 29^ Excitingly, RBF **4a** with extended linker (R = NMe_2_, n = 2) exhibited highly red shifted λ_abs_ at 436 nm and λ_em_ at 640 nm. Again, electronic structure calculations supported our design, showing the lowest HOMO-LUMO energy of *E* = 2.87 eV and emission energy of *E*_em_ = 2.05 eV. In general, the experimental photophysical properties agree well with calculated values, as depicted in **Table S2-S3** and **Figure S7**. The calculated Stokes shifts for **3a-3c** ranged from 186 to 198 nm (**Table 1**). Notably, **3d** (R = Az) and **3e** (R = Mor) exhibited larger shifts of 202 nm and 217 nm, respectively, while **4a** showed a shift of 204 nm. The tabulated photophysical properties of **3a-e** and **4a** in ACN show blue-shifted λ_abs_ and λ_em_ relative to values in ACN/PBS (**Figure S6** and **Table S1**). Overall, these tunable photophysical properties of RBFs make them promising probes for various biological applications.

Fluorescence quantum yields (QYs) of **3a-e** and **4a** in ACN/PBS ranged from 0.5-1.0%, compared to 64.6% for phosphonate **1**, due to the TICT effect in polar protic solvents. In contrast, in ACN, QYs increased with the electron-donating ability of the aryl substituent: 1.5% for **3a** (R = NH_2_), 2.0% for **3b** (R = NHMe), and 7.5% for **3c** (R = NMe_2_). For **3d** (R = Az), the QY rose to 7.9%, likely due to restricted twisting that stabilizes the excited state and reduces TICT formation compared to the **3c** (R = NMe_2_). Similarly, **3e** (R = Mor) showed a QY of 6.9%. Notably, **4a** (R = NMe_2_, n = 2) reached 13.5%, suggesting an even stronger fluorescence turn-on (**Table 1** and **Figure S8**). We assume that modifying the aryl substitution or linker length could control the fluorogenic behavior of RBFs by enhancing *E*_a_ or enabling multiple modes of rotation and isomerization.

### Varying the Aryl Substituent or Extended π-Rich Linkages Modulates Polarity Sensitivity of RBFs

We previously demonstrated that probes **3a** and **3c** exhibit polarity and viscosity sensitivity *via* ICT and TICT mechanisms, respectively (**Figure 3a-b**). Similarly, RBFs **3b** (486–594 nm), **3d** (493–604 nm), **3e** (489–603 nm), and **4a** (532–640 nm) display strong solvatochromism, with emission wavelengths ranging from 486–640 nm depending on solvent polarity (**Figure S9**). **Figure 3c** depicts the strict linearity between the λ_em_ of **3a, 3c**, and **4a** and dielectric constant of solvent, which can enable quantitative analysis of microenvironmental polarity on a protein surface or within cellular compartments. To examine the impact of aryl substituents and linker length on polarity sensitivity, we measured the emission of **3a–3e** and **4a** across solvents with varying dielectric constants (**Figure 3c** and **Figure S10**). Polarity sensitivity increased with stronger electron donation, as observed in **3a** (26.7), **3b** (30.7), and **3c** (32.1), attributed to the ICT mechanism driven by the dipole moment between electron-donating dialkylaminophenylene groups (R = NH_2_, NHMe, NMe_2_) and the electron-withdrawing bimane moiety (**Figure 3d**). This trend was confirmed by electrostatic potential (ESP) mapping and excited-state dipole moment calculations, showing increased sensitivity from NH_2_ to NMe_2_ (22.43 for 3a, 23.16 for 3c) (**Figure 2c** and **Figure S11**).

**Figure 3.**
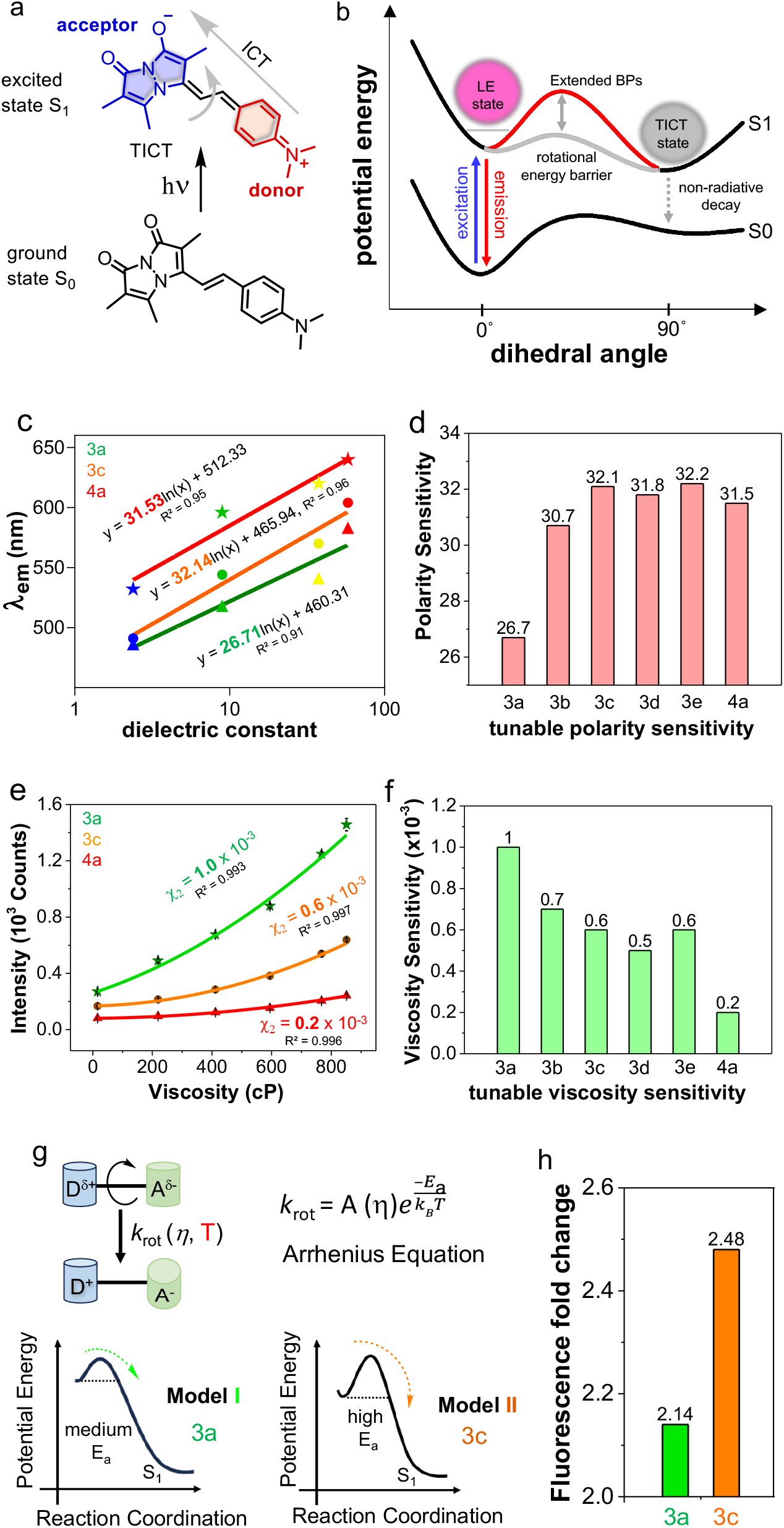
Control of polarity and viscosity sensitivity of RBFs. (a) RBFs with D–π–A system shows the phenomenon of ICT from the donor to the acceptor in the excited state. (b) Jablonski diagram illustrates that an excited fluorophore could return to the ground state either through fluorescence or internal rotation, leading to non-radiative TICT. It shows the effect of aryl substitution (R) or π-rich linkage length (n) to enhance the inherent rotational energy barrier (*E*_a_) of RBFs. (c) Quantitative polarity sensitivity analysis of **3a, 3c**, and **4a**. (d) Tunabe polarity sensitivity of **3a-e** and **4a**. (e) Quantitative viscosity sensitivity analysis of **3a, 3c**, and **4a**. (f) Tunable viscosity sensitivity of **3a-e** and **4a**. (g) Temperature dependence reflects the magnitude of *E*_a_. Proposed models suggest that heights of *E*_a_ determine the fluorescence intensity pattern in the solvents with different viscosity at different temperatures. Here two models were conceptualized with moderate and high energy barriers for **3a** and **3c**, repectively. (h) Viscosity depedence fluorescence fold change of **3a** and **3c** at the temperatures of 25° C and 42 °C. Final concentration of probes is 25 μM in indicated solvents or methanol/glycerol mixtures.

RBF **3d** exhibited lower sensitivity (31.8, μ = 20.42 D at ground state) compared to **3c** (32.1, μ = 20.84 D at ground state), due to rigidity and limited electron donation of the Az ring, while **3e** with a Mor group showed slightly higher sensitivity (32.2) relative to **3c**, driven by the Mor oxygen’s high electronegativity. ESP mapping further confirmed negative electrostatic potential on the Mor oxygen atom that contributes to the increased sensitivity (**Figure S11**). Interestingly, RBF **4a**, with an extended linker, showed similar overall sensitivity (∼31.5, μ = 25.92 D at excited state) but was more sensitive to solvents with low dielectric constants and less sensitive to those with high dielectric constants. Its solvatochromism was notable, showing a 64 nm λ_em_ shift for **4a** compared to 32–57 nm for **3a-e** across solvents with dielectric constants from 2.38 to 8.93. In high-dielectric solvents, **4a** exhibited minimal sensitivity (20 nm λ_em_ shift) compared to **3a-e** (29–42 nm λ_em_ shift) across dielectric constants from 37.5 and 58 (**Figure S12**).

### Varying the Aryl Substituent or Extended **π**-Rich Linkages Modulate Viscosity Sensitivity of RBFs

We next examined viscosity sensitivity, which quantifies the relationship between fluorescence intensity and viscosity, for the newly synthesized RBFs **3b, 3d, 3e**, and **4a**, as well as the previously known compounds **3a** and **3c**. Fluorescence intensity was measured in binary mixtures of ethylene glycol and glycerol (EG/G) with known viscosities ranging from 16 to 851 centipoise (cP). An increase in glycerol concentration leads to an increase in fluorescence intensity and χ_2_ values were derived for each RBF using second-order polynomial regression. Higher χ_2_ values indicate higher fluorescence intensity changes as a function of viscosity. RBFs exhibited decreased χ values with increasing electron donation, as observed χ_2_ value of 1.0 ×10^−3^ for **3a** (R = NH_2_) with 5.4-fold fluorescence intensity change, 0.7 ×10^−3^ for **3b** (R = NHMe, 4.68-fold), 0.6 ×10^−3^ for **3c** (R = NMe_2_, 3.82-fold) (**Figure 3e-3f, Figure S13-15**, and **Table S5**). This indicates that the restriction of TICT is the primary cause of their fluorogenic response and sensitivity. Moreover, we observed that **3c** was exclusively fluorescent in crystalline or amorphous solids (**3b, 3d, 3e**, and **4a**). This indicates that **3c** (CCDC: 2240198) requires well-ordered π-π stacking (J aggregates, head-to-tail packing) to suppress vibrational or rotational motions and activate fluorescence (**Figure S16**).^27^

We observed a decreased χ_2_ value of 0.5 ×10^−3^ for **3d** (R = Az), yet it exhibited a higher fluorescence turn-on (4.07-fold) compared to **3c** (χ_2_ = 0.6 ×10^−3^, 3.82-fold). This is attributed to the rigid Az group in **3d**, which restricts non-radiative decay pathways (TICT), unlike the more flexible NMe_2_ group in **3c**, which allows for greater non-radiative decay. This is reflected in the observed fluorescence QYs, with **3d** at 7.9%, slightly higher than **3c** at 7.5%. The decrease in the χ_2_ value for **3d** compared to **3c** can be explained by the calculated emission energy and red-shifted emission wavelength of **3d**, which are 2.28 eV and 605 nm, respectively, compared to 2.32 eV and 604 nm for **3c**.

This suggests that the Az group acts as a better electron donor in the excited state. Again, **3e** (R = Mor) showed a decreased χ_2_ value of 0.6 ×10^−3^ with a decrease in the fluorescence of 3.81-fold compared to **3d**, similar to **3c**. This is due to the Mor group having a six-membered ring with an oxygen atom, providing slightly more flexibility to undergo non-radiative pathways compared to the Az group. Interestingly, **4a** (R = NMe_2_) with an extended linker, showed the lowest fluorescence change, around 3-fold with χ_2_ = 0.2 ×10^−3^ (**Figure 3e-f**), suggesting that not only varying aryl substituent but also incorporating π-rich linkers may significantly affect non-radiative pathways by exhibiting a unique photochemical mechanism.

Our efforts next focused on understanding the mechanism regulating the viscosity sensitivity of RBFs by varying the substituent and π-rich linkages. Zhang *et al*. demonstrated that π-rich linkages alter the internal rotation rate (*k*_rot_) between planar and twisted configurations of RBF.^18^ Building on this, we hypothesized that substituent variations could further influence *k*_rot_. The Arrhenius equation provides a framework for understanding the temperature and viscosity dependence of the fluorescence intensity of RBFs. We conceptualized two models: Model I with a moderate rotational energy barrier (*E*_a_ > k_B_T) for **3a** (R = NH_2_), predicting moderate fluorescence changes when measured in methanol and glycerol mixtures at different temperatures and Model II with high *E*_a_ (≫k_B_T) for **3c** (R = NMe_2_), predicting larger fluorescence changes (**Figure 3g, Figure S17**). To validate this hypothesis, we measured the fluorescence intensity of RBFs at varying temperatures and viscosities, selecting **3a** and **3c** to test the effects of their substituents on *E*_a_. As shown in **Figure 3h** and **Figure S17, 3c** exhibited a 2.48-fold fluorescence change between 25 °C and 42 °C, compared to 2.14-fold for **3a**, aligning with Model I for **3a** and Model II for **3c**. RBFs with high *E*_a_ retain greater fluorescence by inhibiting rotational nonradiative decay, as reflected in QY in ACN: **3a** (1.5%), **3c** (7.5%), and **4a** (13.5%) (**Table 1**). Consequently, high *E*_a_ RBFs exhibit lower viscosity sensitivity. Our findings also reveal that substituent variations, alongside extended π-conjugation, influence viscosity sensitivity, providing a framework for designing fluorophores with tunable fluorogenicity.

### Structure Variation Retains Its Binding to **α**S Fibrils with Red Shifted Emission

We previously demonstrated that RBF **3c** (R = NMe_2_) selectively binds αS fibrils with a fluorescent turn-on response. Here, we explored how aryl substituents or linker length variations affect polarity, affinity, and selectivity upon binding to αS fibrils. Similarly, the fluorescence measurements of **3a, 3b, 3d, 3e**, and **4a** with αS fibrils showed turn-on fluorescence with different λ_max_. Interestingly, RBF **4a** showed red emission at 620 nm with a 383-fold brightness increase and a fluorescence lifetime (τ) of 1.27 ns, while **3c** emitted orange fluorescence at 580 nm with a 476-fold increase and a τ of 2.27 ns upon binding to αS fibrils (**Figure 4a-b, Figure S19-21, Table S7**). The color variation is due to polarity changes resulting from RBF structural variation, as observed in TD-DFT calculations (**Figure 2c**). All RBFs showed significant Stokes shifts and enhancements in ε, QY, and τ upon binding (**Table S8**). We next assessed the binding affinities of **3a–e** and **4a**. As shown in **Figure 4c** and **Figure S22**, affinity increased with electron donation from **3a** to **3c**, while **3d** and **3e**, with cyclic amines, showed reduced affinity. Notably, **4a**, with an extended linker (n = 2), displayed a dissociation constant (*K*_d_) of 0.42 ± 0.25 μM—3.5 times higher affinity than **3c** (n = 1), which had a *K*_d_ of 1.45 ± 0.43 µM.

**Figure 4.**
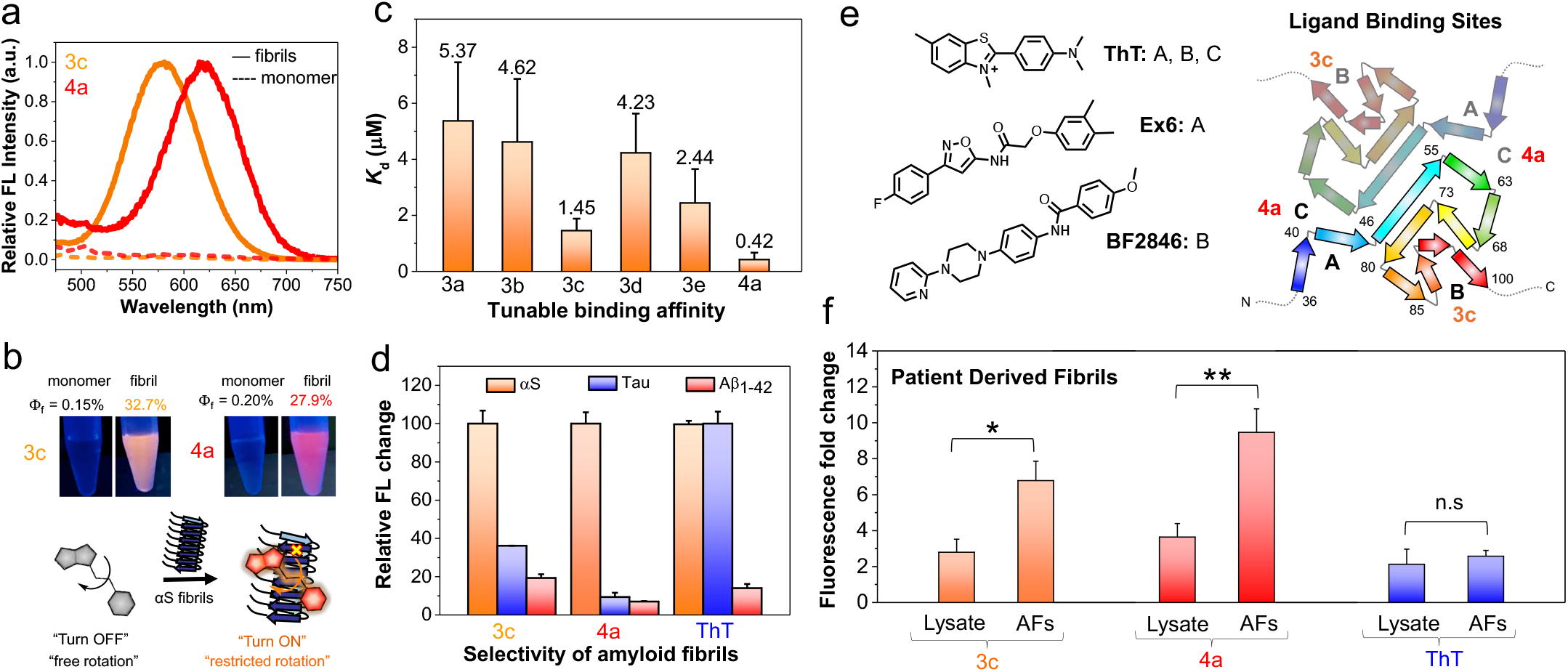
Structural modification affects probe detection of amyloid fibrils. (a) Fluorescence spectral changes of probes **3c** and **4a** (10 μM) in the presence of αS monomer and fibrils (50 μM). (b) Top: Photographs of the **3c** and **4a** (10 μM) in the presence of αS monomer and fibrils (50 μM) under a handheld UV lamp at 365 nm; Bottom: Schematic representation of the “turn on” fluorescence of **3c** and **4a** probes upon binding to αS fibrils, which resulting from the restricted internal rotation. (c) Tunable dissociation constant (*K*_d_) of **3a-e** and **4a** (10 nM–10 μM) with 100 μM αS fibrils. (d) Enhanced amyloid selectivity of the probe **4a** for αS fibrils compared to the **3c** and standard amyloid binding dye ThT (1 μM) among different amyloid fibrils of tau and Aβ_1-42_ that are present in patients with neurodegenerative diseases. (e) Binding sites for known ligands based on cryo-EM (ThT)^30^ and photo-crosslinking (Ex6 and BF2846) shown on top-down view of two-stranded fibril with residues numbered.^31, 32^ (f) Fluorescence fold change of the probes **3c, 4a**, and ThT (1 μM) with brain lysates and AFs (1 μM) derived from PDD patients. The bar graph represents the averaged data based on three different PDD patients derived AFs and lysates. Fold change was subjected to subtraction of free probe fluorescence in PBS buffer. Ex/Em wavelengths: 463/580 nm for **3c**, 484/620 nm for **4a**, 450/482 nm for ThT. Error bars represent the SD of 3 measurements. Statistical analysis: \**p* < 0.01 and \*\**p* < 0.005.

Given the higher binding affinities of **3c** and **4a**, we next explored their key feature—selectivity— making them invaluable tools for imaging biological samples. Therefore, we first evaluated the selectivity in HEK cell lysate (10 mg/mL total protein) spiked with αS fibrils. Probe **4a** showed a 4-fold fluorescence increase, while probe **2a** exhibited an 18-fold increase, demonstrating their selective detection of αS fibrils among other cellular proteins (**Figure S24**). We next assessed the probes’ selectivity for αS fibrils over tau and amyloid-β (Aβ_1-42_), which are prevalent in AD patients’ brains. Distinguishing αS from tau and Aβ_1-42_ is crucial for understanding the overlapping pathology of AD, PD, and other neurodegenerative disorders. Probe **4a** showed significantly higher selectivity for αS fibrils, with 11 times greater turn-on than for tau fibrils and 15 times more than Aβ_1-42_ fibrils, while **3c** exhibited 2.8- and 5.2-fold higher turn-on, respectively. Thus, our probes, especially **4a**, demonstrated superior target protein selectivity compared to ThT (**Figure 4d, Figure S25**).

We sought to elucidate the underlying mechanism for the selectivity of **4a** and **3c** relative to ThT and each other. Cryo-electron microscopy (cryo-EM) structures and computational modeling have revealed that αS fibrils have multiple β-sheet-rich binding pockets with varying affinities for aromatic ligands resembling **3c** and **4a** (**Figure 4e**).^33^ The highest affinity interaction for ThT, which is modeled into the reported cryo-EM structures, involves binding in a pocket formed around Tyr39.^30^ In our efforts to develop positron emission tomography (PET) imaging probes for αS fibrils in PD and multiple system atrophy (MSA), we have observed binding of aromatic ligands to this site as well, as determined by photo-crosslinking and mass spectrometry experiments.^31, 32, 34^ We refer to this site here as Site A. For a different set of ligands, we have observed selective binding to a site formed around Phe94, which we refer to here as Site B.^31, 35^ Site B is identified in initial cryo-EM density as a weaker binding site for ThT as well as B227a and C05-03, aromatic ligands with extended π-linkages similar to our RBF compounds.^30^ The cryo-EM maps also show density for binding of ThT, B227a, and C05-03 at the fibril-fibril interface (Site C).

To better understand the binding of **3c** and **4a**, we performed competition binding studies with ThT, Ex6, and BF2847. Ex6 and BF2847 have been identified as high affinity, selective Site A and Site B binders respectively, based on photo-crosslinking studies and low overlap in competitive binding studies.^31, 33, 34^ We found that both **3c** and **4a** competed with the promiscuous binder ThT, but that some ThT emission remained at saturating concentrations, indicating that there was at least one non-overlapping binding site for each ligand (**Figure S23**). RBF **3c** binding was not displaced by Ex6, but was partially displaced by BF2846 (**Figure S23**). Taken together, these data suggest that the fluorescing **3c** species binds at Site B, but not at Site A, allowing ThT to remain bound at Site A. In contrast, RBF **4a** fluorescence was not affected by either Ex6 or BF2846, implying that the fluorescence species binds at neither Site A nor Site B (**Figure S23**). Note that **4a** must have some binding at these sites in order to displace ThT, but that these do not seem to activate **4a** fluorescence turn-on. Finally, to ensure that **3c** and **4a** did not bind to regions unresolved in the cryo-EM structures, we performed fluorescence titrations with truncated αS_1-100_ fibrils, omitting the C-terminal region (residues 101-140). It has been shown by cryo-EM that this region can be removed without affecting fibril morphology or fibril-fibril packing.^36^ For both **3c** and **4a**, we found similar results to our titrations with full length fibrils (**Figure S28**). Taken together, these results indicate that **3c** and **4a** have several weak binding sites that overlap with ThT, but have well-defined, fluorogenic sites that are not displaced by ThT (**Figure 4e**). For RBF **3c**, this site appears to be Site B. For RBF **4a**, this site appears to be neither Site A nor Site B, but may be the interfacial site identified in cryo-EM density maps for ThT, B227a, and C05-03 (Site C).

Additional analysis of concentration-dependent binding shows λ_em_ for **4a** shifting from 617 to 618 nm and a larger 6 nm shift for **3c** (from 575 to 581 nm) as concentration increased (**Figure S27**), supporting the idea of multiple weak binding sites for **3c**, and one site that is primarily responsible for the fluorescent species for **4a**. By using the fits to the λ_em_ values determined in various solvents (**Figure S10**), one can calculate the dielectric constant of the local environments for **3c** and **4a** bound to αS fibrils. The values obtained for **3c** (35.9 using 581 nm) and **4a** (27.6 using 617 nm) correspond to **3c** binding in a more polar, dimethylformamide-like environment (36.7)^37^ while **4a** binds in an ethanol-like environment (24.6).^37^ This further supports the idea that the two compounds bind at distinct sites, with the **4a** binding site more solvent-exposed.

### 4a Enhances Selectivity for Diagnostic Detection of Amplified Fibrils Derived from Brains of Patients with PDD

We next aimed to explore the clinical potential of **4a**, which shows the greatest *in vitro* selectivity for αS fibrils over tau and Aβ_1-42_ fibrils. Recent studies have identified unique αS fibril polymorphs linked to diseases that differ from *in vitro* forms.^21, 38, 39^ Fibril amplification assays hold promise as PD biomarkers, but ThT cannot detect amplified fibrils (AFs). AFs can be reliably amplified *in vitro* using patient-derived seeds and recombinant αS monomer. We previously showed **3c** can detect AFs in PD with dementia (PDD).^27^ Here, we investigated how structural modification from **3c** to **4a** affects the sensitivity and selectivity for patient-derived αS fibril strains. Notably, **4a** showed 2.6-fold higher fluorescence with PDD-derived AFs (p < 0.005) compared to unamplified material (lysate), while **3c** showed 2.4-fold fluorescence increase (p < 0.01). ThT failed to distinguish AFs from lysate (**Figure 4f**). These results suggest that extending linkages could be a general strategy to improve both sensitivity and selectivity. Specifically, **4a**, with its enhanced selectivity for αS fibrils, shows strong potential for use in a biomarker assay for PD.

### RBFs with Extended Linkages Enable the Early Detection of Low-Viscosity **α**S Misfolding or Oligomer Formation

Current PD treatments focus on alleviating symptoms but do not slow or halt disease progression. A major barrier in PD research is the lack of diagnostic tools for early detection and biochemical monitoring of progression.^40^ αS aggregation is a stepwise process, initiated by monomer misfolding, followed by the formation of low-viscosity soluble oligomers, and ultimately leading to high-viscosity insoluble aggregates (**Figure 5a**).^41, 42^ While many probes reported thus far have focused on detecting mature fibrils, there are some reports that specifically target oligomers, albeit without specificity for αS.^17^ A particular focus on the development of tools to study early aggregation could significantly enhance the understanding of PD progression. Towards addressing this challenge, we monitored αS aggregation using **3c** and **4a** with varying viscosity sensitivity under prolonged agitation. **Figure 5b** shows that probe **4a** exhibited a 15-fold increase in fluorescence intensity within 20 hours, indicating detection of misfolding or oligomers/protofibrils. On the other hand, probe **3c**, although unable to detect changes within 20 h, showed an increase by 40 h, indicating better resolution for later stage oligomers or protofibrils. In contrast ThT, couldn’t detect either misfolding or oligomers/protofibrils accumulating between 0-40 hours, only tracking mature fibrils. Moreover, spectra recorded at different time intervals indicate that **4a** exhibits significant fluorescence as early as 8 hours of aggregation. At this time point, **3c** shows minimal signal, and ThT does not show any fluorescence enhancement (**Figure S29**). These findings suggest RBFs with higher *E*_a_ values (lower χ_2_ value, 4a) detect low viscosity early misfolding or oligomers/protofibrils, while those with lower *E*_a_ values (higher χ_2_ value, 3c) detect later stages and high-viscosity aggregates.

**Figure 5.**
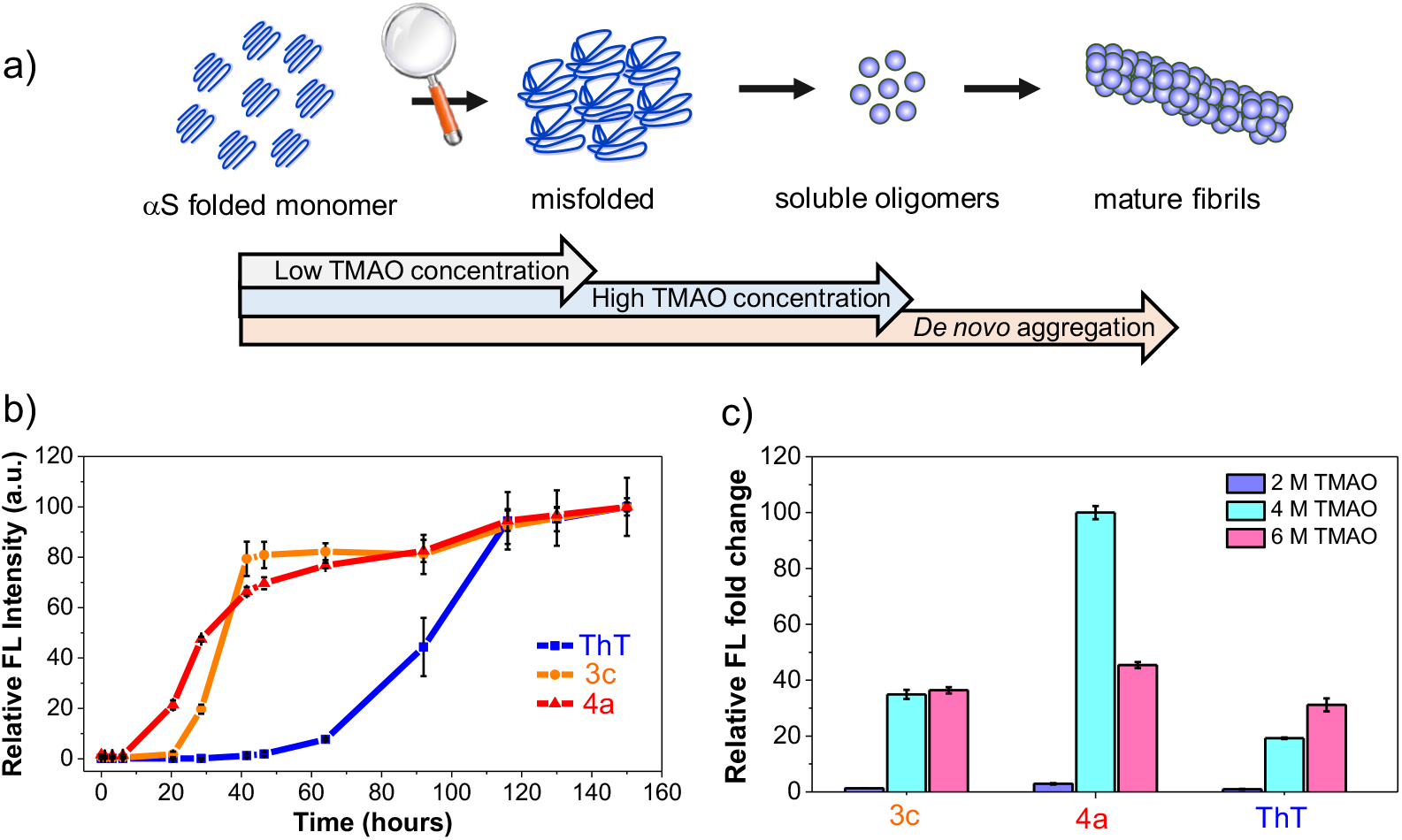
RBFs with extended linkages enable the detection of early misfolding and oligomers formation *in vitro*. (a) Schematic representation of the stepwise pathway for the formation of αS mature fibrils and αS misfolding or oligomer formation induced by varied TMAO concentration (b) Monitoring the αS aggregation over prolonged periods using **3c, 4a**, and ThT in PBS buffer at 37 °C. (c) Fluorescence fold change of RBFs **3c** and **4a** compared to the ThT with αS/TMAO solutions (50 μM of αS monomer and 2 M, 4 M, and 6 M of TMAO). The fluorescence fold change was determined relative to the fluorescence of the free probe in Tris buffer that contains 2 M, 4 M, and 6 M TMAO. Probes were used at a final concentration of 10 μM, mixed with 50 μM αS monomer. Ex/Em wavelengths: 463 nm/580 nm for **3c**, 484 nm/620 nm for **4a**, and 450 nm/482 nm for ThT. Error bars represent the SD of 3 measurements.

Next, we aimed to study whether these probes with different viscosity sensitivities can differentiate between early misfolding and oligomer formation. To this end, we exposed monomeric αS to varying concentrations of trimethyl-amine *N*-oxide (TMAO), a natural osmolyte well-known to induce compaction/misfolding at lower [TMAO] and oligomer formation at higher [TMAO] (**Figure 5a**).^43, 44^ Interestingly, **4a** exhibits some fluorescence turn-on at 2M TMAO concentrations, indicating misfolding, with 2.2 and 3.1-fold higher sensitivity than **3c** and ThT, respectively. At 4 M TMAO, **4a** shows the highest turn-on, corresponding to a mixture of misfolding and slight oligomer formation, with 2.9 and 5.2-fold higher sensitivity than **3c** and ThT, respectively. Finally, at 6 M TMAO with significant oligomer formation, **4a** shows similar sensitivity to **2a** and ThT, with just 1.2 and 1.5-fold greater turn-on, respectively (**Figure 5c**). Together, these results suggest that **4a** exhibits higher selectivity for early misfolding compared to later-stage oligomer formation.

Additionally, we observed distinct λ_em_ values for **4a** with varied TMAO concentrations (**Figure S30**). The initial red shift in λ_em_ from 604 to 606 nm when TMAO changes from 2 M to 4 M suggests an increase in polarity during misfolding, while the blue-shifted λ_em_ of 601 nm at 6 M TMAO indicates a decrease in polarity during oligomer formation. In contrast, **3c** is unable to monitor initial misfolding and only shows sensitivity to oligomer formation, with a decrease in λ_em_ to 566 nm to 560 nm at 4 M and 6 M TMAO, respectively (**Figure S31**). Together, these findings indicate that **4a** holds potential for understanding early stages of PD.

### RBFs with extended linkages facilitate fluorescence imaging of αS protein condensates *in vitro*

Studying biomolecular condensates typically requires protein labeling with genetically encoded tags (e.g., green fluorescent protein, HaloTag, SnapTag) or chemical conjugation of small organic dyes.^45, 46^ Given the sensitivity of our RBF probes to low viscosity aggregates, we wished to test their ability to detect unmodified αS in protein condensates.

αS is known to undergo LLPS through electrostatic interaction of its negatively charged C-terminal region with positively charged pLK^47^. We used a three-color imaging strategy to study the staining of αS (100 μM)/pLK (0.25 mg/mL) condensates by RBFs. The condensates contained 5% fluorescently labeled αS at position 94 with acridon-2-yl alanine (Acd, termed αS_94Acd_) and 5% pLK labeled with fluorescein isothiocyanate (FITC, termed pLK_FITC_). αS/pLK condensate formation was confirmed by confocal laser scanning microscopy (CLSM) of the Acd signal in the 430–480 nm channel for αS and the 500–540 nm channel for FITC-pLK (**Figure 6a**). Excitingly, we observed fluorescence signal of our RBFs within the condensates (570–620 nm channel for **3c** and the 630–680 nm channel for **4a)** (**Figure 6a**).

**Figure 6.**
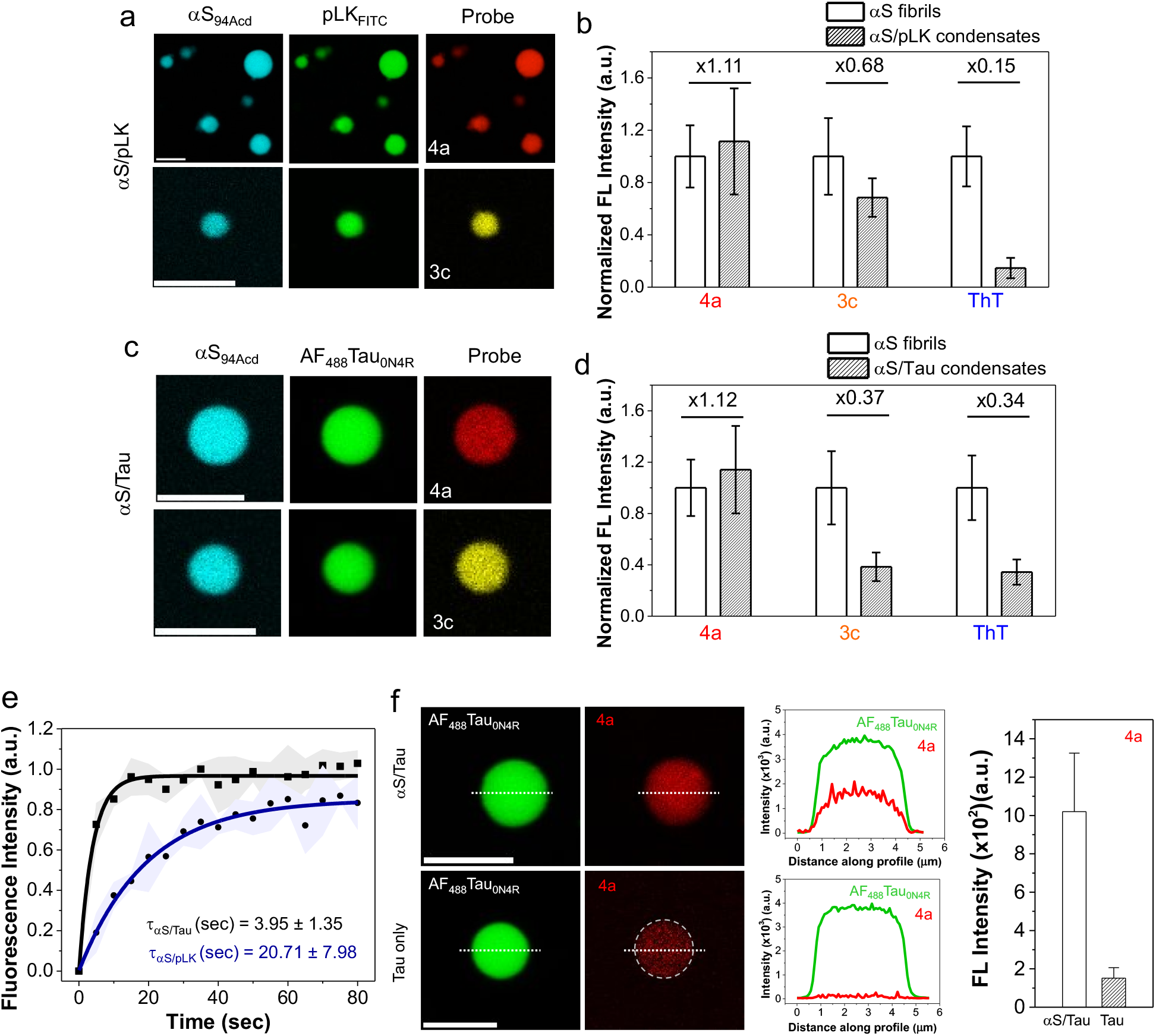
RBFs with extended linkages facilitate fluorescence imaging of αS protein condensates *in vitro*. (a) Staining of WT αS/pLK (100 μM/0.25 mg/mL) condensates by RBFs **3c** and **4a** in LLPS buffer (doped with 5% FITC-labeled pLK and 5% Acd-labeled αS) using CLSM imaging. (b) Analysis of fluorescence intensity of **3c, 4a**, and ThT with WT αS/pLK condensates compared to αS fibrils. Ratio of probe intensities (condensates to fibril) shown above the bars. (c) CLSM images of αS/Tau_0N4R_ condensates (80 μM of αS and 40 μM of tau in LLPS buffer) stained by RBFs **3c, 4a**, and ThT (doped with 5% AF_488_-labeled Tau_0N4R_ and 5% Acd-labeled αS). (d) Analysis of staining capacity of **3c, 4a**, and ThT with αS/Tau_0N4R_ condensates compared to αS fibrils. The ratio of intensities, condensate to fibril, shown for each probe. (e) FRAP profile showing the translational mobility of αS in αS/pLK and αS/Tau condensates (doped with 5% Acd-labeled αS). (f) Left: CLSM images of condensates of Tau_0N4R_ at a concentration of 40 μM with and without αS (80 μM) in LLPS buffer stained by **4a** (doped with 5% AF_488_-labeled Tau_0N4R_). Middle: Fluorescence intensity profile along the dotted white line showing higher fluorescence intensity for Tau_0N4R_ condensates with αS than without αS. Right: Analysis of fluorescence intensity of **4a** with Tau_0N4R_ condensates with and without αS. Error bars represent the standard deviation of >3 measurements. Ex: 488 nm and Em: 500-540 nm for AF_488_Tau_0N4R_, Ex: 561 nm and Em: 570-620 nm for **3c**, Ex: 561 nm and Em: 630-680 nm for **4a**, and Ex: 405 nm and Em: 430-480 nm for ThT and Acd-labeled αS. All experiments were performed in an LLPS buffer: 20 mM HEPES, 150 mM NaCl, and 1 mM TCEP at pH 7.4, with 15% PEG-8000, at room temperature. Scale bars are 5 μm.

Next, we analyzed the staining efficiency of **3c, 4a**, and ThT for αS/pLK LLPS condensates relative to αS fibrils (**Figure 6b** and **Figure S33**). RBF **4a** exhibited the highest staining efficiency (1.11-fold), followed by **3c** (0.68-fold), while ThT showed minimal staining (0.15-fold) in the LLPS condensates but excellent staining of fibrils, consistent with our *in vitro* aggregation kinetics and TMAO experiments. In addition, we observed that **4a** having low viscosity sensitivity showed the highest staining of condensates compared to fibrils.

Following these proof-of-concept experiments, we explored αS/tau protein condensates due to recent studies showing that these proteins are known to interact and mutually promote aggregation *in vitro* and *in vivo*, indicating potential crosstalk of these two proteins in AD, PD, and related dementias. To test the enrichment of RBFs for αS/tau protein condensates (80 μM/40 μM), we labeled the 0N/4R tau isoform of tau with Alexa Fluor 488 (referred to as AF_488_Tau_0N4R_) along with labeled αS_94Acd_.We employed a three-color imaging strategy to confirm the successful incorporation and fluorescence of the RBFs. As shown in **Figure 6c**, αS/tau condensate formation was confirmed using 5% labeled αS_94Acd_ and AF_488_Tau_0N4R_ via CLSM, with the 430–480 nm channel for αS and the 500–540 nm channel for tau (**Figure 6c**). RBF fluorescence was observed in the 570–620 nm channel for **3c** and the 630–680 nm channel for **4a** (**Figure 6c**). Moreover, we assessed the fluorescence of **3c, 4a** and ThT for αS/tau condensates in comparison to αS fibrils imaged under the same settings. As shown in **Figure 6d**, RBF **4a** exhibited the highest staining ability, with a 1.12-fold increase for αS/Tau condensates. In contrast, RBF **3c** demonstrated reduced staining capacity (0.37-fold), similar to ThT (0.34-fold) (**Figure S34**).

The difference in the staining efficacy of **3c** between αS/pLK and αS/Tau condensates was further investigated using fluorescence recovery after photobleaching (FRAP).^48^ FRAP revealed that αS/pLK condensates exhibit gel-like properties having diffusion time (τ_D_) around 20.71 ±7.98 s, while αS/tau condensates display liquid-like behavior with τ_D_ around 3.95 ±1.35 s (**Figure 6e** and **Figure S35**). Nevertheless, **4a** showed more diffuse staining of condensates than **3c**, which showed punctate staining. This difference likely results from the lower viscosity sensitivity of **4a**, allowing it to stain regions of varying viscosity, whereas the higher viscosity sensitivity of **3c** restricts its staining to the more viscous regions of the condensates (**Figure S36**).^49^ Together, these findings suggest that **4a** holds promise for studying liquid-like αS/tau condensates and understanding their molecular interplay and early co-aggregation mechanisms, whereas **3c** is more responsive to gel-like or solid-like structures, characteristic of later-stage aggregation species.

RBF **4a** has shown high *in vitro* selectivity for αS fibrils, including those amplified from patient-derived PDD samples (**Figure 4d** and **4f**). We sought to determine whether probe **4a** also exhibits selectivity for αS condensates over tau condensates. To test this, we prepared Tau_0N4R_ condensates (5% AF_488_-labeled Tau_0N4R_) with and without WT αS and assessed the incorporation of **4a** using a two-color imaging strategy *via* CLSM. Excitingly, **4a** exhibited minimal staining in tau-only condensates but showed strong fluorescence in αS/tau condensates, as observed in the 630–680 nm channel (**Figure 6f**). Analysis revealed a 3.8-fold higher fluorescence of **4a** into αS-containing tau condensates compared to tau-only condensates (**Figure 6f**). Collectively, these results demonstrate that probe **4a** can selectively stain αS condensates and holds promise for combined use with labeled tau to advance our understanding of the molecular mechanisms driving complex neurodegenerative diseases.

## Conclusion

In summary, we have shown that incorporating extended π-rich linkages and further changing the electron donation of aryl substituents on bimane chromophore provides a general approach to modulate the photophysical properties, polarity, viscosity sensitivity, and target selectivity of RBFs. These chemical modifications on RBFs result in a D-π-A framework that features 1) red-shifted photophysical properties with large Stokes shifts and 2) high control of the viscosity sensitivity through increased *E*_a_ between their planar fluorescent and twisted dark forms, as well as control of the polarity sensitivity through increased ICT.

RBFs with varied aryl substitution or linker lengths exhibit 1) tunable binding affinity and 2) fluorescent turn-on with distinct emission colors due to polarity shifts, supported by TD-DFT-calculated fluorescence energies, electrostatic potential maps, and observed dipole moments. Additionally, RBFs with an extended linker length not only showed high selectivity for αS fibrils *in vitro* over other amyloid fibrils such as tau and Aβ_1-42_ but also patient derived amplified fibrils of PDD. Mechanistic studies further reveal that the high selectivity of RBF **4a** stems from its specific binding to high-affinity sites, while **3c** binds nonspecifically to multiple sites.

RBF **4a**, with lower viscosity sensitivity, detects early misfolding and αS protein oligomers, whereas RBF **3c** is sensitive to later-stage species like oligomers. In contrast, the standard amyloid-binding dye ThT detects only mature fibrils. These findings align with TMAO-based fluorescence measurements, highlighting RBF **4a** as a superior probe for studying early stages of PD. Finally, we showed for the first time the use of small molecule based fluorescent probes for selective imaging of phase-separated αS. Chemical modifications provide RBFs **3c** and **4a** with tailored viscosity sensitivity, enabling successful staining of αS /pLK and αS/tau condensates. Notably, RBF **4a** showed selective imaging of αS condensates over tau condensates. Thus, **4a** complements other RBFs that are useful for selective imaging of αS fibrils, such as **Tg-52**, which has been shown to have specificity toward αS in PD over MSA.^16^

We anticipate that the environmental sensitivity and tunable properties of RBFs **3c** and **4a** may open avenues for various applications in biology and materials chemistry. As shown in **Figure S12**, RBF **4a** demonstrated higher sensitivity to low-polarity environments, making it a promising probe for studying the local polarity of cellular components or microenvironments in biological tissues. Additionally, RBFs **3c** and **4a**, with distinct sensitivity to high and low viscosities, respectively, can be combined to investigate the molecular mechanisms of αS aggregation, which progresses through a stepwise process, from low-viscosity misfolded oligomers to high-viscosity mature fibrils. As shown in **Figure S37**, bleed-through experiments with **3c** and **4a** confirmed no emission overlap with the Alexafluor 488 fluorophore channels when excited at 488 nm. This suggests that **3c** and **4a** can be effectively combined with the Alexafluor 488 probe to study proteins interacting with αS pathology, such as tau. In another example, these probes exhibit high fluorescence in the crystalline state, as shown in **Figure S16**, indicating their potential for use in organic materials. We will explore all of these applications in future studies.

## Notes

The authors declare no competing financial interest.

## ACKNOWLEDGMENT

This research was supported by the National Institutes of Health (NIH U19-NS110456 to E.J.P. and RF1-NS103873 to E.J.P.). Instruments supported by the NIH and NSF include NMR (NSF CHE-1827457), mass spectrometers (NIH RR-023444 and NIH S10-OD030460), and a computing cluster (NIH S10-OD023592). V.Y. and E.J.P. thank Virginia Lee and Nicholas Marotta for amplified fibrils from PDD patient samples and they thank Robert Mach and Ryann Perez respectively for contributing BF2846 and Ex6.

