## Supporting Information for "Strategic Modulation of Polarity and Viscosity Sensitivity of Bimane Molecular Rotor-Based Fluorophores for Imaging α-Synuclein"

**Table of Contents**


| | |
|---|---|
| General Information | S3 |
| Synthesis and Characterization of **3b, 3d, 3e,** and **4a** | S4-9 |
| General Photophysical Characterization Methods | S10 |
| Photophysical Properties of **3b, 3d, 3e,** and **4a** in ACN | S11 |
| Computational Studies (DFT and TD-DFT) | S12-13 |
| Fluorescent Quantum Yields and Solvatochromism of **3b, 3d, 3e,** and **4a** | S14-16 |
| Measurement of Spectral Properties at Different Polarities | S17-18 |
| Electrostatic Potential Maps and Polarity Analysis of **3a-e** and **4a** | S19 |
| Viscosity Sensitivity Measurements | S20-22 |
| Crystallization Induced Emission (CIE) Property of **3c** | S23 |
| Mechanistic Study of Rotational Energy Barrier $E_a$ | S24-25 |
| Fibril preparation and Fibril Binding Characterization of **3b, 3d, 3e,** and **4a** | S26-27 |
| Fluorescence Lifetime Measurements of Probes **3c**, **3e**, and **4a** with αS Fibrils | S28-29 |
| Tabulated Photophysical Properties of **3b-e** and **4a** with αS Fibrils | S29 |
| Fluorescent QY Measurements of **3c** and **4a** with αS Fibrils | S30 |
| Determination of the Dissociation Constant of Probes **3a-e** and **4a** with αS Fibrils | S31 |

| | |
|---|---|
| Displacement Assay of **3c** and **4a** | S32 |
| Probe Selectivity for $\alpha$S fibrils in HEK Cell Lysate | S33 |
| Fluorescence and Excitation Spectra of **4a** with Tau and $Ab_{1-42}$ Fibrils | S34 |
| Fractionation Assay of **3c** and **4a** | S35 |
| Dose-dependent fluorescence of **3c** and **4a** with $\alpha$S and $\alpha S_{1-100}$ fibrils | S36 |
| Time-dependent $\alpha$S aggregation monitored by RBFs **3c**, **4a**, and **ThT** | S37 |
| Experimental Procedure for the Fluorescent Measurements of **4a** with TMAO | S38 |
| Procedure for Fluorescence Measurements of **4a** with Patient Tissue Samples | S38 |
| Experimental Procedure for Liquid-Liquid Phase Separated Protein Condensates | S39 |
| Confocal Laser Scanning Microscopy (CLSM) Imaging of Protein Condensates | S39-40 |
| Procedure for Fluorescence Recovery After Photobleaching (FRAP) Measurements | S40 |
| CLSM imaging of $\alpha$S/pLK and $\alpha$S /Tau protein condensates *in vitro* | S42-43 |
| FRAP analysis of $\alpha$S dynamics within $\alpha$S /pLK and $\alpha$S /Tau | S44 |
| Diffusive and Punctate Staining of Condensates by **4a** and **3c** | S45 |
| Bleed-through Fluorescence Imaging of $\alpha$S/pLK and $\alpha$S/Tau Condensates | S46 |
| Cartesian Coordinates of the Optimized Geometries at Ground and Excited States | S47-53 |
| References | S54 |

### General Information

**Materials.** 4-(Methylamino)benzaldehyde, 4-(4-morpholinyl)benzaldehyde, and 4-(Dimethylamino)cinnamaldehyde was purchased from Millipore Sigma (St. Louis, MO, USA). 4-(azetidin-1-yl)benzaldehyde was purchased from Ambeed. Flash column chromatography was performed using Silicycle silica gel (40-63 μm (230- 400 mesh), 60 Å irregular pore diameter). Thin-layer chromatography (TLC) was performed on TLC Silica gel 60G F254 plates from Millipore Sigma. Reagents were purchased of the highest commercial quality and used without further purification, unless otherwise stated. The β-amyloid (1-42) peptide, human (Cat. No. RP10017) was purchased from Genscript (Piscataway, NJ). Ex6 and BF2846 were synthesized as previously described.[1]

**Instruments.** Nuclear magnetic resonance (NMR) spectra were obtained on a Bruker UNI-400 MHz or UNI-600 MHz instrument (Billerica, MA, USA) and are calibrated using peaks from residual protic solvent in deuterated solvent. The following abbreviations were used to denote multiplicities: s = singlet, d = doublet, t = triplet, q = quartet, m = multiplet, br = broad). Low-resolution mass spectra (LRMS) were obtained on a Waters Acquity Ultra Performance LC connected to a single quadrupole detector mass spectrometer (Waters Corp.; Milford, MA, USA). High-resolution mass spectra (HRMS) were obtained by Dr. Charles Ross III at the University of Pennsylvania's Mass Spectrometry Facility on a high-resolution electrospray ionization mass spectra (ESI-HRMS) using Waters LCT Premier XE liquid chromatograph/mass spectrometer. Absorbance readings for the DC assay were made on a Tecan M1000 plate reader (Mannedorf, Switzerland). UV-Vis absorption spectra were acquired on a Thermo Scientific Genesys 150 UV-Vis spectrometer (Waltham, MA, USA) using quartz cells with a 1 cm cell path length (Starna Cells, Inc 120 μL UV cells). Fluorescence spectra were acquired on a Tecan M1000 plate reader. Quantum yield (QY) measurements were performed using a Jasco FP-8300 Fluorimeter with ILF-835 integrating sphere attachment (Easton, MD, USA). Fluorescence lifetime measurements were made using a Photon Technology International (PTI) QuantaMaster™ 40 fluorescence spectrometer (Birmingham, NJ, USA).

**Horner-Wadsworth-Emmons (H-W-E) General procedure:** To a solution of phosphonate **1** (1 mmol) in dry DMF, the aryl aldehyde (1 mmol) and NaOMe (1.5 mmol) were added, and the resulting solution was stirred at RT for 40 h under argon atmosphere. The reaction progress was monitored by TLC, and the organic layer was extracted with DCM, washed with water (2 x 50 mL), dried using $Na_2SO_4$, and concentrated under vacuum to yield the crude product. Then, the product was purified by preparative thin layer chromatography (prepTLC) using 3% methanol in DCM as an eluent.

**Scheme S1.** Horner-Wadsworth-Emmons (H-W-E) reaction of bimane methyl phosphonate **1** and aryl aldehyde **2**[a].

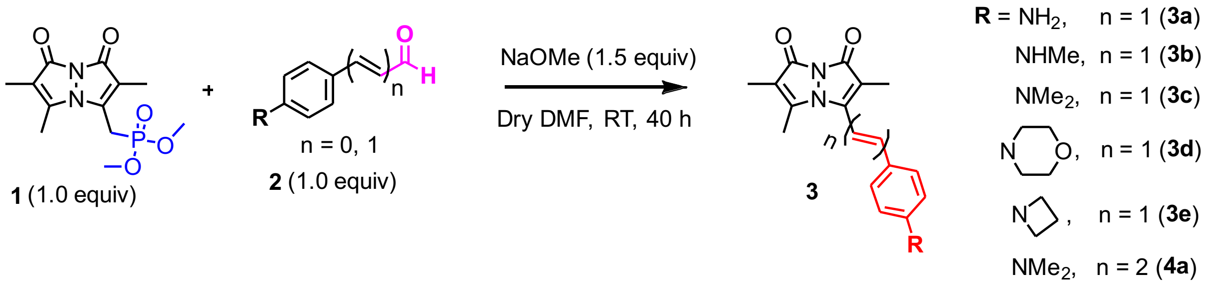


[a]Synthesis of compounds **3a** and **3c**, and their characterization by $^1$H, $^{13}$C NMR, HRMS were previously reported.[2]

**(*E*)-2,3,6-trimethyl-5-(4-(methylamino)styryl)-1*H*,7*H*-pyrazolo[1,2-a]pyrazole-1,7-dione (3b):**

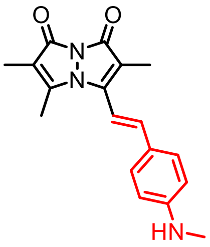

Following General Procedure with a solution of bimane methyl phosphonate **1** (50 mg, 0.167 mmol, 1 equiv) and 4-(methylamino)benzaldehyde (22.5 mg, 0.167 mmol, 1 equiv, purchased from Aldrich) at room temperature for 40 h afforded 20.4 mg (40 % isolated yield, orange solid) of the title compound after purification by prepTLC (3% methanol in dichloromethane, $R_f$ = 0.4). $^1$H NMR (600 MHz, DMSO) δ 7.49 (d, *J* = 8.7 Hz, 2H), 7.15 (d, *J* = 16.3 Hz, 1H), 6.82 (d, *J* = 16.3 Hz, 1H), 6.57 (d, *J* = 8.7 Hz, 2H), 6.32 (q, *J* = 5.1 Hz, 1H), 2.73 (d, *J* = 5.0 Hz, 3H), 2.34 (s, 3H), 1.91 (s, 3H), 1.73 (s, 3H). $^{13}$C NMR (151 MHz, DMSO) δ 161.6, 161.5, 152.2, 150.5, 149.9, 142.6, 129.9, 122.9, 112.2, 112.0, 109.8, 108.0, 29.8, 12.6, 8.6, 7.0. HRMS (ESI$^+$) calcd for $C_{18}H_{19}N_3O_2$ $[M + H]^+$, 310.1556; found: 310.1563.

**(*E*)-3-(4-(azetidin-1-yl)styryl)-2,5,6-trimethyl-1*H*,7H-pyrazolo[1,2-a]pyrazole-1,7-dione (3d):**

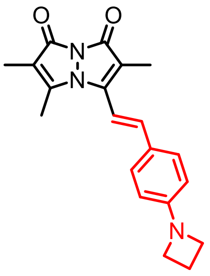

Following General Procedure with a solution of bimane methyl phosphonate **1** (60 mg, 0.2 mmol, 1 equiv) and 4-(azetidin-1-yl)benzaldehyde (32.2 mg, 0.2 mmol, 1 equiv, purchased from Aldrich) at room temperature for 40 h afforded 34.5 mg (51.5 % isolated yield, red solid) of the title compound **3d** after purification by prepTLC (3% methanol in dichloromethane, $R_f$ = 0.42). $^1$H NMR (600 MHz, DMSO) δ 7.56 (d, *J* = 8.6 Hz, 2H), 7.18 (d, *J* = 16.3 Hz, 1H), 6.89 (d, *J* = 16.3 Hz, 1H), 6.43 (d, *J* = 8.6 Hz, 2H), 3.90 (t, *J* = 7.3 Hz, 4H), 2.53 – 2.52 (m, 2H), 2.34 (s, 3H), 1.91 (s, 3H), 1.74 (s, 3H). $^{13}$C NMR (151 MHz, DMSO) δ 161.2, 161.0, 152.8, 150.1, 149.2, 141.8, 129.1, 123.6, 111.8, 110.8, 109.8, 108.6, 51.6, 40.1, 39.9, 39.8, 39.7, 39.5, 39.4, 39.2, 39.1, 16.2, 12.1, 8.1, 6.6. HRMS (ESI$^+$) calcd for $C_{20}H_{21}N_3O_2$ $[M + H]^+$, 336.1712; found: 336.1721.

**(*E*)-2,3,6-trimethyl-5-(4-morpholinostyryl)-1H,7H-pyrazolo[1,2-a]pyrazole-1,7-dione (3e):**

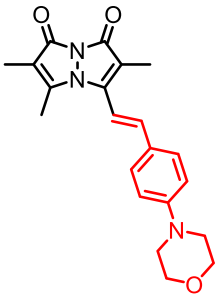


Following General Procedure with a solution of bimane methyl phosphonate **1** (100 mg, 0.33 mmol, 1 equiv) and 4-(4-morpholinyl)benzaldehyde (63.4 mg, 0.33 mmol, 1 equiv, purchased from Aldrich) in 2 mL dry DMF at room temperature for 40 h afforded 72.4 mg (59.5 % isolated yield, orange solid) of the title compound **3e** after purification by prepTLC (3% methanol in dichloromethane, $R_f$ = 0.44). $^1$H NMR (400 MHz, $CDCl_3$) δ 7.44 (d, J = 8.9 Hz, 2H), 7.03 (d, J = 16.3 Hz, 1H), 6.91 (d, J = 8.9 Hz, 2H), 6.60 (d, J = 16.3 Hz, 1H), 3.87 (t, J = 4.9 Hz, 4H), 3.26 (t, J = 4.9 Hz, 4H), 2.28 (s, 3H), 2.01 (s, 3H), 1.84 (s, 3H). $^{13}$C NMR (101 MHz, $CDCl_3$) δ 162.0, 152.6, 148.7, 148.5, 141.2, 128.9, 126.0, 115.1, 113.7, 112.6, 110.4, 66.8, 48.3, 12.7, 8.6, 7.2. HRMS (ESI$^+$) calcd for $C_{21}H_{23}N_3O_3$ $[M + H]^+$, 366.1818; found: 366.1811.

**3-((1*E*,3*E*)-4-(4-(dimethylamino)phenyl)buta-1,3-dien-1-yl)-2,5,6-trimethyl-1*H*,7*H*-pyrazolo[1,2-a]pyrazole-1,7-dione: (4a)**

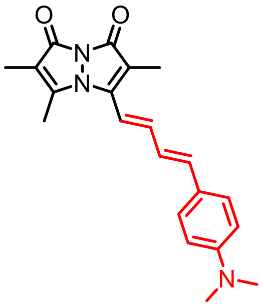


Following General Procedure with a solution of bimane methyl phosphonate **1** (100 mg, 0.33 mmol, 1 equiv) and 4-(dimethylamino)cinnamaldehyde (58.3 mg, 0.33 mmol, 1 equiv) in 2 mL of dry DMF at room temperature for 40 h afforded 51.6 mg (44.5 % isolated yield, red solid) of the title compound **4a** after purification by prepTLC (3% methanol in dichloromethane, $R_f$ = 0.48) to $^1$H NMR (400 MHz, $CDCl_3$) δ 7.37 (d, J = 9.0 Hz, 2H), 6.94 (dd, J = 15.1, 9.4 Hz, 1H), 6.83 – 6.74 (m, 2H), 6.68 (d, J = 9.0 Hz, 2H), 6.23 (d, J = 15.8 Hz, 1H), 3.02 (s, 6H), 2.30 (s, 3H), 2.01 (s, 3H), 1.84 (s, 3H). $^{13}$C NMR (151 MHz, DMSO) δ 161.0, 160.8, 150.7, 149.7, 148.3, 143.1, 139.5, 128.5, 123.7, 123.3, 114.1, 112.0, 111.6, 109.9, 39.9, 39.8, 39.6, 39.5, 39.4, 39.2, 39.1, 12.2, 8.2, 6.6. HRMS (ESI$^+$) calcd for $C_{21}H_{23}N_3O_2$ $[M + H]^+$, 350.1869; found: 350.1870.

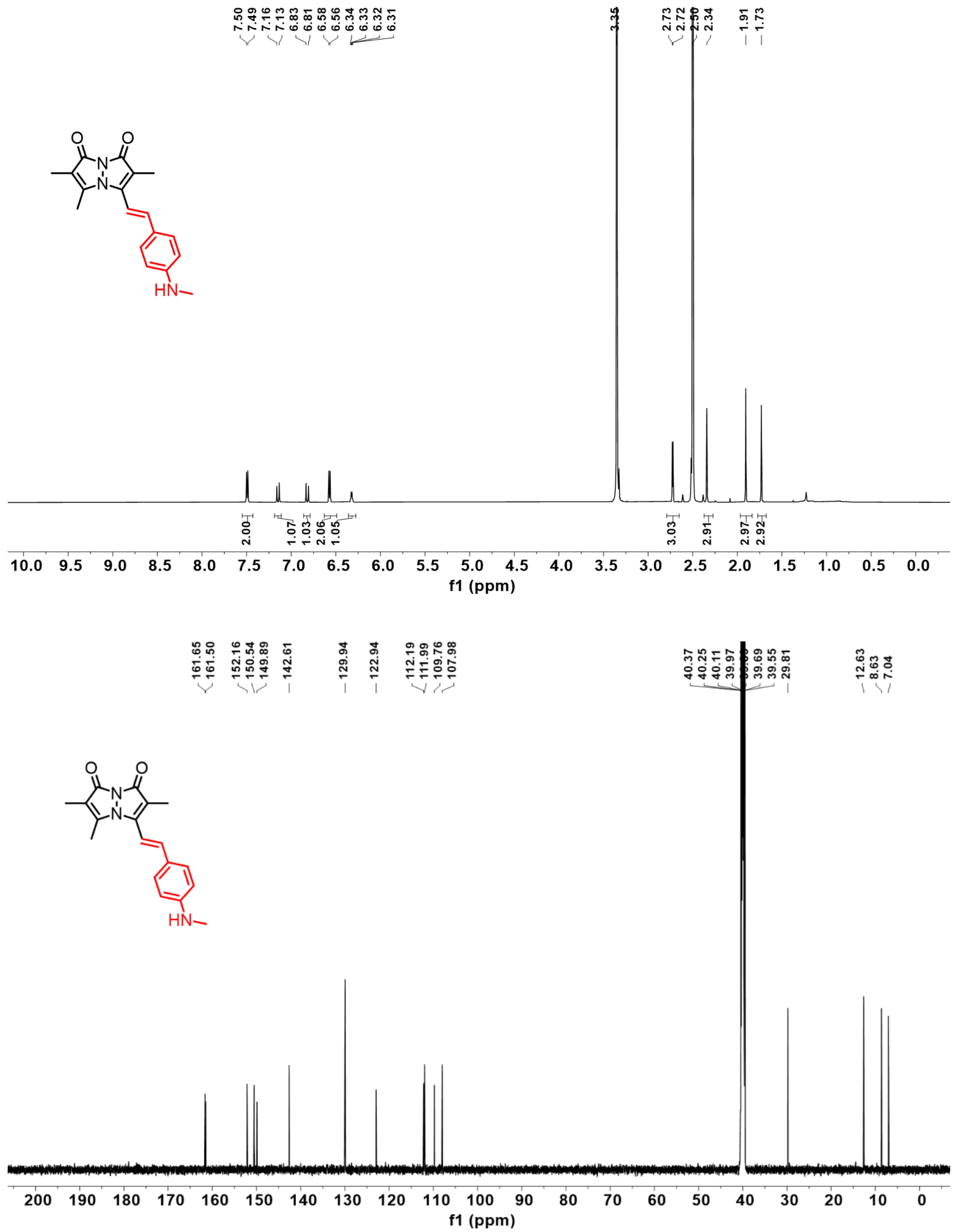


**Figure S1**. $^{1}H$ NMR and $^{13}C$ NMR spectrum of **3b** in DMSO-$d_6$.

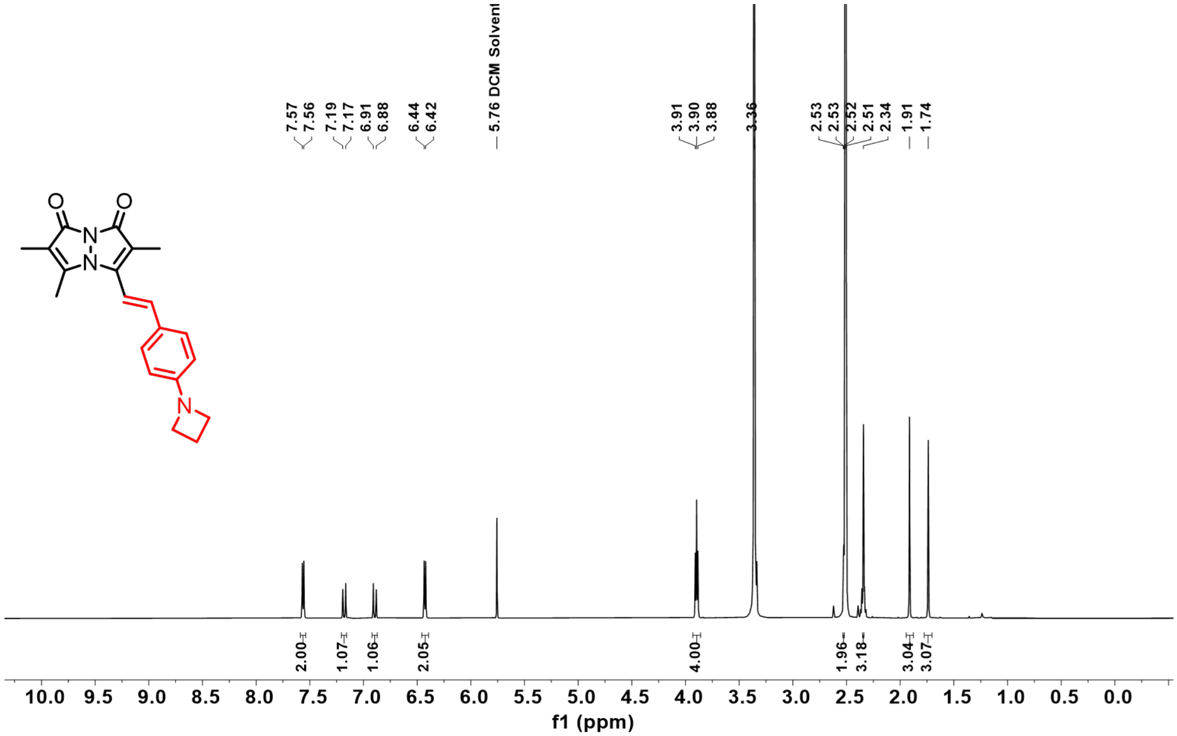

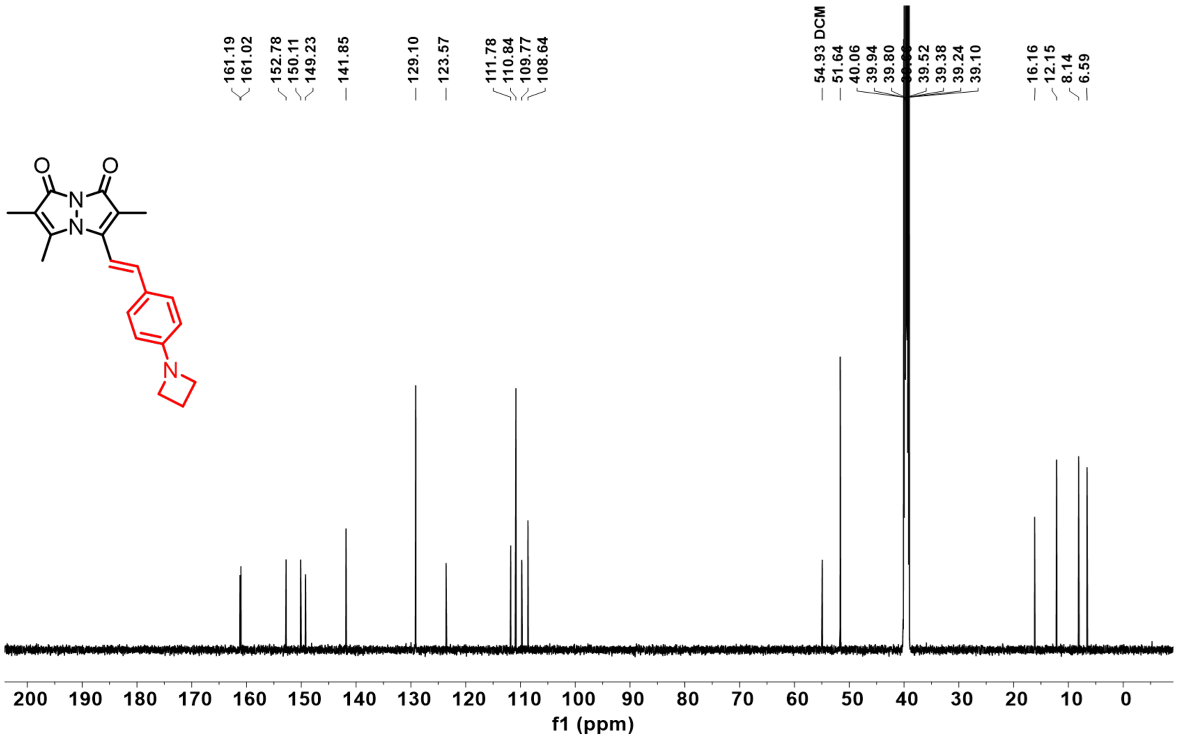

**Figure S2**. $^{1}H$ NMR and $^{13}C$ NMR spectrum of **3d** in DMSO-$d_6$.

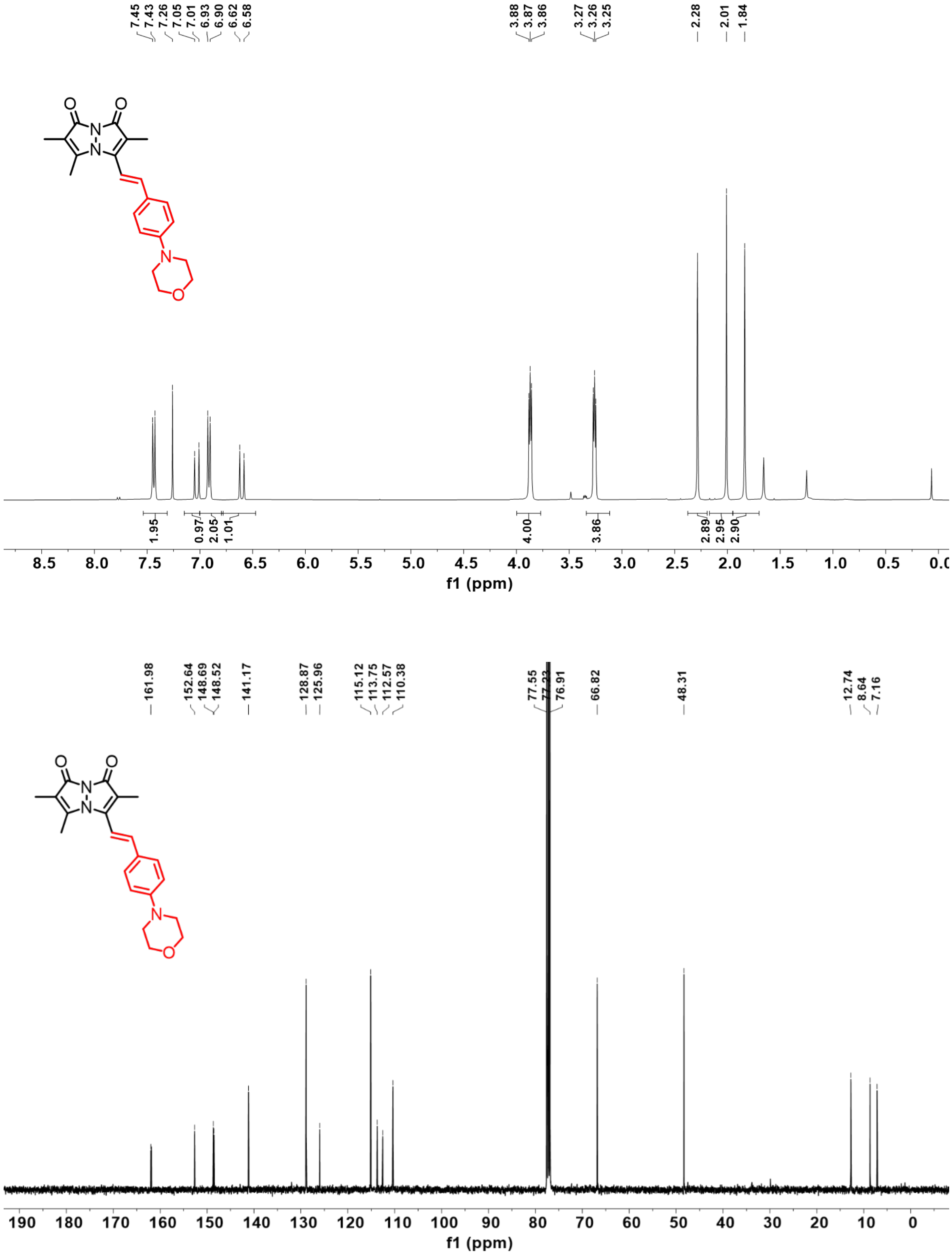


**Figure S3**. $^1$H NMR and $^{13}$C NMR spectrum of **3e** in $CDCl_3$.

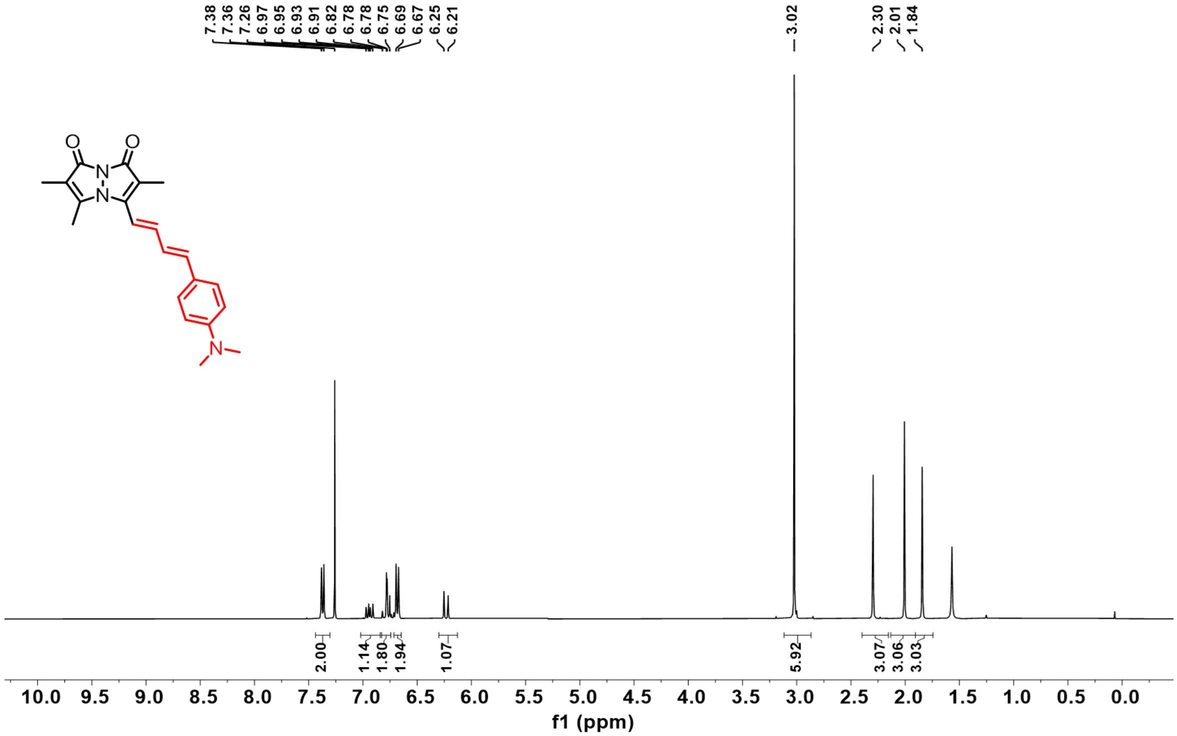


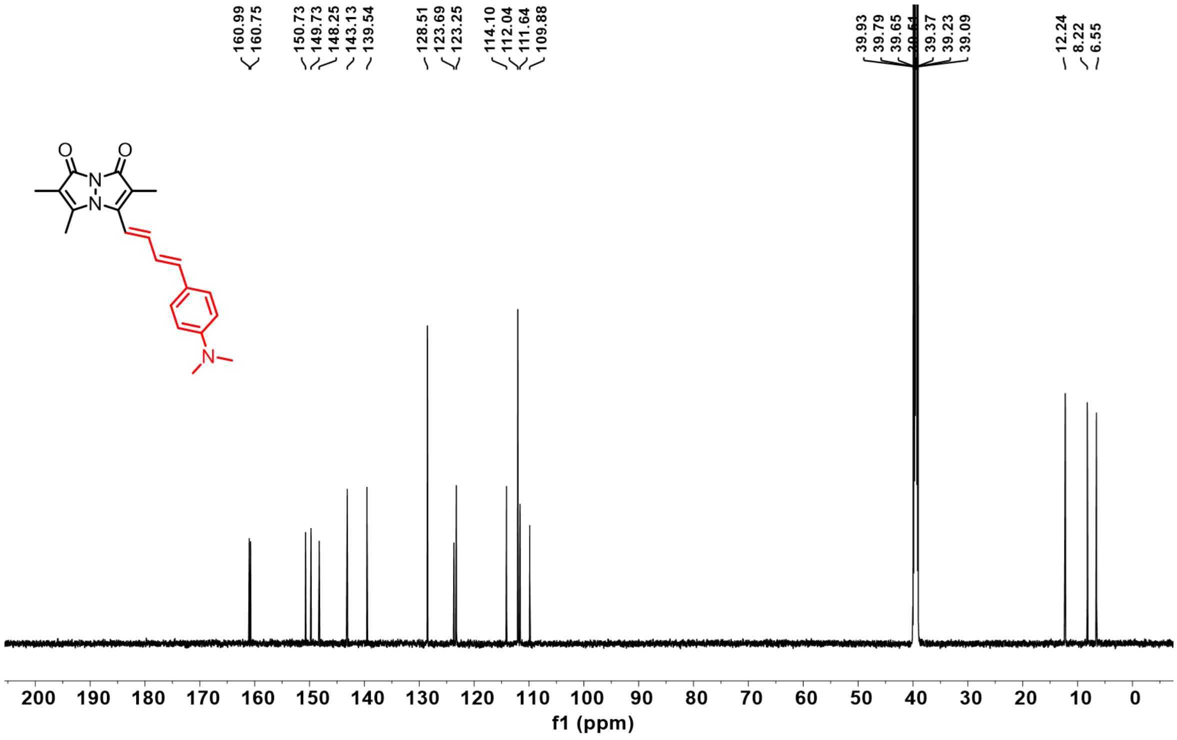


**Figure S4**. $^{1}$H NMR and $^{13}$C NMR spectrum of **4a** in DMSO-$d_6$.

### General Photophysical Characterization Methods

**Absorbance measurements.** Bimane derivatives **3a-e** and **4a** were dissolved in DMSO to make 10 mM starting stock solutions. Samples were individually diluted to the final concentration of 25 μM (500 μL) using 50:50 acetonitrile in phosphate buffered saline, pH 7.5 (ACN/PBS). The absorbance spectrum of each derivative (150 μL) was measured on a Thermo Scientific Genesys 150 UV-Vis spectrometer, using 50:50 ACN/PBS as a blank.

**Fluorescence measurements**. 10 mM stock solutions of **3a-e** and **4a** in DMSO were individually diluted to the final concentration of 25 μM (500 μL) using 50:50 ACN/PBS. The fluorescence spectrum of each derivative (150 μL) was measured on a Photon Technology International (PTI) QuantaMaster™ 40 fluorescence spectrometer by excitation at the maximum absorption wavelength for each compound.

**Molar absorptivity determination.** 10 mM stock solutions of **3a-e** and **4a** in DMSO were individually diluted to the final concentration of 25 μM (500 μL) using 50:50 ACN/PBS. The absorbance spectrum of each derivative (150 μL) was measured on a Thermo Scientific Genesys 150 UV-Vis spectrometer, using 50:50 ACN/PBS as a blank. Then, the molar absorptivity of each derivative was calculated by using the Beer–Lambert law for solutions, $A = \varepsilon lc$, where A = absorbance at maximum wavelength, l = optical path length in cm, c = concentration of the solution (25 μM), $\varepsilon$ = molar absorptivity.

**Solvatochromic measurements.** 10 mM stock solution of **3b**, **3d**, **3e**, and **4a** in DMSO were individually diluted into six different organic solvents (toluene, dichloromethane, ACN, dimethyl formamide, dimethyl sulfoxide, and ethanol) to a final concentration of 25 μM (1000 μL). The fluorescence spectrum of each derivative (150 μL) was measured on a PTI QuantaMaster™ 40 fluorescence spectrometer by excitation at the maximum absorption wavelength for each compound. The remaining 850 μL solution was used to obtain fluorescence images under a handheld 365 nm UV lamp.

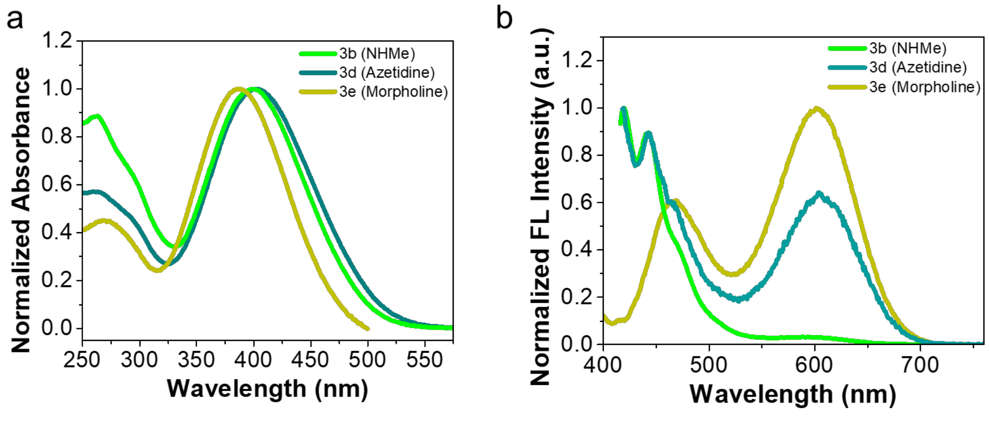


**Figure S5**. (a) Absorption spectra of **3b, 3d,** and **3e** and (b) Fluorescence spectra of **3b, 3d,** and **3e in** in ACN:PBS. Final probe concentration is 25 μM. Fluorescence spectra were measured at their maximum absorption wavelength.

#### Photophysical Properties of 3a-e, and 4a in ACN

**Table S1**. Photophysical properties of the bimane derivatives **3a-e,** and **4a** in ACN.

| RBF Dye | Substituent (Ph-R) | Absorbance $\lambda_{abs}$(nm)[a] | Emission $\lambda_{em}$ (nm)[b] | Molar Absorptivity (ε)($M^{-1}cm^{-1}$)[c] | Stokes shift (nm)[d] | Fluorescence quantum yield (%)[e] |
|---|---|---|---|---|---|---|
| **3a** | $NH_2$ | 378 | 540 | 32091 | 162 | 1.5 |
| **3b** | NHMe | 380 | 558 | 22861 | 178 | 2.0 |
| **3c** | $NMe_2$ | 402 | 568 | 24742 | 166 | 7.5 |
| **3d** | Azetidine | 385 | 576 | 21783 | 191 | 7.9 |
| **3e** | Morpholine | 378 | 565 | 28890 | 187 | 6.9 |
| **4a**[f] | $NMe_2$ | 410 | 618 | 28639 | 208 | 13.5 |

[a]Maximum absorption wavelength, [b]Maximum emission wavelength, [c]Molar absorption coefficients at maximum absorption wavelength, [d]Difference between maximum absorption wavelength and maximum emission wavelength, [e]Fluorescence quantum yield (error limit within ±5). [f]Probe with diene linker, Final probe concentration is 25 μM in ACN.

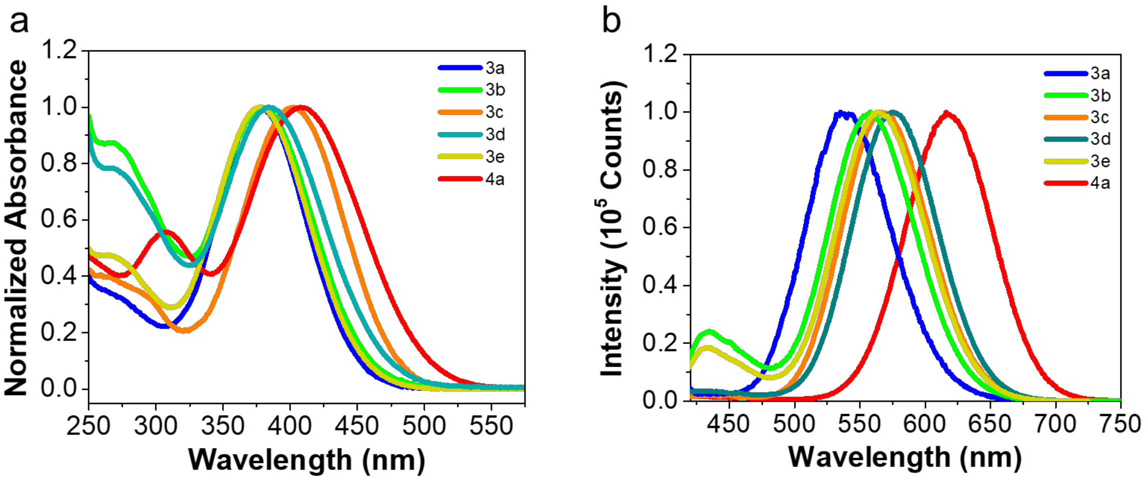


**Figure S6**. Comparing the photophysical properties of **3a-e** and **4a** in ACN solvent. (a) Absorption spectra of **3a-e** and **4a**. **(**b) Fluorescence spectra of **3a-e** and **4a**. Final probe concentration is 25 μM. Fluorescence spectra were measured at their maximum absorption wavelength.

### Computational Studies

**Methods.** All calculations were performed employing the APF-D density functional as implemented in the Gaussian16™ suite of programs with the 6-311+G (2d,p) basis set.[3] Geometry optimizations and energies were calculated for bimane derivatives **3a**, **3c**, **3d**, **3e** and **4a**. The HOMO-LUMO energy gaps between the ground state ($S_0$) and the first excited state ($S_1$) of the bimane derivatives were calculated and are shown below. All structures are ground-state minima according to the analysis of their vibrational frequencies, which showed no negative value.

**Table S2**. Calculation of the Energy Gaps of Bimane Derivatives at Ground State in Water ($H_2O$) Solvent.

| Compound | Absorbance Wavelength (nm)[a] | HOMO (eV) | LUMO (eV) | #LUMO-#HOMO Transition | LUMO-HOMO Δ(eV) |
|---|---|---|---|---|---|
| **3a** ($NH_2$) | 385 | -5.812 | -2.383 | 79-78 | 3.429 |
| **3c** ($NMe_2$) | 418 | -5.528 | -2.356 | 87-86 | 3.172 |
| **3d** (Azetidine) | 403 | - 5.575 | - 2.366 | 90-89 | 3.209 |
| **3e** (Morpholine) | 386 | -5.725 | -2.379 | 98-97 | 3.346 |
| **4a** ($NMe_2$)[b] | 436 | -5.380 | -2.508 | 94-93 | 2.872 |

[a]Absorbance spectra recorded in ACN:PBS buffer (50:50 v/v), [b]Probe with diene linker.

**Table S3**. Calculation of the Emission Energy of Bimane Derivatives in Water ($H_2O$) Solvent.

| Compound | Emission Wavelength (nm)[a] | Emission Energy (eV) |
|---|---|---|
| **3a** ($NH_2$) | 583 | 2.40 |
| **3c** ($NMe_2$) | 604 | 2.32 |
| **3d** (Azetidine) | 605 | 2.28 |
| **4a** ($NMe_2$)[b] | 640 | 2.05 |

[a]Emission spectra recorded in ACN:PBS buffer (50:50 v/v), [b]Probe with diene linker.

**HOMO-LUMO gap calculations:** We have analyzed the correlation between the photophysical properties of bimane derivatives and the electronic properties of the substituents to understand the mechanism of fluorescence tuning of the bimane scaffold. It is generally accepted that the absorption wavelength ($\lambda_{abs}$) and emission wavelength ($\lambda_{em}$) correlate well with the energy gap between HOMO and LUMO as determined by simple electronic structure calculations. In this study, geometry optimizations in the ground state were performed using density functional theory (DFT), and excited-state optimizations were conducted with time-dependent DFT (TD-DFT), both employing the APF-D density functional as implemented in the Gaussian16™ suite of programs with the 6-311+G (2d,p) basis set. RBFs **3a** ($NH_2$, n = 1, 385 nm), **3c (**$NMe_2$, n = 1, 418 nm), and **4a** ($NMe_2$, n = 2, 436 nm) with electron donating groups at the *para*-position, showed a decrease in HOMO-LUMO gap that matched well with red-shifts of the UV-Vis absorbance (Table S2 and Figure S5a).

Again, the calculated emission energy values for **3a**, **3c**, and **4a** matched well with experimentally measured red shifted emission wavelengths (Table S3 and Figure S5b). For **3a**, **3c**, **3d**, and **4a**, the electrons move from a HOMO centered on the dimethylaminophenylene group to a LUMO centered on the bimane group.

Interestingly, for RBF **3d** (Azetidine, n =1, 403 nm), we have observed blue-shifted absorption compared to the **3c (**$NMe_2$, n = 1, 418 nm) that correlates well with the increased energy gap between HOMO and LUMO around 3.209 eV compared to the 3.172 for **3c**. In contrast, we observed that the red-shifted emission for **3d** around 605 nm compared to 604 nm for **3c** correlates well with the decreased emission energy of 2.28 eV for **3d** compared to 2.32 eV for **3c** (Table S3 and 4 and Figure S5a and b).

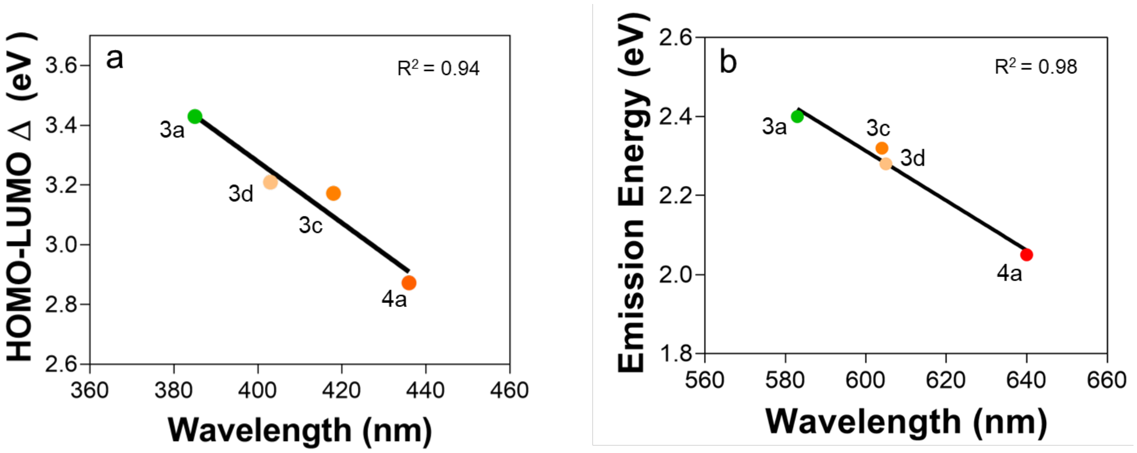


**Figure S7**. (a) Correlation between HOMO-LUMO energy gap (eV) compared to experimentally measured UV-Vis absorption maximum (nm). (b) Correlation between calculated emission energy (eV) compared to experimentally measured emission maximum (nm). Maximum UV-Vis absorption and emission wavelengths were obtained in ACN/PBS buffer (50:50 v/v). Symbol circle shows the experimental data, and line shows the linear fitting data.

**Fluorescent Quantum Yield Measurements in ACN and ACN/PBS buffer**

**Fluorescent quantum yield (QY) measurements**. For each QY measurement, the incident excitation light spectrum was collected with 1 mL of solvent. After measuring the incident light intensity, probes **3b, 3d, 3e,** and **4a** were added to the solvent from a concentrated stock solution to a final concentration of 500 μM and the new spectrum (fluorescence intensity and new incident light intensity) was collected. Using the JASCO Quantum Yield Software, the dye QY was calculated by dividing the dye fluorescence intensity by the difference in incident light intensity in the presence and absence of dye. The minimum excitation wavelength for a full excitation incident spectrum using this setup is 360 nm. QY values are reported in the main text Table 1.

**General Procedure for the Measurement of Spectral Properties at Different Polarities**

10 mM stock solutions of **3a-e** and **4a** in DMSO were individually diluted to the final concentration of 25 μM (1000 μL, 0.25% DMSO) in toluene, dichloromethane (DCM), acetonitrile (ACN), and acetonitrile/water mixtures (50:50 v/v). The fluorescence spectrum of each solution (150 μL) was measured on a Photon Technology International (PTI) QuantaMaster™ 40 fluorescence spectrometer by excitation at the maximum absorption wavelength for each compound. Then, the dielectric constant $\varepsilon$ of the solvents was used as an index of its polarity. Here, we used the dielectric constants 2.38 for toluene, 8.93 for DCM, 37.5 for ACN and the ACN/$H_2O$ mixture dielectric constant $\varepsilon_{mix}$ can be calculated using Eq. 1.[4]

$$\varepsilon_{mix} = \phi_1 \,.\, \varepsilon_1 + \phi_2 \,.\, \varepsilon_2 \qquad \text{———————} \qquad (1)$$

Where $\varepsilon$ stands for the dielectric constant, with the subscripts 1 and 2 referring to acetonitrile and $H_2O$, respectively. Factor $\phi$ stands for the weight fraction of each solvent. To calculate $\phi$, the values 0.786 and 1.00 have been used for the density of ACN and $H_2O$, respectively. Dielectric constants 37.5 (ACN) and 80.1 ($H_2O$) were used to estimate the dielectric constant of the mixtures.

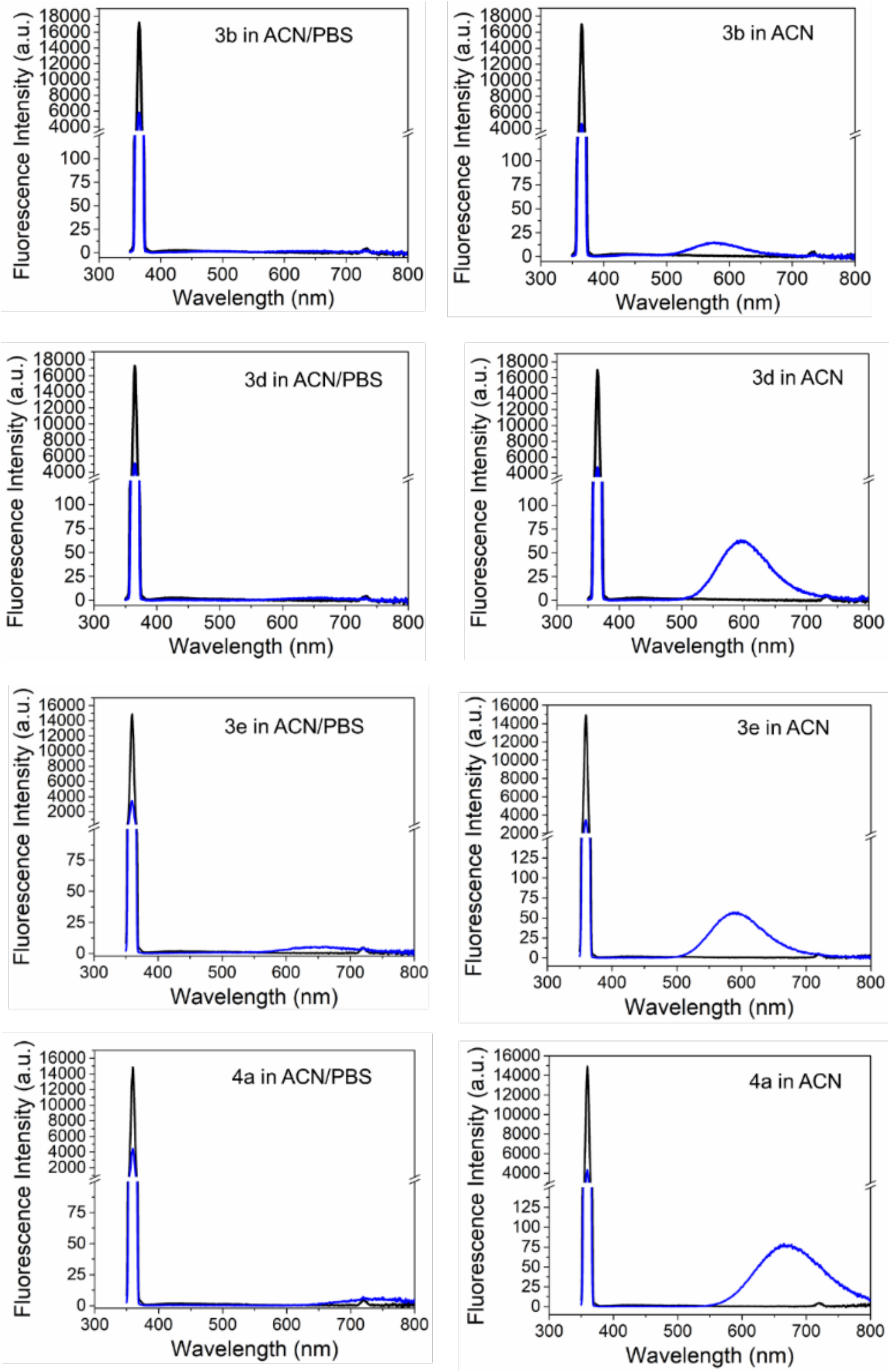


**Figure S8**. Representative QY acquisitions of **3b**, **3d**, **3e**, and **4a** in ACN/PBS buffer (50:50 v/v) and ACN solvents. Excitation at 360 nm and spectral collection from 350–800 nm. Final concentration of the probe is 100 μM. Black line indicates incident light, and blue line indicates sample spectrum.

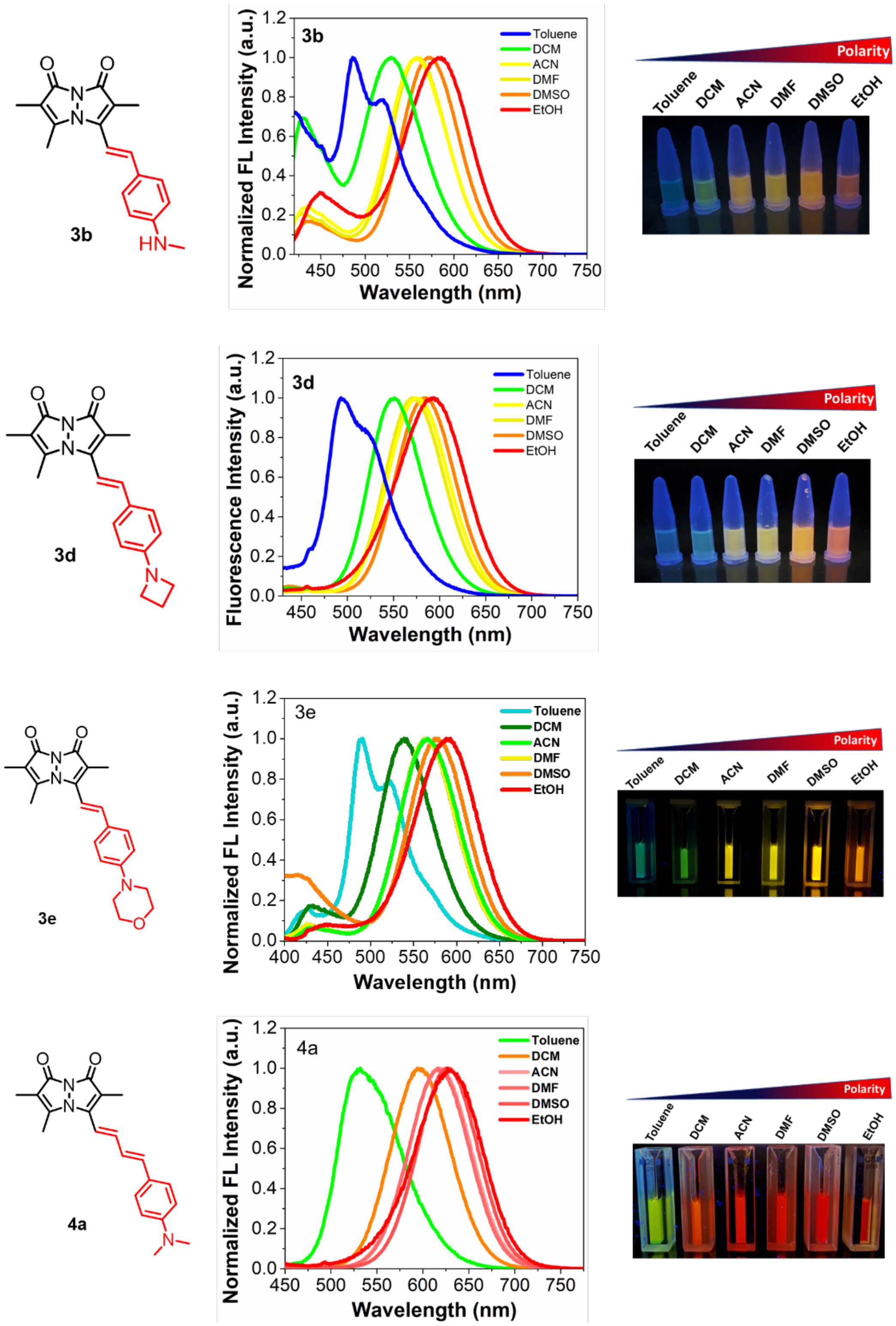


**Figure S9**. Solvatochromism of **3b**, **3d**, **3e**, and **4a**. Left: Normalized fluorescence spectra in different solvents. Probe concentration 25 μM. Right: Fluorescence image taken under handheld UV lamp at 365 nm.

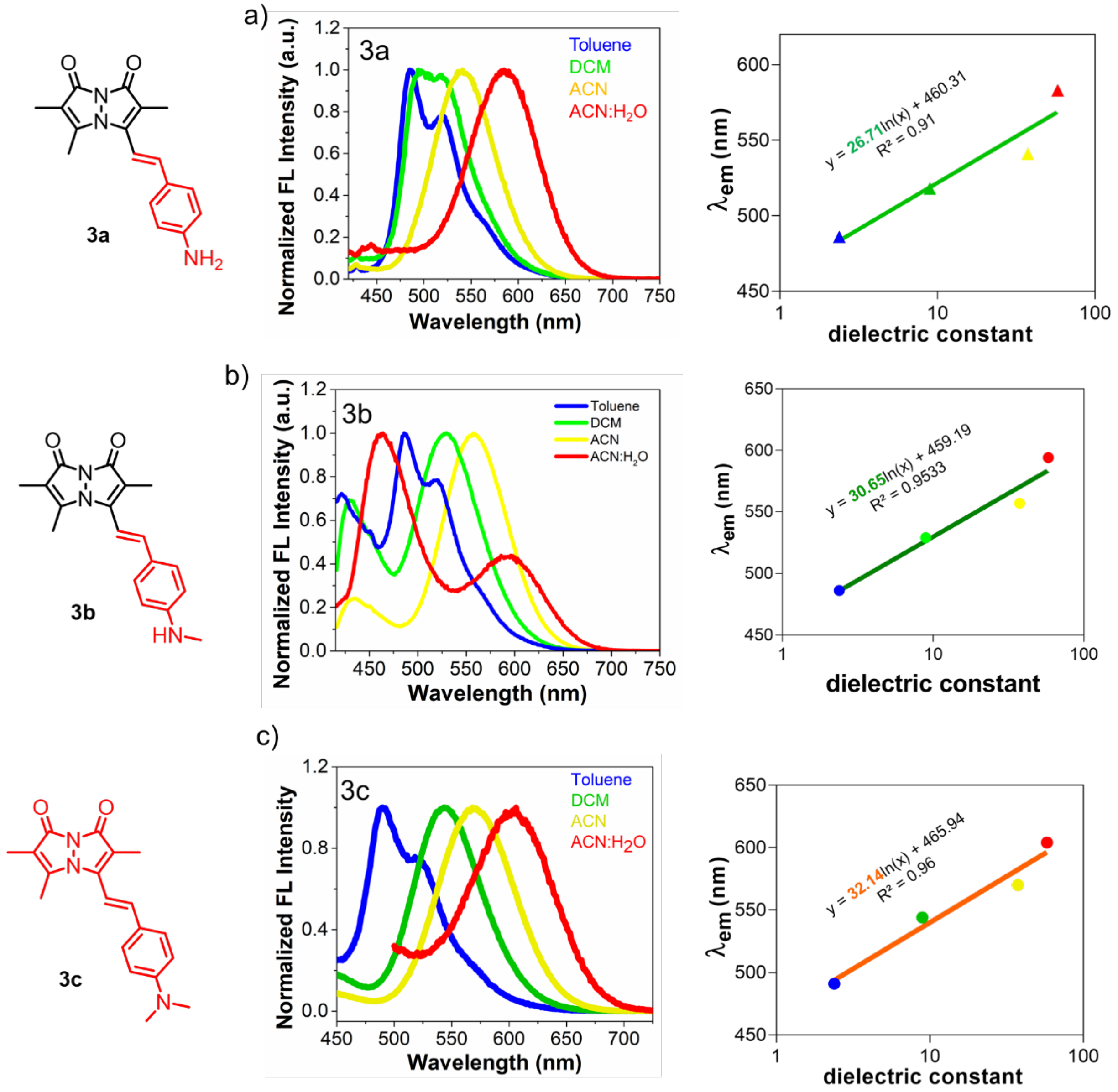


**Figure S10**. Control of polarity sensitivity by varying the substitution or linker length. (a) Solvatochromism of RBFs in solvents with different polarity (left) and Quantitative polarity sensitivity of RBFs (right). a) **3a**, b) **3b**, c) **3c**, d) **3d**, e) **3e,** and f) **4a**. Final concentration of probe is 25 μM in indicated solvents. Ex: 385 nm for **3a**, 399 nm for **3b**, 418 nm for **3c**, 403 nm for **3d**, 386 nm for **3e** and 436 for **4a.**

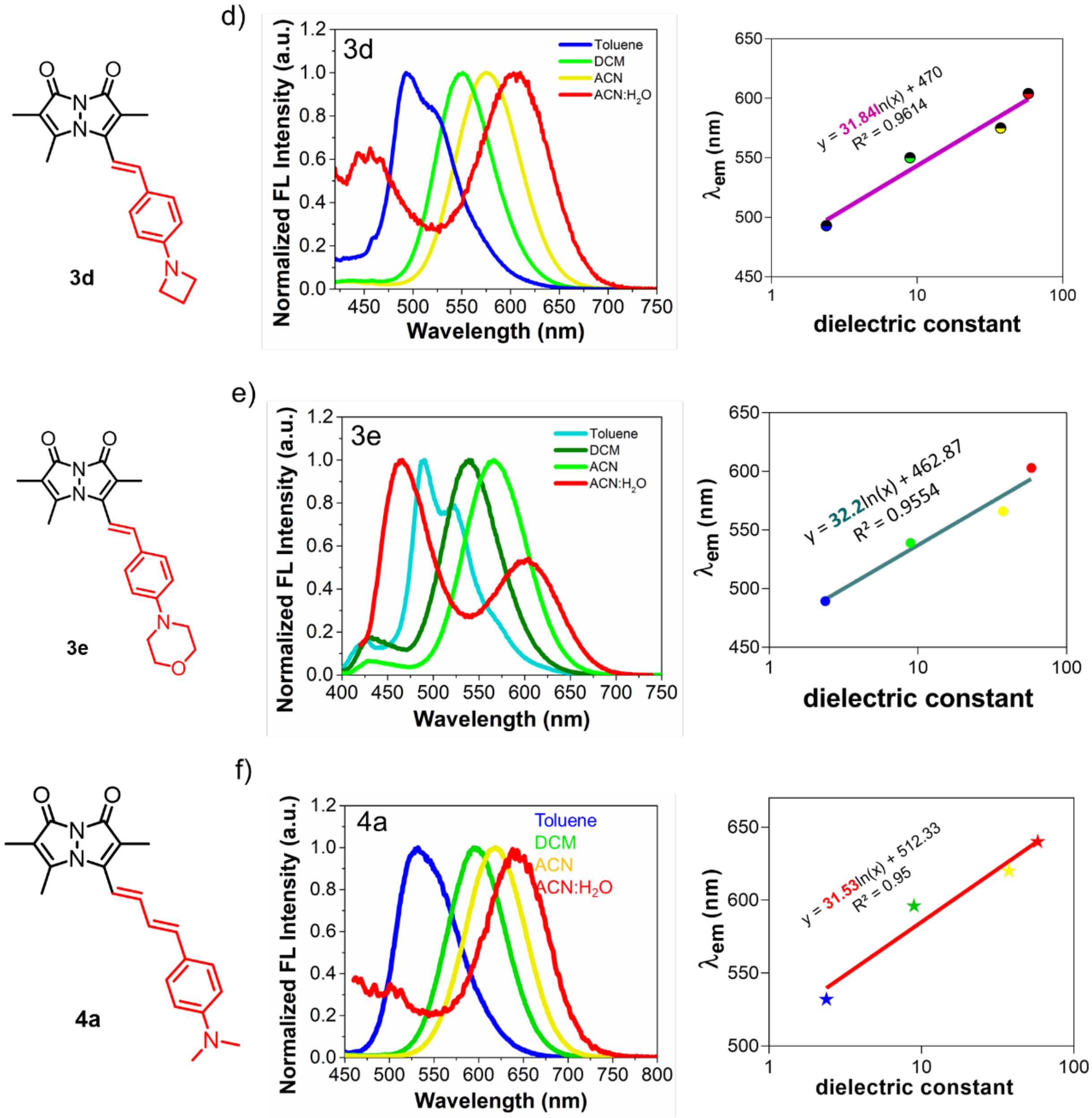


**Figure S10 (cont'd)**. Control of polarity sensitivity by varying the substitution or linker length. (a) Solvatochromism of RBFs in solvents with different polarity (left) and Quantitative polarity sensitivity of RBFs (right). a) **3a**, b) **3b**, c) **3c**, d) **3d**, e) **3e,** and f) **4a**. Final concentration of probe is 25 μM in indicated solvents. Ex: 385 nm for **3a**, 399 nm for **3b**, 418 nm for **3c**, 403 nm for **3d**, 386 nm for **3e** and 436 for **4a.**

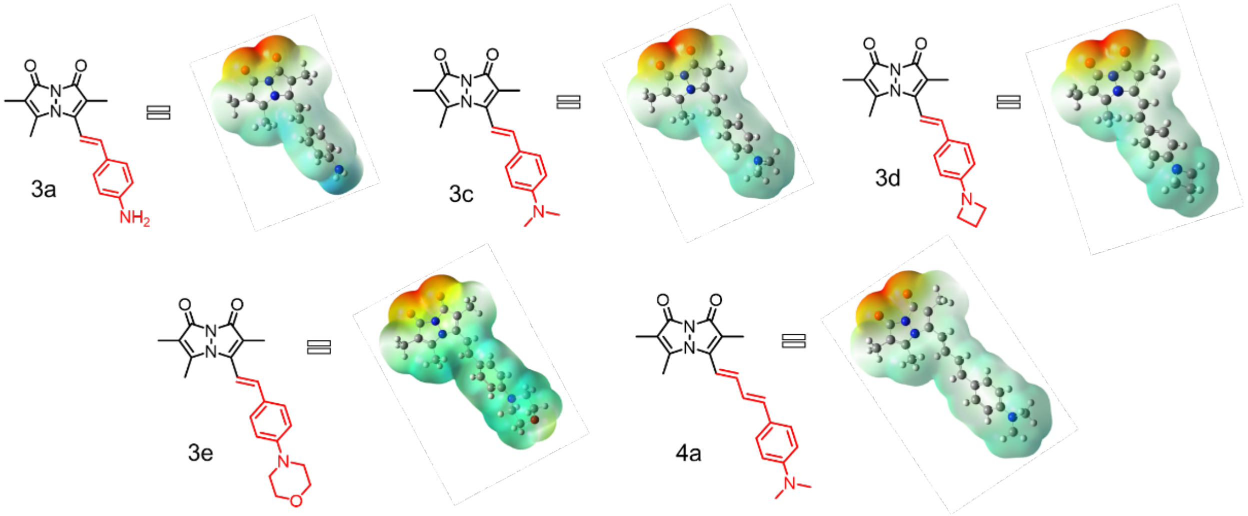


**Figure S11**. Electrostatic potential maps based on the electron density of **3a**, **3c**, **3d**, **3e**, and **4a** in water further explains the polarity sensitivity, with red color indicating negative electrostatic potential regions with the highest electron density and light blue color indicating the most positive electrostatic potential regions with the lowest electron density.

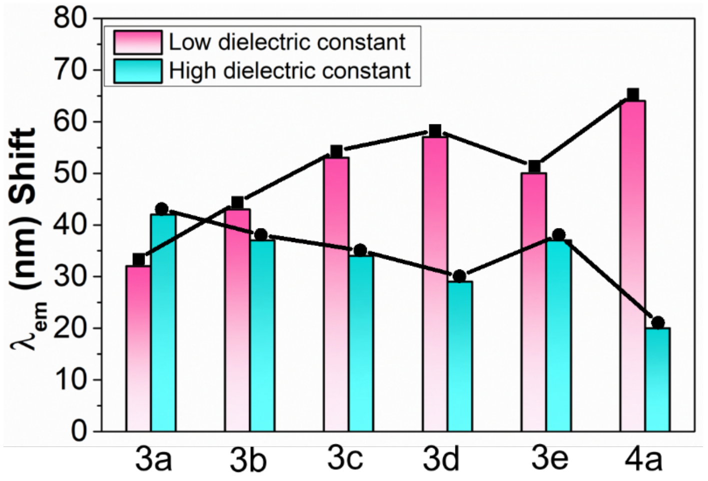


**Figure S12**. Fluorescence emission maximum shift of compounds **3a–e** and **4a** in the presence of solvents with low and high dielectric constants. The pink bars represent the emission wavelength ($\lambda_{em}$) shift (nm) between solvents with low dielectric constants (2.38 and 8.93). The cyan bars represent the $\lambda_{em}$ shift (nm) between solvents with high dielectric constants (37.5 and 58).

### Viscosity sensitivity measurements

To reduce the impact of solvent polarity on fluorescence emission intensity, a glycerol-ethylene glycol mixture was selected for viscosity sensitivity measurements due to their closely matched dielectric constants (glycerol: 46.5, ethylene glycol: 37.0). RBFs **3a-e** and **4a** are diluted from 10 mM DMSO stock into a series of ethylene glycol and glycerol mixtures (0%, 20%, 40%, 60%, 80%, and 90% glycerol in ethylene glycol) at 25 °C with a final concentration 25 μM. Viscosity of the two-element mixtures was calculated according to the previously reported method at 25 °C as shown is in **Table S4**.[4] Fluorescence intensity measurements were made using a Tecan Spark Fluorescence Plate Reader in Greiner 384 Flat Transparent plates. Excitation and emission wavelength were 385 nm/583 nm for **3a**, 399 nm/591 for **3b**, 418 nm/604 nm for **3c**, 403 nm/605 nm for **3d**, 386 nm/603 nm for **3e,** and 436 nm/640 nm for **4a**.

The viscosity-dependent curve was plotted using solvent viscosity as X-axis and emission intensity as Y-axis, and the viscosity dependence parameters $\chi_1$ and $\chi_2$ was determined by non-linear polynomial regression of the second order as shown in **Eq.2**.

$$I = c + \chi_1\eta + \chi_2\eta^2 \qquad (2)$$

Where "I" is the emission peak intensity, "$\eta$" is the solvent viscosity, "c" is the intercept of the curve, and "$\chi_1$ and $\chi_2$" represent the viscosity sensitivity values, where $\chi_1$ represents the linear change in intensity with viscosity, and $\chi_2$ represents the curvature or non-linearity in the relationship. Error bar: standard error (n =3).

**Table S4.** Solvent composition and viscosity of ethylene glycol and glycerol mixture.

| Composition Glycerol *vol*% | 0% glycerol | 20% glycerol | 40% glycerol | 60% glycerol | 80% glycerol | 90% glycerol |
|---|---|---|---|---|---|---|
| Viscosity (cP) at 25 ˚C | 16 | 219 | 412 | 594 | 768 | 851 |

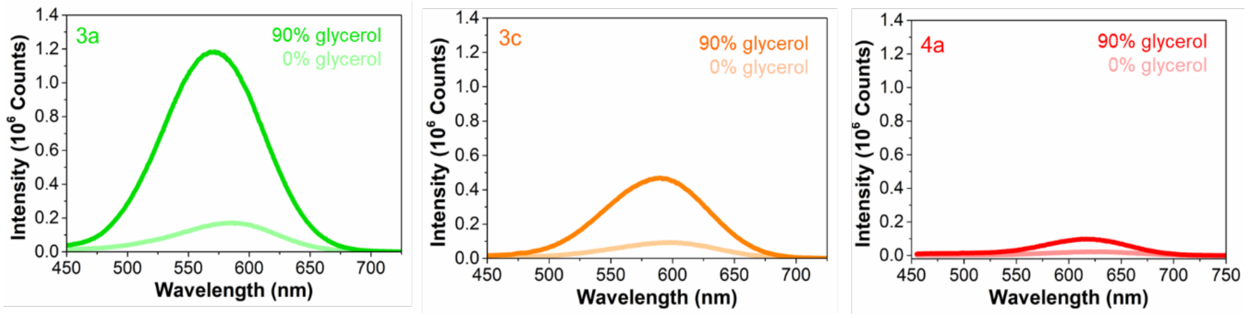


**Figure S13**. Viscosity sensitivity of RBFs is significantly and systematically affected by either change in aryl substitution or the introduction of extended linkages between donor and acceptor. Fluorescence spectra of RBFs **3a** (left), **3c** (middle), and **4a** (right) in an ethylene glycol mixture containing 0% and 90% glycerol. Ex/Em: 385 nm for **3a**, 418 nm for **3c**, and 436 nm for **4a**. Final probe concentration is 25 μM.

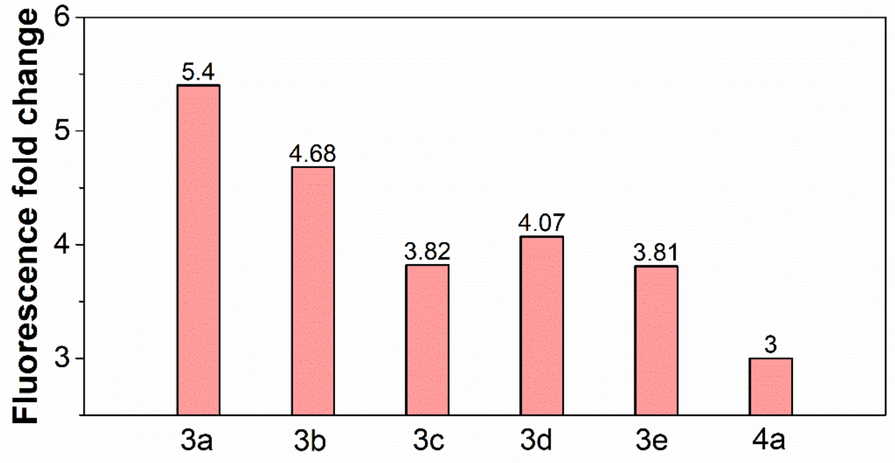


**Figure S14**. Fluorescence fold change of RBFs **3a-e** and **4a** in an ethylene glycol mixture containing 0% and 90% glycerol. Ex/Em: 385 nm/583 nm for **3a**, 399 nm/591 for **3b**, 418 nm/604 nm for **3c**, 403 nm/605 nm for **3d**, 386 nm/603 nm for **3e,** and 436 nm/640 nm for **4a**. Final probe concentration is 25 μM.

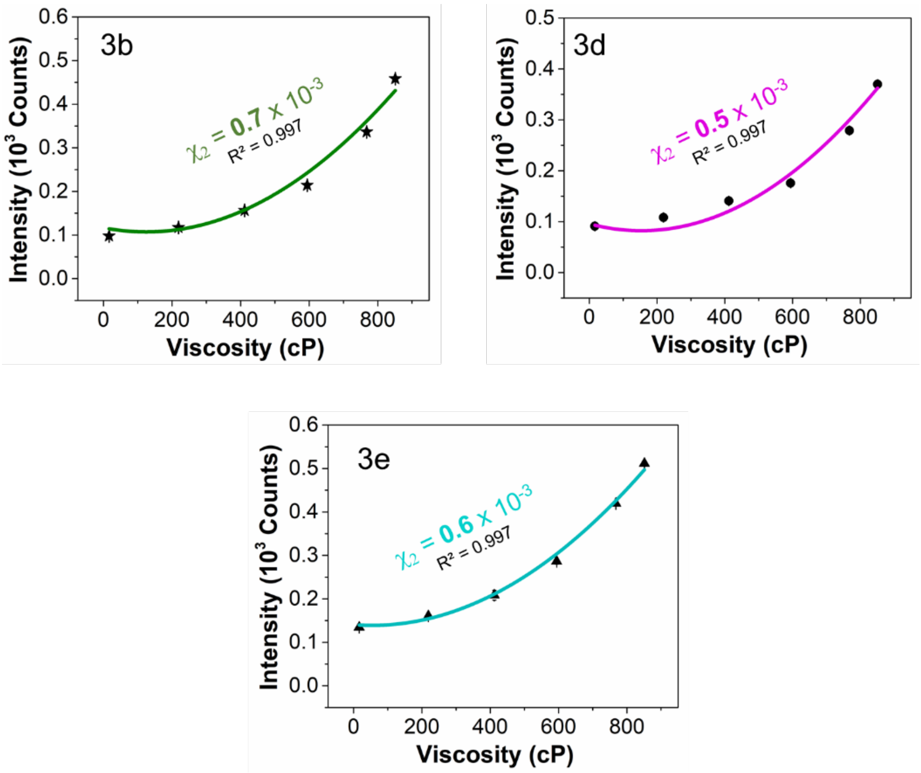


**Figure S15**. Quantitative viscosity sensitivity of **3b**, **3d,** and **3e**, measured in an ethylene glycol and glycerol mixtures with varied viscosity. Ex/Em: 399 nm/591 for **3b**, 403 nm/605 nm for **3d**, 386 nm/603 nm for **3e.** Final probe concentration is 25 μM.

**Table S5.** Fitting parameters of **3a-e** and **4a** by second-order polynomial regression.

| **RBF** | **non-linear change ($\chi_2$)** | **linear change ($\chi_1$)** | **Intercept (C)** | $R^2$ |
|---|---|---|---|---|
| **3a** ($NH_2$) | $1.0 \times 10^{-3}$ | 0.4975 | 287.6 | 0.993 |
| **3b** (NHMe) | $0.7 \times 10^{-3}$ | -0.2008 | 111.9 | 0.980 |
| **3c** ($NMe_2$) | $0.6 \times 10^{-3}$ | 0.00009 | 173 | 0.998 |
| **3d** (Azetidine) | $0.5 \times 10^{-3}$ | -0.155 | 102.9 | 0.976 |
| **3e** (Morpholine) | $0.6 \times 10^{-3}$ | -0.1058 | 142.5 | 0.996 |
| **4a** ($NMe_2$)[a] | $0.2 \times 10^{-3}$ | 0.0007 | 82.7 | 0.996 |

[a]Probe with diene linker

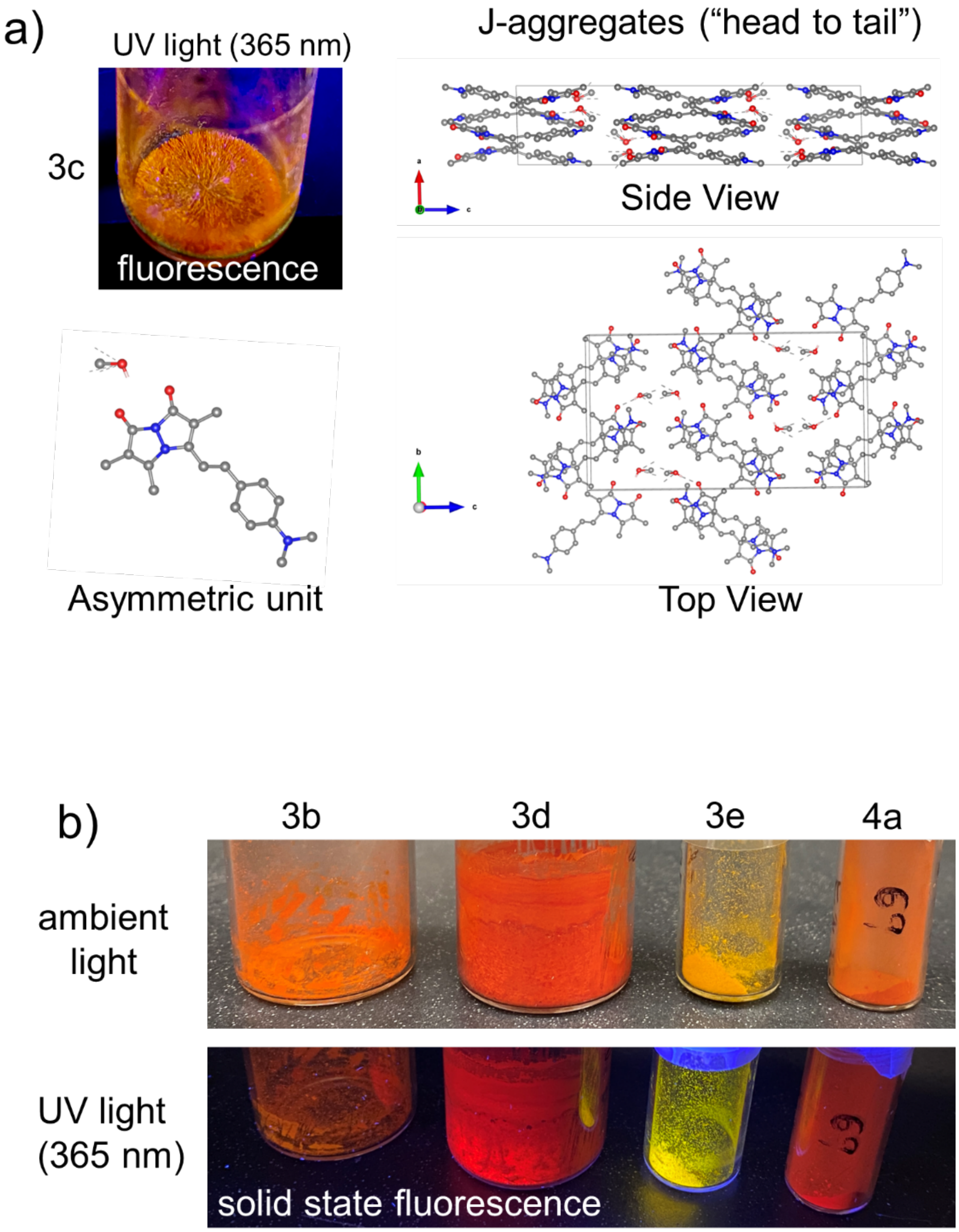


**Figure S16.** (a) Photograph showing the fluorescence in crystalline form of RBF **3c** (R = $NMe_2$) under handheld UV lamp (365 nm), prepared by slow evaporation from a dichloromethane and methanol mixture (8:2 v/v), along with its crystal arrangement (CCDC: 2240198; structure previously reported).[2] Atoms are colored as follows: C, gray; N, blue; O, red. Hydrogen atoms have been omitted for clarity. (b) Top: Photographs of RBFs **3b**, **3d**, **3e**, and **4a** under ambient light; Bottom: Solid-state fluorescence under a handheld UV lamp (365 nm).

#### Mechanistic study of rotational energy barrier $E_a$

Emission intensities for RBFs **3a** and **3c** were recorded in methanol/glycerol mixtures at 25.0˚C, 30.0 ˚C, 34.0 ˚C, 37.0 ˚C and 42.0 ˚C using a Tecan infinite M1000Pro fluorescence microplate reader with temperature control function. RBFs **3a** and **3c** are diluted from 10 mM DMSO stock into a series of methanol and glycerol mixtures (0%, 20%, 40%, 60%, 80%, and 90% glycerol in methanol, 0.25% DMSO) with a final concentration of 25 μM and excited at a wavelength corresponding to their absorption peak. The mixture viscosity ($\eta_{\text{mix}}$) can be calculated using **Eq. 3**.

$$\eta_{\text{mix}} = w_1.\eta_1 + w_2.\eta_2 \tag{3}$$

Where $\eta_{mix}$ stand for the viscosity of the mixture and $\eta_1$, $\eta_2$ are the viscosity of methanol and glycerol, respectively. Factors $w_1$ and $w_2$ stand for the weight fraction of methanol and glycerol. To calculate $w_1$ and $w_2$, the values 0.79 and 1.26 have been used for the density of methanol and glycerol, respectively. The weight fraction can be calculated by using **Eq. 4**. Viscosity values for pure solvents at 25 ˚C were used to estimate the viscosity of the mixtures.

$$\text{Weight Fraction } (w) = (\rho_1 * V_1) / ((\rho_1 * V_1) + (\rho_2 * V_2)) \tag{4}$$

Where $\rho_1$ is the density of glycerol, $V_1$ is the volume fraction of glycerol in the solution, $\rho_2$ is the density of the methanol in the solution, and $V_2$ is the volume fraction of the methanol in the solution.

**Table S6.** Solvent composition and viscosity of methanol and glycerol mixtures.

| Composition Glycerol *vol*% | 0% glycerol | 20% glycerol | 40% glycerol | 60% glycerol | 80% glycerol | 90% glycerol |
|---|---|---|---|---|---|---|
| Viscosity (cP) at 25 ˚C | 0.59 | 268 | 484 | 661 | 809 | 874 |

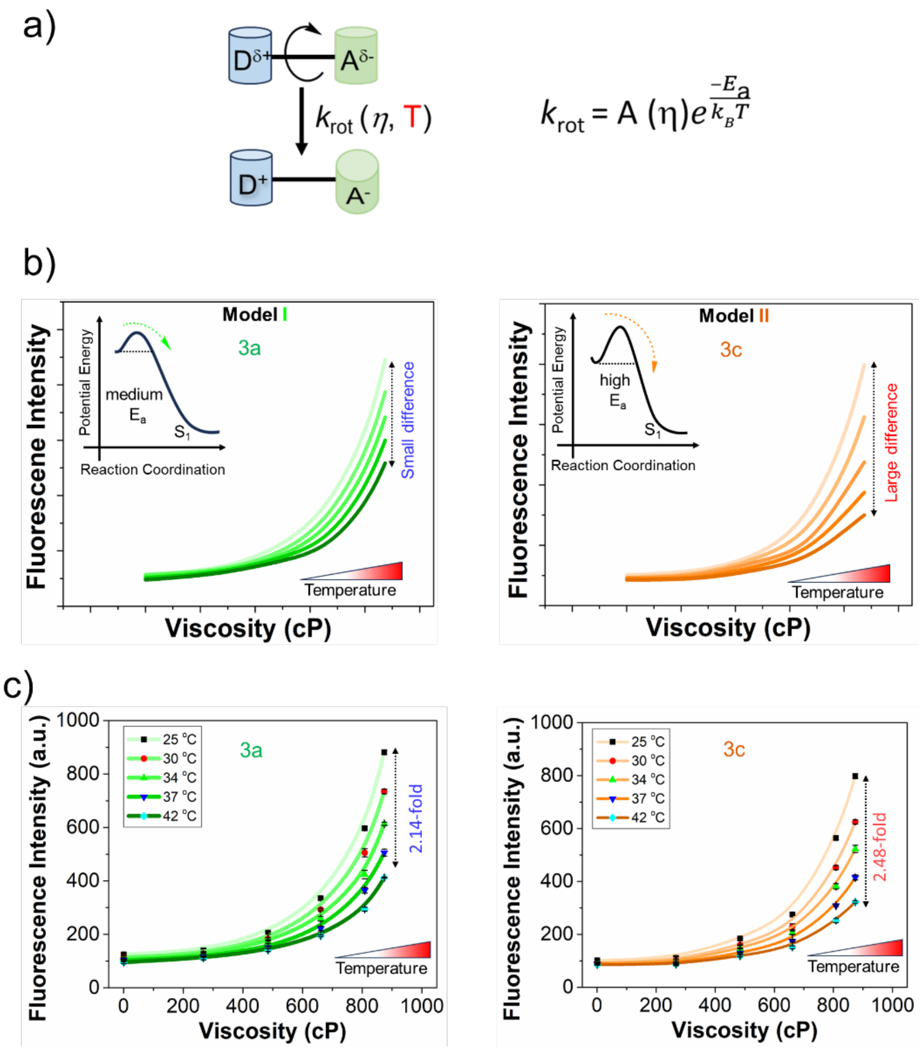


**Figure S17**. Control of viscosity sensitivity of RBFs (**3a** and **3c**) by varying the substitution. (a) Schematic representation of the internal rotation rate ($k_{rot}$), primarily influenced by both temperature and viscosity, according to the Arrhenius equation. (b) Temperature dependence reflects the magnitude of $E_a$. Proposed models suggest that heights of $E_a$ determine the fluorescence intensity pattern in the solvents with different viscosity at different temperatures. Here two models were conceptualized with moderate and high energy barriers for **3a** and **3c**, respectively. (c) Fluorescence pattern of **3a** (left) and **3c** (right) with varying viscosity and temperature. Final concentration of probe is 25 μM in ethylene glycol/glycerol mixtures. Ex/Em: 435 nm/575 nm for **3a** and 463 nm/580 nm for **3c.** Error bars represent SD of 3 measurements.

**αS, Tau 1N4R and A$\beta_{1-42}$fibril preparation:**

We have prepared the αS, Tau$_{1N4R}$ and A$\beta_{1-42}$ fibrils as reported previously[2] and quantified fibril formation by SDS-PAGE.

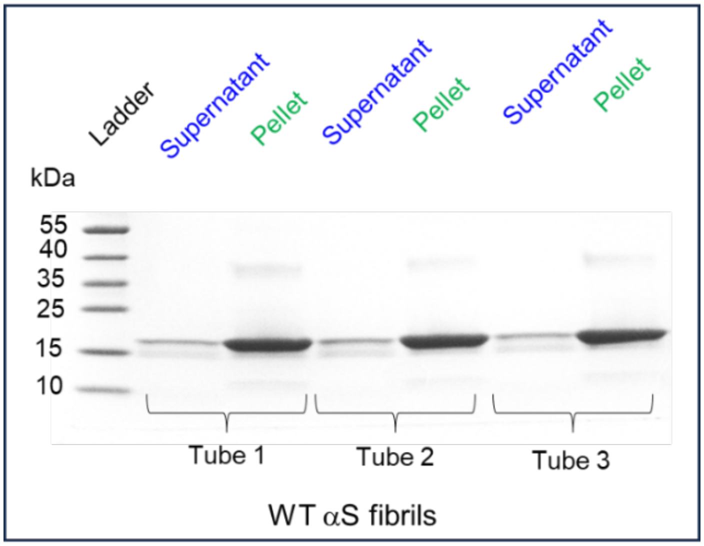


**Figure. S18.** Gel-based quantification of αS protein following fibril formation.

**Absorbance spectra with αS fibrils.** Absorption measurements were performed using Thermo Scientific Genesys 150 UV Vis spectrometer. Absorption spectra of 10 μM of **3b**, **3d**, **3e**, and **4a** (concentrated stock solution prepared as 10 mM in DMSO) in PBS buffer were measured in the presence of 50 μM αS fibrils or folded αS monomer (stock solution 100 μM αS fibrils in PBS buffer).

**Fluorescence spectra with aS fibrils.** Fluorescence measurements were performed using a PTI QuantaMaster™ 40 fluorescence spectrometer. Fluorescence spectra of 10 μM probes concentrated stock solution prepared as 10 mM in DMSO) (**3b**, **3d**, **3e**, and **4a**) in PBS buffer were measured by excitation at their absorbance maximum wavelength in the presence of 50 μM αS fibrils or folded αS monomer (stock solution 100 μM αS fibrils in PBS buffer).

**Excitation spectra with αS fibrils.** Excitation spectral measurements were performed using a PTI QuantaMaster™ 40 fluorescence spectrometer. Excitation spectra of 10 μM solutions of **3b**, **3d**, **3e**, and **4a** (concentrated stock solution prepared as 10 mM in DMSO) in PBS buffer were measured in the presence of 50 μM αS fibrils or folded αS monomer (stock solution 100 μM αS fibrils in PBS buffer).

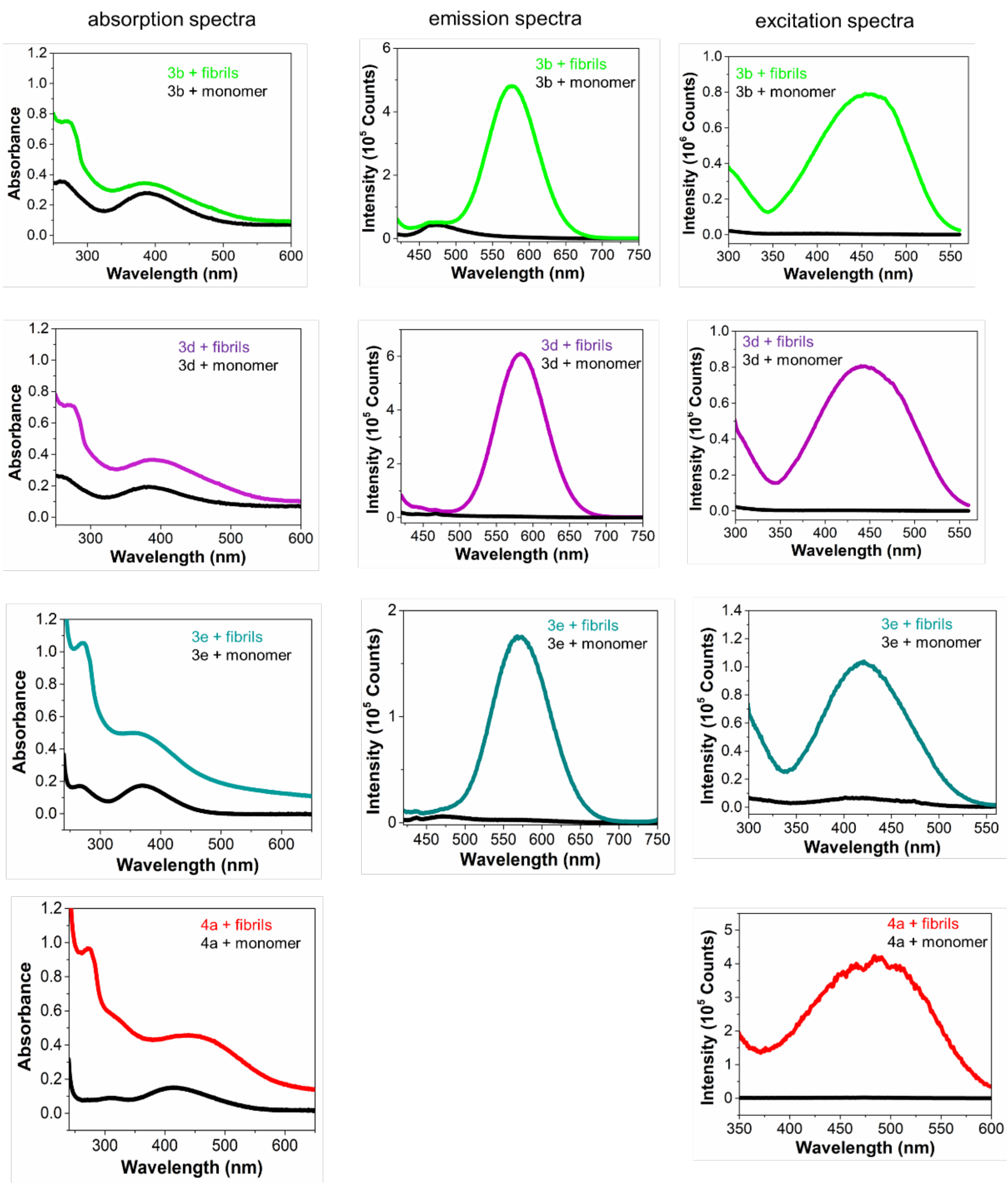


**Figure S19.** Absorption (left), fluorescence (middle); and excitation (right) spectra of **3b**, **3d**, **3e**, and **4a** with $\alpha$S fibrils and folded monomer. The final concentrations of $\alpha$S fibrils and monomer were 50 $\mu$M, and the probe concentration was 10 $\mu$M. Ex: 399 nm for **3b,** 403 nm for **3d,** 386 nm for **3e**, and 436 nm for **4a.** Excitation spectra recorded using emission wavelength 577 nm for **3b**, 584 nm for **3d,** 573 nm for **3e**, and 620 nm for **4a**. Ex/Em slit widths: 3 nm/3 nm.

**Fluorescence lifetime measurements of probes with αS fibrils.** TCSPC measurements of fluorescence lifetime decays for 100 μM solutions of dyes in the presence of 100 μM αS fibrils were collected with the PTI Quantamaster TM 40 using a pulsed LED with a maximum emission at 486 nm. Fluorescence emission was collected at the indicated wavelength for each dye with 20 nm slit widths. The IRF was collected under identical conditions. Data analysis was performed with FluoFit software using an exponential decay model.

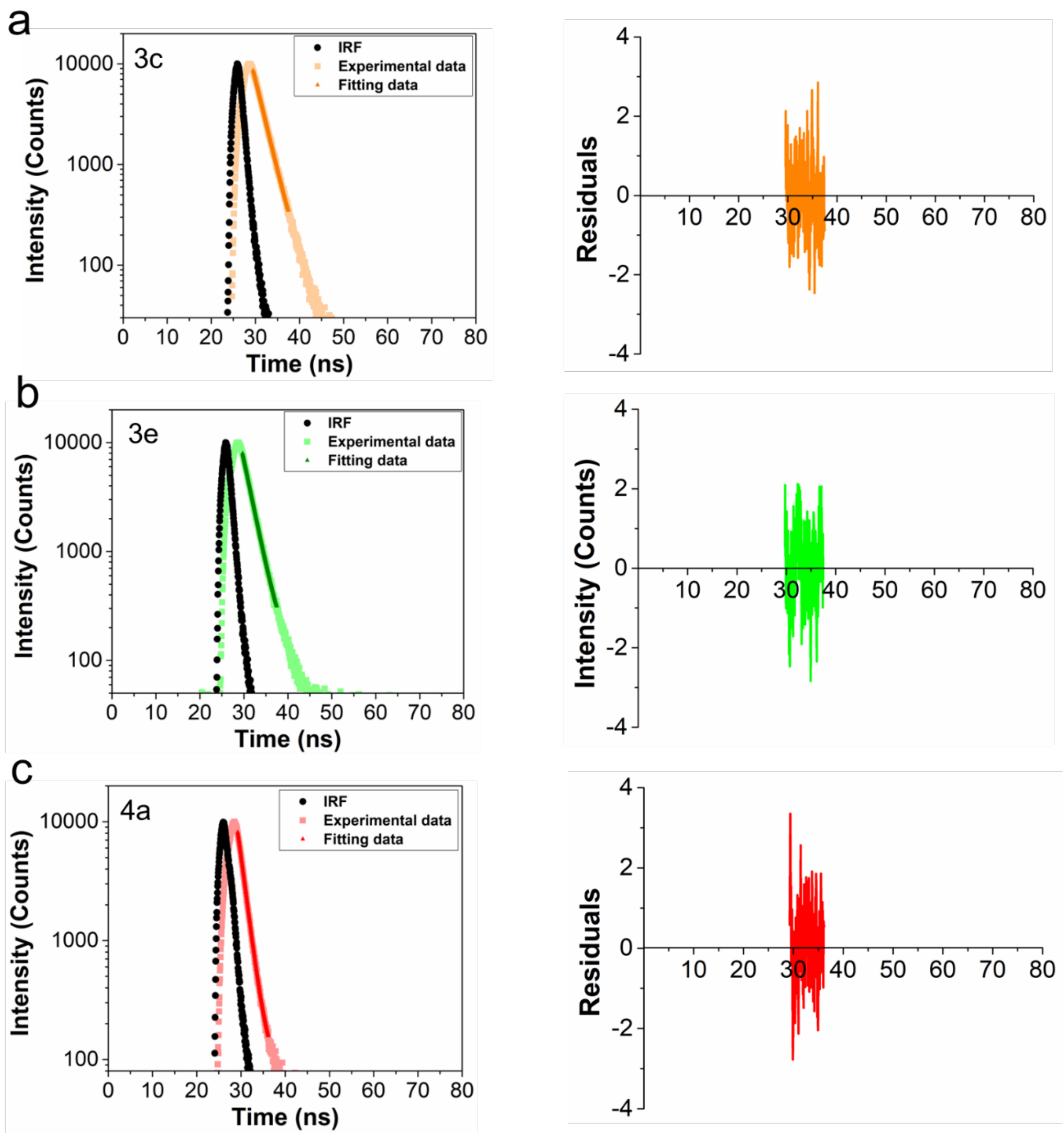


**Figure S20**. Fluorescence lifetime (τ) measurements of **3c** (a), **3e** (b), and **4a** (c) with αS PFFs by time-correlated single photon counting (TCSPC) technique and lifetime was not detected for probe alone. Emission wavelength collected at 580 nm for **3c**, 570 nm for **3e**, and 620 nm for **4a**. Final concentrations of the probe and αS fibrils were 100 μM.

**Table S7**. Fluorescence lifetime measurements of $\alpha$S fibrils with probes **3c, 3e,** and **4a** in PBS buffer and fit $\chi^2$ values.[a]

| Probe | Lifetime (ns) | $\chi^2$ |
|---|---|---|
| **3c** | 2.27 ± 0.01 | 1.079 |
| **3e** | 2.09 ± 0.01 | 1.117 |
| **4a** | 1.27 ± 0.01 | 1.161 |

[a]Emission wavelength collected at 580 nm for **3c**, 570 nm for **3e**, and 620 nm for **4a**. Final concentrations of the probe and $\alpha$S fibrils were 100 $\mu$M.

**Table S8**. Spectral properties of **3b**, **3c**, **3d**, **3e** and **4a** in the presence of $\alpha$S fibrils (+) and folded monomer (–).

| RBF dye | PFFs | $\lambda_{abs}$ (nm)[a] | $\lambda_{ex}$ (nm)[b] | $\lambda_{em}$ (nm)[c] | $\varepsilon$ ($M^{-1}cm^{-1}$)[d] | Stokes shift (nm)[e] | QY (%)[f] | Relative Brightness[g] | Fluorescence Lifetime (ns)[f] |
|---|---|---|---|---|---|---|---|---|---|
| **3b** | - | 390 | - | - | 27430 | - | 0.17 | - | - |
| | + | 384 | 452 | 577 | 34109 | 125 | 34.8 | 254 | n.d |
| **3c** | – | 393 | - | - | 20071 | - | 0.15 | - | - |
| | + | 386 | 463 | 580 | 43864 | 117 | 32.7 | 476 | 2.27 |
| **3d** | – | 386 | - | - | 19048 | - | 0.25 | - | - |
| | + | 389 | 445 | 584 | 36418 | 139 | 37.4 | 286 | n.d |
| **3e** | – | 370 | - | - | 17205 | - | 0.21 | - | - |
| | + | 356 | 424 | 573 | 49869 | 149 | 29.2 | 403 | 2.09 |
| **4a** | – | 416 | - | - | 19024 | - | 0.20 | - | - |
| | + | 442 | 484 | 620 | 52234 | 136 | 27.9 | 383 | 1.27 |

[a]Maximum absorption wavelength ($\lambda_{abs}$) in the presence of $\alpha$S PFFs corresponds to a mixture of bound and unbound dye. [b]Maximum excitation wavelength ($\lambda_{ex}$) better represents the bound form of dye. [c]Maximum emission wavelength ($\lambda_{em}$) corresponds to a bound dye. [d]Molar absorptivity ($\varepsilon$) was measured at $\lambda_{abs}$. [e]Stokes shift was determined as the difference between $\lambda_{ex}$ and $\lambda_{em}$. Final dye and PFF concentrations were 10 and 50 $\mu$M, respectively, for measurements of $\lambda_{abs}$, $\lambda_{ex}$, $\lambda_{em}$, and Stokes shift. [f]Fluorescence quantum yield (QY) (error limit within ± 5) and lifetime measurements were made with 100 μM of dye and PFFs. [g]Relative brightness was determined as the ratio of $\varepsilon$•QY in the presence and absence of $\alpha$S fibrils. $\lambda_{ex}/\lambda_{em}$ = 463/580 nm for **3c,** 424/573 nm for **3e,** and 484/620 nm for **4a**. n.d is not determined.

**Fluorescent QY measurements of 3c and 4a with αS fibrils.** For each QY measurement, the incident excitation light spectrum was collected with 100 μL of probes (100 μM) in PBS buffer. After measuring the incident light intensity, probes (**3c** and **4a**) were added to the αS fibrils (100 μM) from a concentrated stock solution (10 mM in DMSO) to a final concentration of 100 μM and the new spectrum (fluorescence intensity and new incident light intensity) was collected. Using the JASCO Quantum Yield Software, the dye QY was calculated by dividing the dye fluorescence intensity by the difference in incident light intensity in the presence and absence of fibrils.

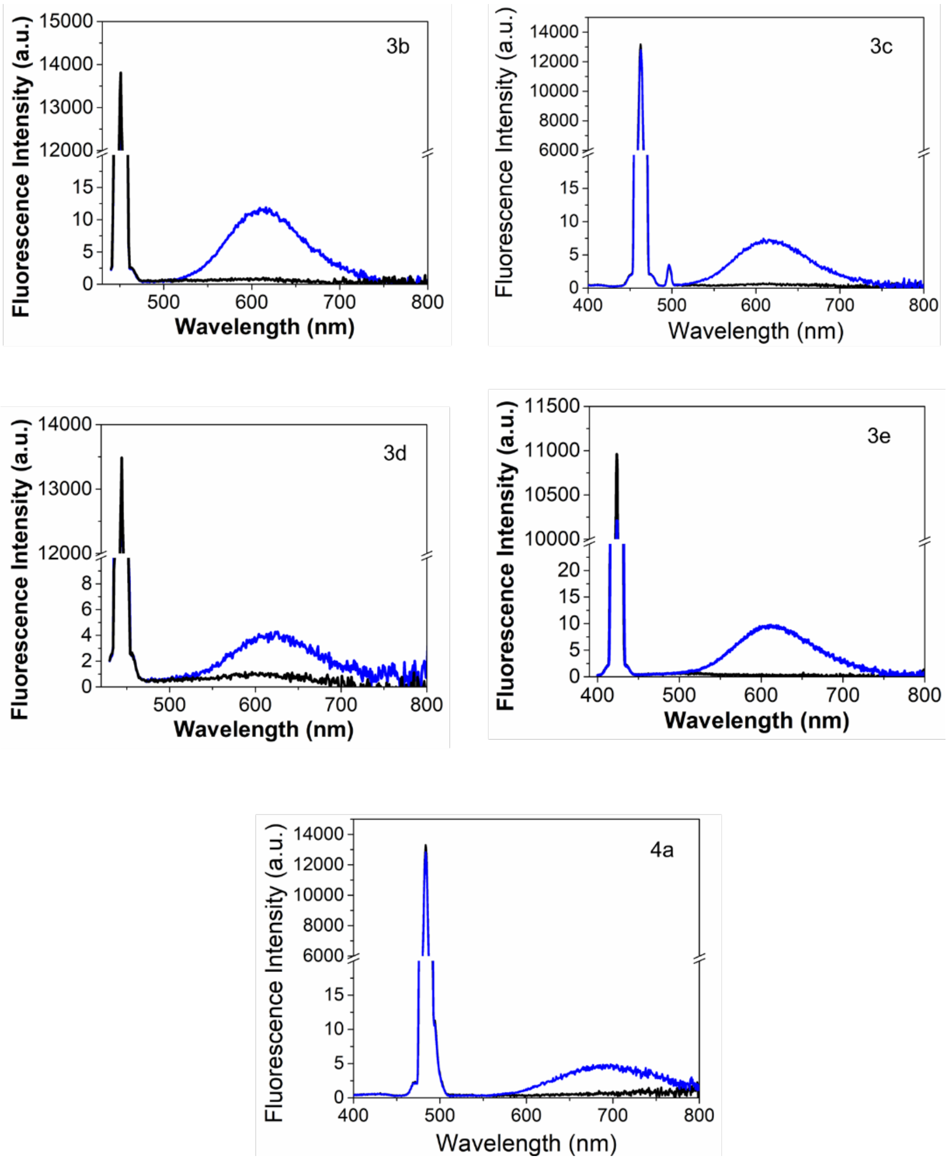


**Figure S21**. Representative QY acquisitions of **3c** and **4a** with αS fibrils. Black line indicates incident light (αS fibrils without probe) and blue line indicates sample (αS fibrils with probe). Excitation at 463 nm for **3c**, and 484 nm for **4a** and spectral collection (400-800 nm). Final concentrations of the probe and αS fibrils are the 100 μM. Ex/Em slit widths: 5 nm/5 nm.

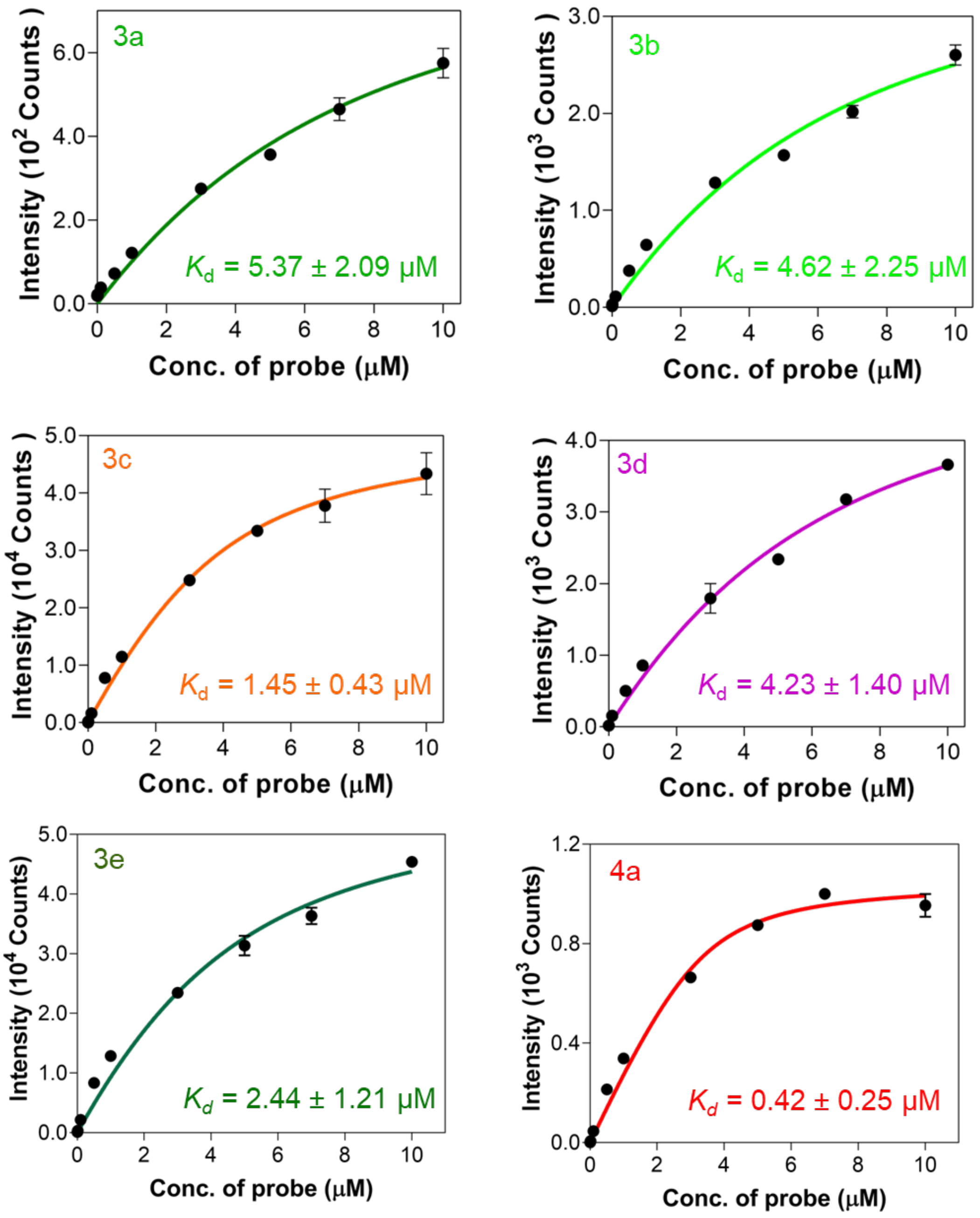


**Figure S22**. Determination of the dissociation constant of probes **3a-e** and **4a** (10 nM-10 μM) binding to 100 μM αS-PFFs. Ex/Em: 435/575 nm for **3a**, 452/577 nm for **3b**, 463/580 nm for **3c,** 445/584 nm for **3d,** 424/573 nm for **3e,** and 484/620 nm for **4a.**

**Displacement Assay of 3c and 4a with known amyloid binding dye ThT:**

αS fibrils (50 μM) in PBS buffer were incubated with 10 μM of ThT (10 mM DMSO stock solutions) for 15 min in Greiner 96 well flat black ½ area plates at 37 ˚C with shaking at 500 rpm for 15 min in an IKA MS3 control orbital shaker (Wilmington, NC, USA). After 15 min of incubation, we competed with varying **3c** concentrations (0, 1, 3, 5, 10, and 20 μM) or **4a** (0, 1, 3, 5, 10, 20, and 30 μM) and again incubated for 15 min. After 15 min incubation, fluorescence intensity measurements were obtained with a Tecan Spark plate reader (Männedorf, Switzerland) by excitation with $\lambda_{ex}/\lambda_{em}$ = 463/580 nm for **3c**, 484/620 nm for **4a,** and 450/482 nm for ThT using the following parameters: excitation and emission bandwidth 5 nm, delay time 0 μs, integration time 40 μs. Similarly, we carried out competition binding of probe **3c** (10 μM) with varying concentrations of BF2846 or Ex-6 (0, 3, 5, 10, and 20 μM), known αS fibril-binding ligands.[1, 5]

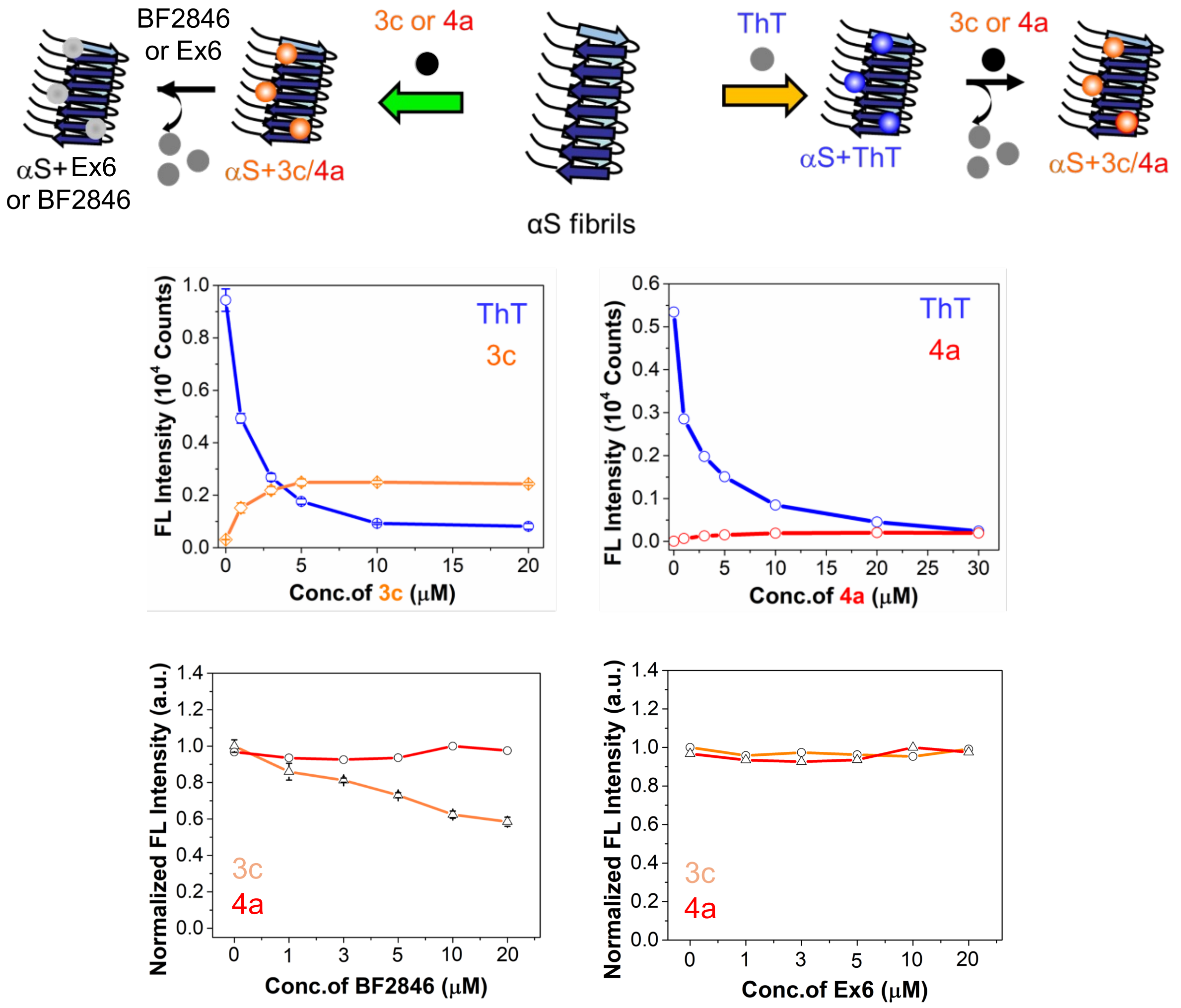


**Figure S23**. Top: Schematic representation of the displacement assay. Bottom: Displacement of the well-known amyloid-binding dye ThT by **3c/4a** or of **3c/4a** by known αS fibril binding ligands BF2846 or Ex-6. Excitation and emission wavelength for fluorescence measurements: Ex/Em: 463 nm/580 nm for **3c,** Ex/Em: 484 nm/620 nm for **4a**, and 450 nm/482 nm for ThT.

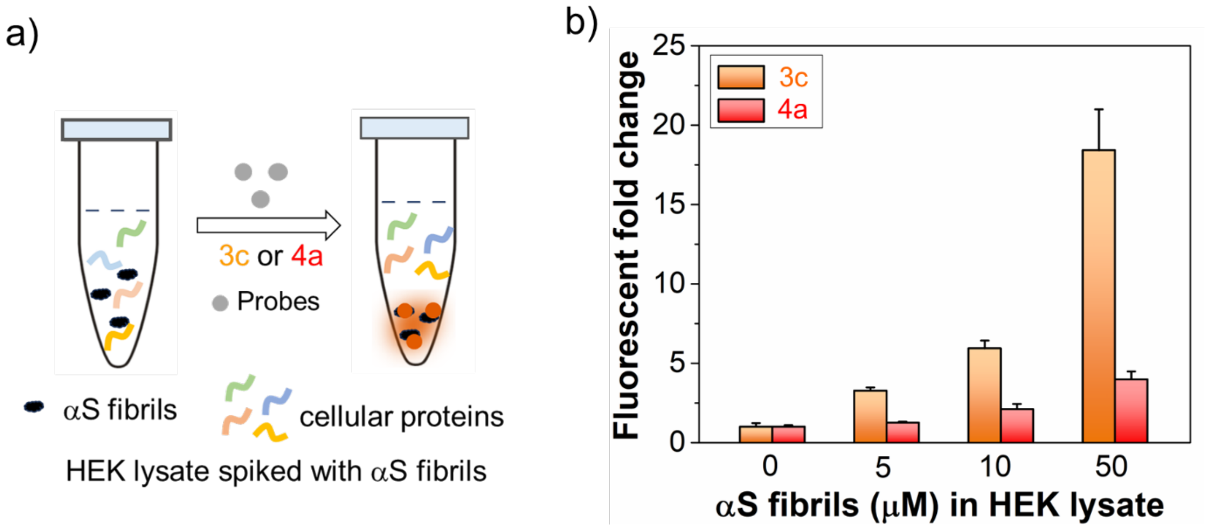


**Figure S24**. (a) Schematic representation of selective detection of αS fibrils in the presence of cellular proteins that are abundant in the cytosolic HEK cell lysate. (b) Fluorescence fold change of **3c** and **4a** (1 μM) with αS PFFs (0 – 50 μM) at 10 mg/mL HEK lysate concentration.

### Fluorescence and Excitation Spectral Measurements of 4a with Tau and $Ab_{1-42}$ Fibrils

Fluorescence measurements were performed using a PTI QuantaMaster™ 40 fluorescence spectrometer. 1) Fluorescence or Excitation spectra of 5 μM of **4a** (concentrated stock solution prepared as 10 mM in DMSO) in PBS buffer were measured in the absence and presence of 25 μM tau fibrils (stock solution 32 μM tau fibrils in PBS buffer) and 2) Fluorescence or Excitation spectra of 10 μM of **4a** (concentrated stock solution prepared as 10 mM in DMSO) in PBS buffer were measured in the absence and presence of 50 μM $A\beta_{1-42}$ fibrils (stock solution 1 mM $A\beta_{1-42}$ in 10 mM phosphate buffer, pH 7.4).

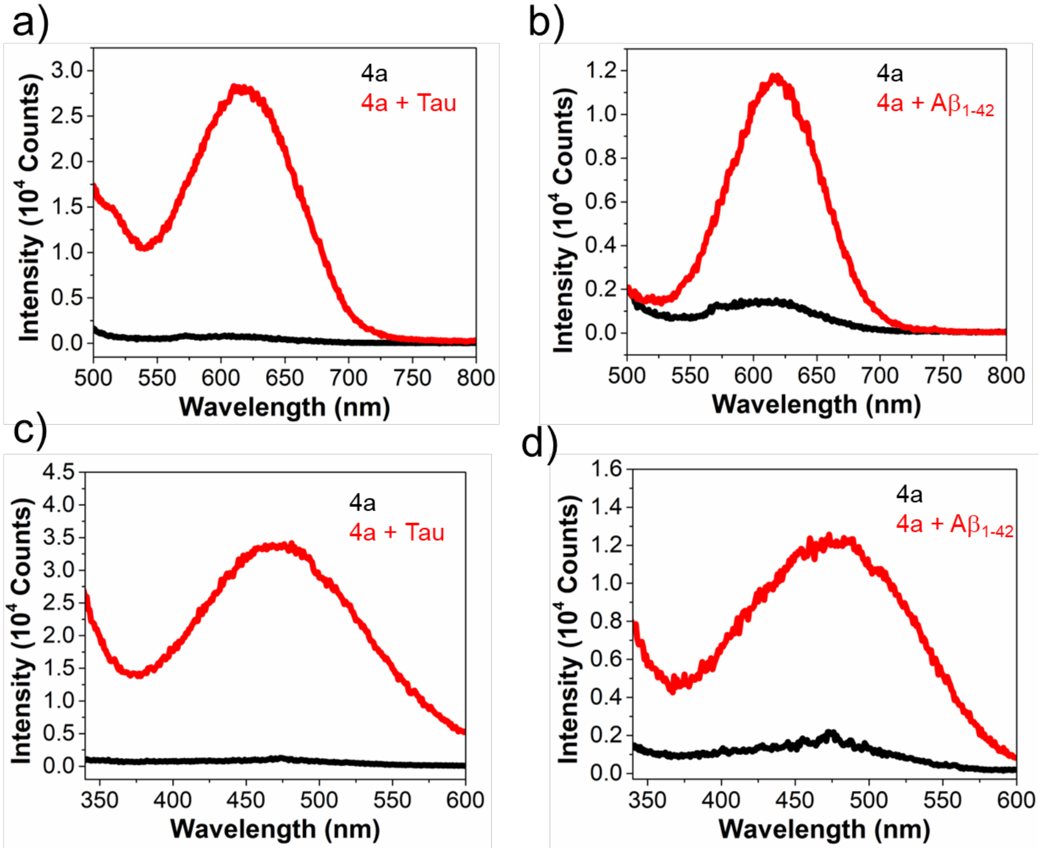


**Figure. S25**. Fluorescence spectra (a-b) and excitation spectra (c-d) of **4a** with and without tau and $A\beta_{1-42}$ fibrils. Final probe and tau fibril concentrations were 5 μM and 25 μM, respectively. Final probe and $A\beta_{1-42}$ fibril concentrations were 10 μM and 50 μM, respectively. Excitation at 484 nm for **4a**. Emission wavelength is 620 nm for **4a** . Ex/Em slit widths: 3 nm/3 nm.

**Fractionation Assay of 3c and 4a:** αS fibrils (50 μM) in PBS buffer were incubated with 10 μM of either probe **3c** or **4a** (from 10 mM DMSO stock solutions) in Greiner 96-well flat black ½ area plates at 37 °C with shaking at 500 rpm for 15 minutes on an IKA MS3 control orbital shaker (Wilmington, NC, USA). Following incubation, 100 μL of the mixture was collected for total fluorescence measurements. The samples were then centrifuged at 13,000 rpm for 30 minutes. After centrifugation, the supernatant and pellet fractions were carefully separated. The pellet was resuspended in an equal volume of PBS buffer to its original sample volume. Fluorescence intensity measurements were recorded for the total (T), supernatant (S), and pellet (P) fractions using a Tecan Spark plate reader (Männedorf, Switzerland) with the following settings: excitation/emission wavelengths of 463/580 nm for **3c** and 484/620 nm for **4a**, excitation and emission bandwidths of 5 nm, delay time of 0 μs, and integration time of 40 μs.

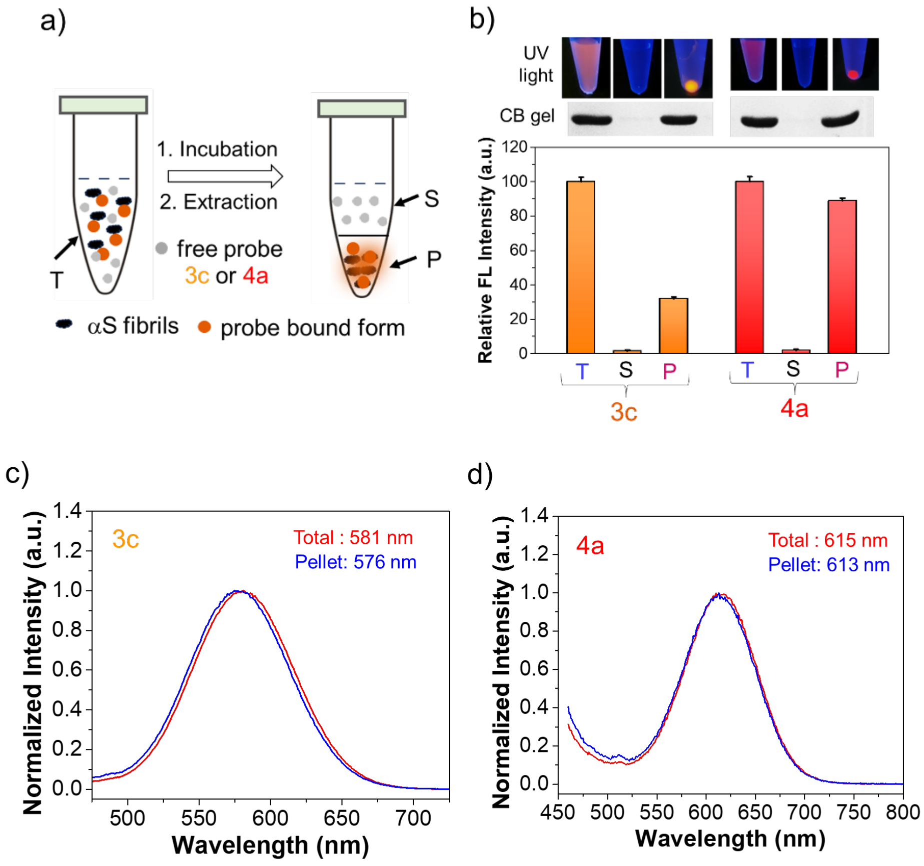


**Figure S26**. (a) Schematic representation of selective detection of αS fibrils over monomer. (b) Fractionation of αS fibrils revealed that **4a** retained its relative fluorescence in the pellet fraction more than **3c**, T: total; S: supernatant; P: Pellet; top: Coomassie blue gel of T, S, P fractions and their photographs fractions under handheld UV light (365 nm). $\lambda_{ex}/\lambda_{em}$ = 463/580 nm for **3c** and 484/620 nm for **4a**. Error bars represent SD of 3 measurements. (c-d) Fluorescence spectra of total and pellet fractions of **3c** (left) and **4a** (right). Ex: 418 nm for **3c** and 436 nm for **4a.**

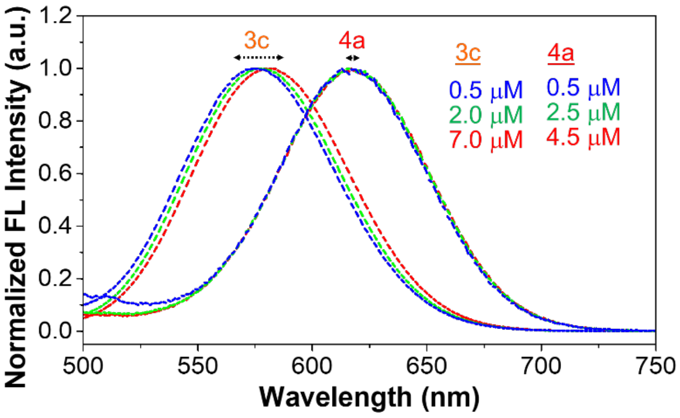


**Figure S27**. Concentration-dependent shift in the emission maxima of probes **4a** and **3c** in the presence of αS fibrils (50 μM). Ex: 418 for **3c** and 436 for **4a**.

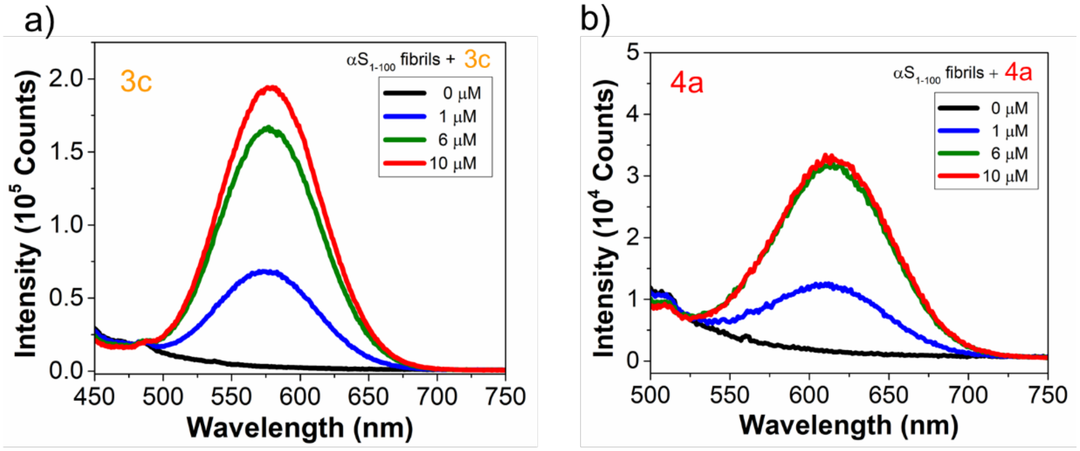


| 3c (μM) | 1 | 6 | 10 |
|---|---|---|---|
| $\lambda_{max}$(nm) | 574 | 577 | 578 |

| 4a (μM) | 1 | 6 | 10 |
|---|---|---|---|
| $\lambda_{max}$(nm) | 610 | 614 | 615 |

**Figure S28**. Dose-dependent fluorescence 'turn-on' of **3c** and **4a** with C-terminal truncated $\alpha S_{1-100}$ fibrils (50 μM). Top: Fluorescence spectra of varied probe concentration with C-terminal truncated αS fibrils, and bottom: tabulated $\lambda_{max}$ ($\lambda_{em}$) at varied **3c** concentration with C-terminal truncated $\alpha S_{1-100}$ fibrils (a) **3c** and (b) **4a**. Ex: 418 for **3c** and 436 for **4a**.

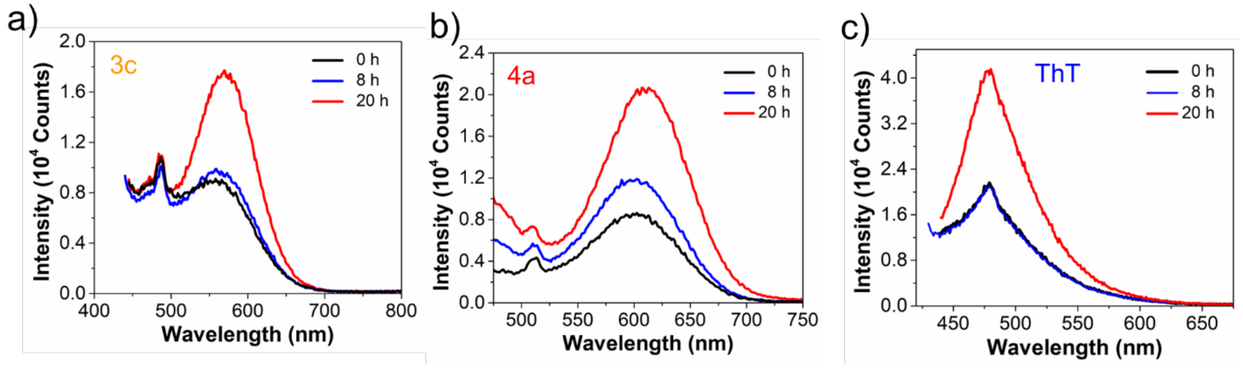


**Figure S29**. Time-dependent $\alpha$S aggregation monitored by RBFs **3c**, **4a,** and ThT. Fluorescence spectra recorded at different time intervals (0, 8, and 20 h). (a) **3c**, (b) **4a**, and (c) ThT. Ex: 418 for **3c**, 436 for **4a,** and 412 nm for ThT. Final probe concentration is 10 $\mu$M and $\alpha$S monomer concentration is 50 $\mu$M.

**Experimental procedure for the fluorescent measurements of 4a with TMAO:** A 1 mM stock solution of **4a** in DMSO and a 6.5 M stock solution of trimethylamine *N*-oxide (TMAO) in Tris buffer at pH 7.4 were prepared. From these stocks, the fluorescence measurements of probe **4a** (10 $\mu$M) were carried out with and without αS monomer (50 $\mu$M) at different concentrations of TMAO (2 M, 4 M and 6 M) diluted with the Tris buffer. The solutions were mixed and then the spectra were measured using a Photon Technology International (PTI) QuantaMaster™ 40 fluorescence spectrometer with excitation at 436 nm.

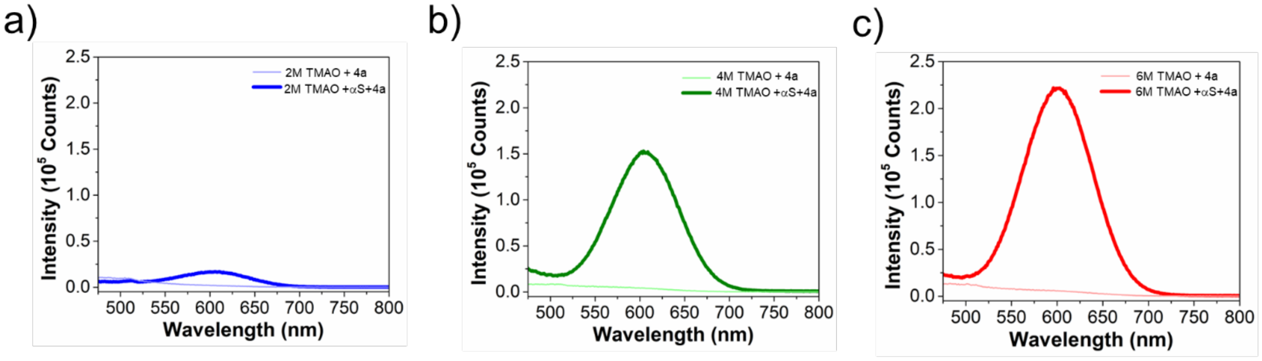


d)

| Conc. of TMAO (M) | 2 | 4 | 6 |
|---|---|---|---|
| $\lambda_{max}$ of 4a | 604 | 606 | 601 |

**Figure. S30**. Fluorescence spectra of probe **4a** (10 $\mu$M) with and without $\alpha$S monomer (50 $\mu$M) in different TMAO concentrations. (a) 2 M TMAO, (b) 4 M TMAO, (c) 6 M TMAO, and tabulated $\lambda_{max}$ ($\lambda_{em}$) of **4a** with increasing TMAO concentrations. Excitation wavelength for **4a** is 436 nm. Ex/Em slit widths: 3 nm/3 nm.

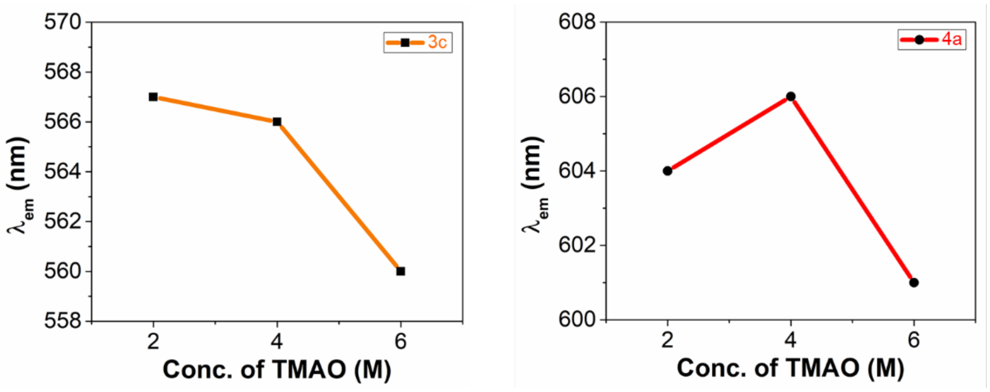


**Figure. S31**. Emission maximum ($\lambda_{em}$) shift of **3c** and **4a** (10 μM) with αS monomer (50 μM) in different 2 M, 4 M, and 6 M TMAO concentrations. Left: The gradual decrease in the $\lambda_{em}$ with an increase in the TMAO concentration suggests that viscosity increases. Right: The initial increase in the $\lambda_{em}$ from 2 M to 4 M TMAO concentration suggests that polarity increases, followed by the decrease in the $\lambda_{em}$ at 6 M TMAO suggests that viscosity increase. Excitation wavelength for **3c** is 418 nm and for **4a** is 436 nm. $\lambda_{em}$ of **3c** with αS monomer (50 μM) at 2 M, 4 M, and 6 M TMAO were previously reported.[1]

### Fluorescence Measurements with Patient Tissue Lysate and Amplified αS Fibril Samples

Fluorescence measurements of probe **4a** were conducted with tissue samples from three PDD patient cases (PDD1, PDD2, and PDD3), comparing the results with probe **3c** and **ThT** that were previously reported.[2] Probe **4a** binding was assessed at a 7 μM stock concentration of patient tissue lysates, prepared under two conditions: lysate only and amplified αS fibrils (AFs), as described previously.[2] For each condition, samples were diluted to a final concentration of 1 μM in a Greiner 384-well small volume microplate, using PBS buffer to reach a final volume of 5 μL per well. Probe **4a** was then added at a 1 μM final concentration (from a 5 mM DMSO stock solution), and samples were incubated at 37 °C with shaking at 500 rpm for 15 minutes on an IKA MS3 control orbital shaker. After incubation, fluorescence intensity measurements were taken using a Tecan Spark plate reader with the following settings: excitation at 484 nm and emission at 620 nm (5 nm bandwidth for both), 0 μs delay time, and 40 μs integration time.

**Experimental procedure for liquid-liquid phase separated protein condensates:**

**$\alpha$S/poly L-lysine (pLK) protein condensates:**

**Stock and working solutions:** We first prepared the liquid-liquid phase separation (LLPS) buffer that consists of 20 mM HEPES with 150 mM NaCl and 1 mM TCEP, adjusted to pH 7.4 supplemented with 30% (w/v) PEG 8000 stock solution by dissolving 3 g of PEG 8000 in 10 mL of the LLPS buffer. We purchased the commercially available Poly-L-lysine hydrobromide (pLK) (mol wt. 15,000-30,000) (Product Number: P7890) and FITC-labelled poly-L-lysine ($pLK_{FITC}$) (mol wt. 15,000-30,000) (Product Number: P3543) and dissolved 1.2 mg in 120 $\mu$L of HEPES buffer to make a 1 mg/mL stock solution. We transiently expressed the wild-type (WT) $\alpha$S and the fluorescently labeled $\alpha$S at position 94 with acridon-2-yl alanine (Acd), termed as $\alpha S_{94Acd}$, by genetic code expansion by using a previously described procedure.[6] The desired protein condensate solution (25 $\mu$L) was prepared with WT $\alpha$S (100 $\mu$M) and pLK (0.25 mg/mL) having 5% $\alpha S_{94Acd}$ and 5% $pLK_{FITC}$ with 15% of PEG 8000 in LLPS buffer. The reaction mixture was gently mixed by pipetting up and down to ensure homogeneity. Then the reaction mixture was incubated for 5 min at room temperature before imaging.

**$\alpha$S/$Tau_{0N4R}$ protein condensates:**

**Stock and working solutions:** We expressed $Tau_{0N4R}$ using a previously described procedure[7] and labeled it with 5 equivalents of Alexa Fluor™ 488 NHS Ester (Succinimidyl Ester) in Ni-NTA Buffer C (25 mM Tris, pH 8.0; 100 mM NaCl; 1 mM EDTA; and 1 mM TCEP) at 4 °C with gentle rocking overnight. The protein was desalted using a Trap desalting column to isolate labelled protein from free dye, termed as $AF_{488}$-$Tau_{0N4R}$. The protein concentrations of both the labeled ($AF_{488}$-$Tau_{0N4R}$) and unlabeled ($Tau_{0N4R}$) constructs were quantified using a Nanodrop spectrophotometer. The desired protein condensate solution (25 $\mu$L) prepared with WT $\alpha$S (80 $\mu$M) and $Tau_{0N4R}$ (40 $\mu$M) having 5% $\alpha S_{94Acd}$ and 5% $AF_{488}$-$Tau_{0N4R}$ and 15% of PEG 8000 in LLPS buffer (20 mM HEPES, 150 mM NaCl, 1 mM TCEP, pH 7.4). The reaction mixture was gently mixed by pipetting up and down to ensure homogeneity. Then the reaction mixture was incubated for 5 min at room temperature before imaging.

**Confocal laser scanning microscopy (CLSM) imaging of protein condensates by RBFs 3c and 4a:**

Confocal microscopy imaging was performed using an Olympus IX83 inverted microscope, equipped with FluoView3000 scanning system(Olympus, Center Valley, PA). Images were taken at room temperature using a 60x 1.2 NA water immersion objective lens (Olympus). Imaging of multi-protein systems was performed *via* orthogonal fluorescence labeling of proteins using Alexa 488 ($\lambda_{ex}$ 488 nm, $\lambda_{em}$ 500–540 nm), Acd ($\lambda_{ex}$ 405 nm, $\lambda_{em}$ 420–480 nm), and FITC ($\lambda_{ex}$ 488 nm, $\lambda_{em}$ 500 –540 nm), RBF **3c** ($\lambda_{ex}$ 561 nm, $\lambda_{em}$ 570–620 nm), RBF **4a** ($\lambda_{ex}$ 561 nm, $\lambda_{em}$ 630–680 nm). The excitation lasers were alternated to minimize crosstalk between the different channels using a sequential line scan mode. Images were analyzed with ImageJ (version 1.52a) and Python (version 3.7.1) programs. For imaging droplets in solution, glass coverslips (25 × 25 $mm^2$, Fisher Scientific) were passivated with BSA (2 mg/mL in HEPES buffer). Solutions (10 µL) containing the droplets incubated with the RBFs **3c**, **4a**, and ThT (10 µM) for 5 min and were imaged in a closed chamber created by sandwiching two coverslips (25 × 25$mm^2$, Fisher Scientific) using vacuum grease.

**CLSM imaging of αS fibrils with RBFs 3c, 4a, and ThT:** Confocal microscopy imaging of αS fibrils with RBFs **3c**, **4a**, and ThT (10 µM) was performed via the same confocal set up as described above. For the analysis, 100 µM αS fibrils were used for αS/pLK protein condensates, and 80 µM αS fibrils were used for αS/$Tau_{0N4R}$ protein condensates.

**Fluorescence recovery after photobleaching (FRAP) of αS/poly-L-lysine (pLK) and αS/$Tau_{0N4R}$ condensates:**

FRAP experiments of αS/pLK and αS/$Tau_{0N4R}$ condensates were performed with the same confocal set up as described above. A circular region of interest (ROI) within a protein condensate settled on the glass coverslip was bleached using short exposures (~ 80 s) of 405 nm lasers at 100% laser power. The collection of images for the recovery stage started immediately after the bleaching. FRAP data were analyzed using ImageJ (for image intensity extraction) and Microsoft Excel (for quantitative analysis). For each time frame, mean intensities were estimated for an ROI within the bleached region and of another ROI from the unbleached region. The ratio of intensities at bleached ($I_{bleach}(t)$) and unbleached regions ($I_{unbleach}(t)$) was determined at each time point and normalized to 1 for the intensity ratio before photobleaching ($q(t_{prebleach})$) and to 0 for the intensity ratio at 0 s after photobleaching ($q(t_0)$) using the following formula:

$$I_{norm}(t) = \frac{q(t) - q(t_0)}{q(t_{prebleach}) - q(t_0)}$$

$$where\ q(t) = \frac{I_{bleach}(t)}{I_{unbleach}(t)}$$

The normalized intensities were plotted against time to obtain a fluorescence recovery profile. The recovery profile was fit to a single exponential model as described elsewhere**.**

$$I(\mathrm{t}) = \mathrm{A}(1 - \mathrm{e}^{-\mathrm{t}/\mathrm{t}})$$

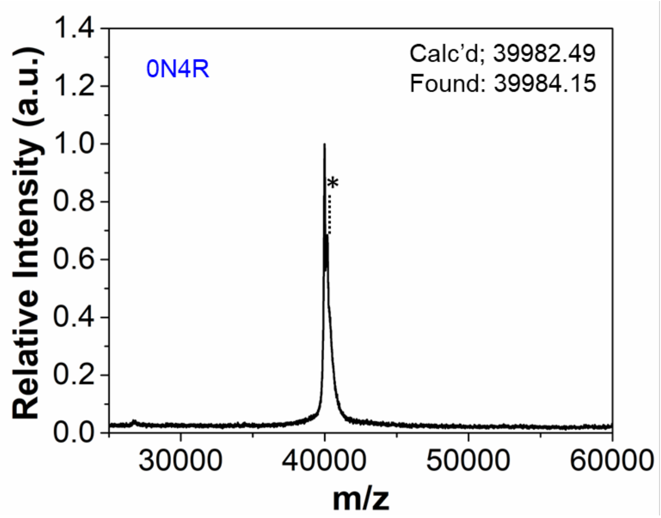


**Figure S32.** MALDI MS characterization of $Tau_{0N4R}$. On the plot matrix adduct peaks are marked with *.

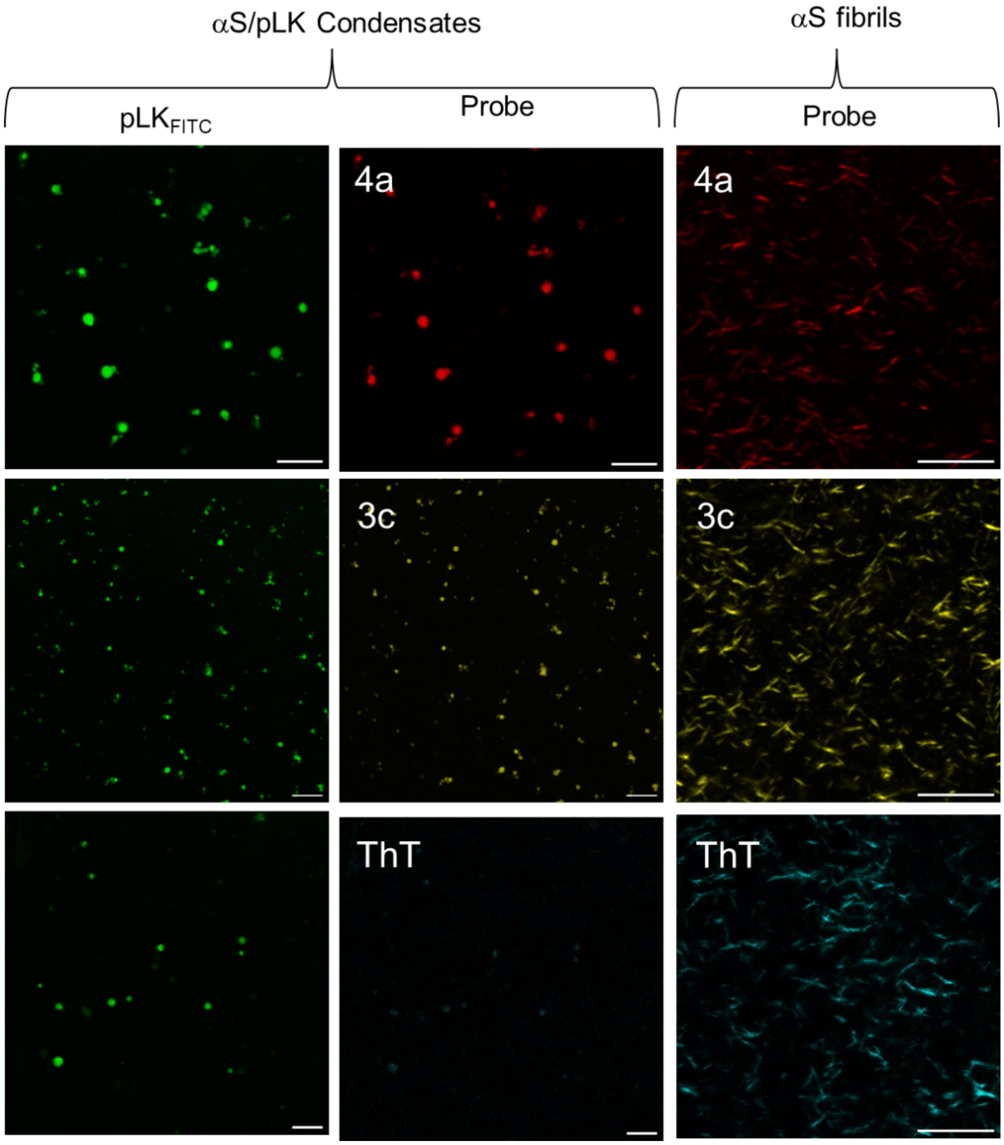


**Figure. S33. Fluorescence imaging of αS/pLK protein condensates *in vitro*.** Left: Confocal laser scanning microscopy (CLSM) images of WT αS/pLK condensates at a concentration of 100 μM αS and 0.25 mg/mL of pLK in LLPS buffer (doped with 5% FITC-labeled pLK) uptake by RBFs **3c**, **4a,** and ThT. Right: Right: CLSM images of WT αS fibrils uptake by RBFs **3c**, **4a**, and ThT. Ex: 488 nm and Em: 500-540 nm for FITC-labeled pLK, Ex: 561 nm and Em: 570-620 nm for **3c**, Ex: 561 nm and Em: 630-680 nm for **4a**, and Ex: 405 nm and Em: 430-480 nm for ThT. All experiments were performed in LLPS buffer. The scale bars are 10 μm.

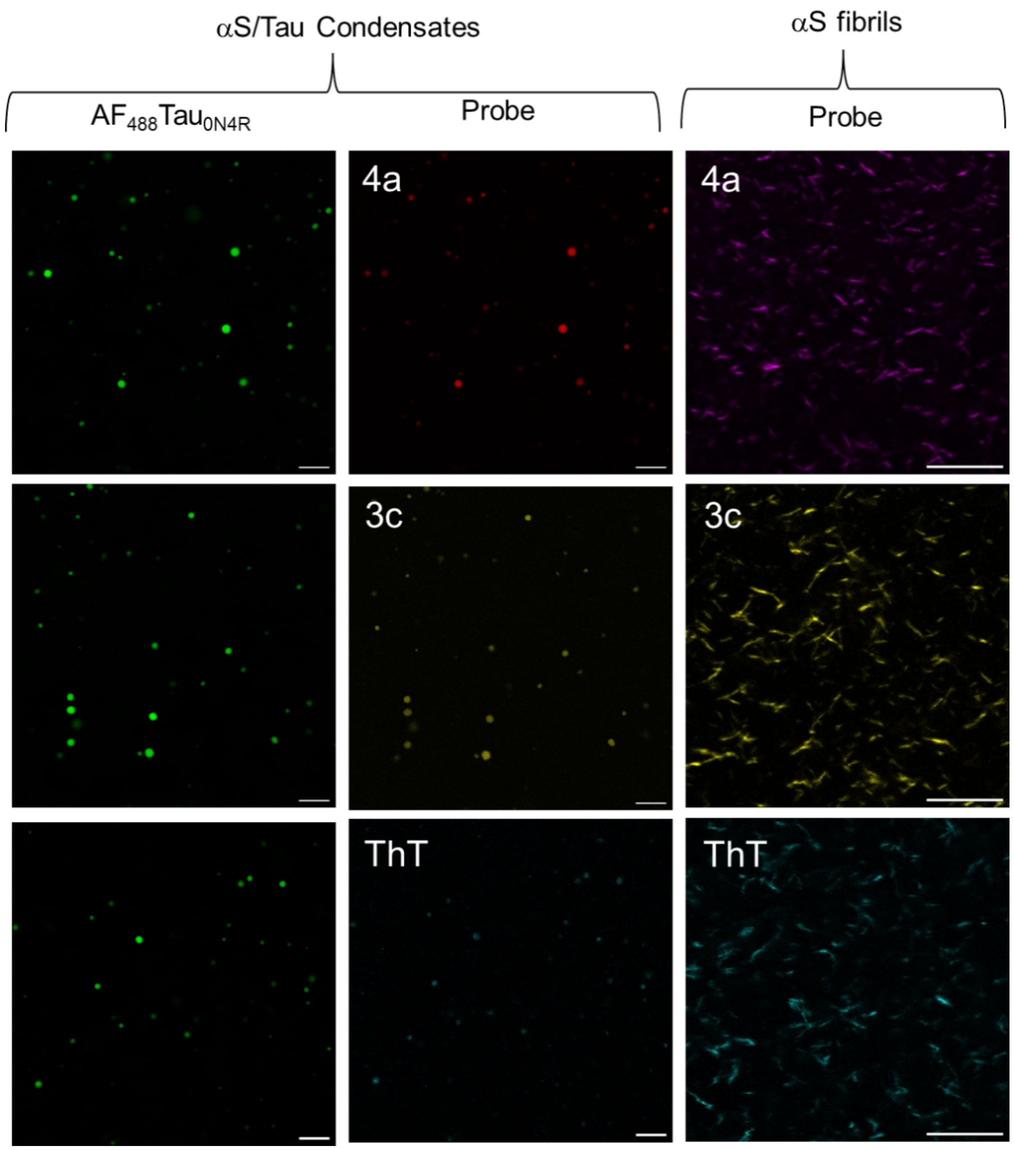


**Figure. S34. Fluorescence imaging of αS/Tau$_{0N4R}$ protein condensates *in vitro*** : Left: CLSM images of WT αS/Tau$_{0N4R}$ condensates (80 μM of αS and 40 μM of Tau in LLPS buffer) uptake by RBFs **3c**, **4a**, and ThT (doped with 5% AF$_{488}$-labeled Tau$_{0N4R}$). Right: CLSM images of WT αS fibrils uptake by RBFs **3c**, **4a**, and ThT. Ex: 488 nm and Em: 500-540 nm for AF$_{488}$-labeled Tau$_{0N4R}$, Ex: 561 nm and Em: 570-620 nm for **3c**, Ex: 561 nm and Em: 630-680 nm for **4a**, and Ex: 405 nm and Em: 430-480 nm for ThT. All experiments were performed in an LLPS buffer: 20 mM HEPES, 150 mM NaCl, and 1 mM TCEP at pH 7.4, with 15% PEG-8000, at room temperature. The scale bars are 10 μm.

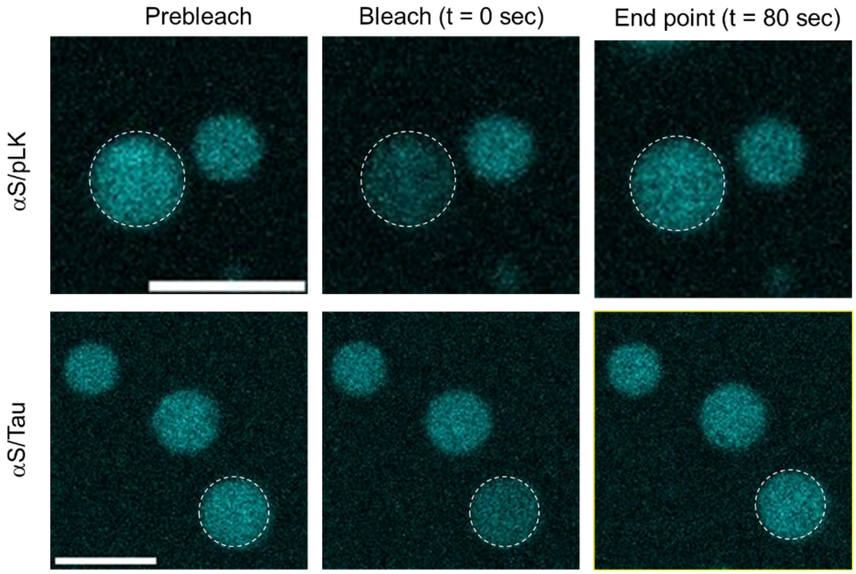


**Figure. S35**. Representative images of fluorescence recovery after photobleaching (FRAP) of αS within αS/pLK (top) and αS/Tau (bottom) condensates (doped with 5% Acd-labeled αS). The white circle indicates the bleached area. Ex: 405 nm and Em: 430-480 nm for Acd-labeled αS. Scale bars are 5 μm.

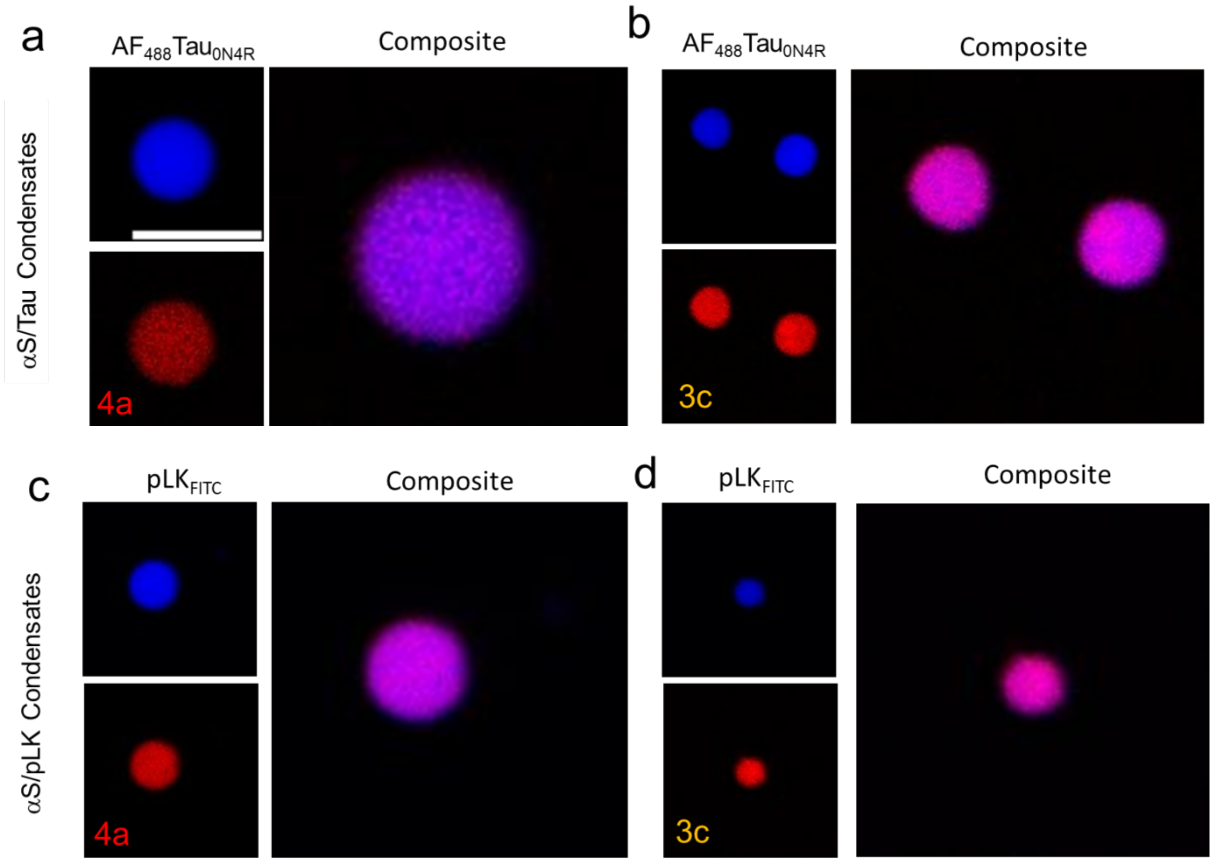


**Figure. S36**. RBF **4a,** exhibiting the lowest viscosity sensitivity, enables more diffusive staining of αS/pLK and αS/Tau condensates compared to **3c**. (a-b) CLSM imaging of αS/Tau condensates stained with **3c** and **4a**. (c-d) CLSM imaging of αS/pLK condensates stained with **3c** and **4a**. Ex: 488 nm and Em: 500-540 nm for $AF_{488}$-labeled $Tau_{0N4R}$ or FITC labeled pLK. Ex: 561 nm and Em: 570-620 nm for **3c**, Ex: 561 nm and Em: 630-680 nm for **4a**. The scale bar is 5 μm.

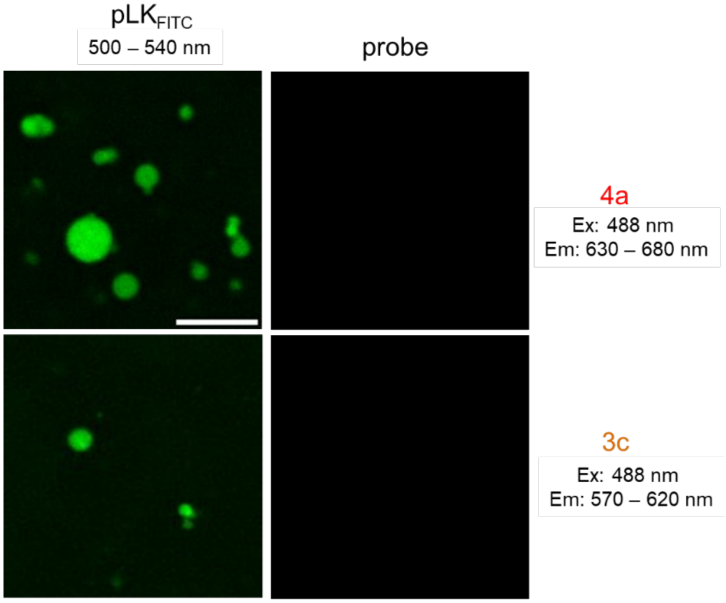


**Figure. S37**. Fluorescence imaging control to verify absence of significant bleed-through of αS/pLK condensates by **4a** and **3c**. Ex: 488 nm and Emission collected for 500-540 nm for $pLK_{FITC}$, 570-620 nm for **3c**, and Em: 630-680 nm for **4a**. The scale bar is 10 μm.

**Cartesian coordinates of the optimized geometries: 3a**

Cartesian coordinates of **3a** optimized at **# APFD/6-311+G (2d, p)** at ground state[2]

optimized at **# APFD/6-311+G (2d, p)** at first excited state

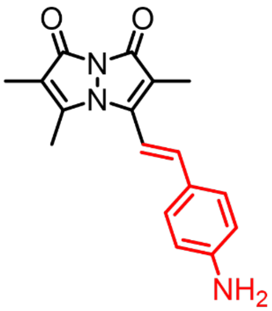

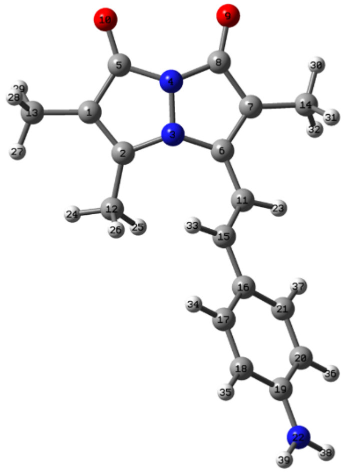

| Atom | X | Y | Z | | Atom | X | Y | Z |
|---|---|---|---|---|---|---|---|---|
| C | 3.111515 | -1.862805 | 0.179518 | | C | -3.480191 | 0.837823 | 0.600434 |
| C | 1.805149 | -1.461545 | 0.285987 | | N | -6.911188 | -0.254800 | 0.043936 |
| N | 1.706191 | -0.155576 | -0.155364 | | H | -0.946236 | 1.806117 | 0.649072 |
| N | 2.973657 | 0.302745 | -0.429244 | | H | 0.990741 | -3.111533 | 1.314970 |
| C | 3.904004 | -0.742530 | -0.245796 | | H | 0.100549 | -1.601149 | 1.558301 |
| C | 0.899110 | 0.941390 | 0.149764 | | H | -0.068858 | -2.492966 | 0.047720 |
| C | 1.710926 | 2.087433 | 0.128886 | | H | 2.972347 | -3.874490 | 0.913702 |
| C | 3.038234 | 1.709261 | -0.235107 | | H | 4.065046 | -3.673934 | -0.456663 |
| O | 4.060739 | 2.384668 | -0.395266 | | H | 4.552602 | -3.115142 | 1.138262 |
| O | 5.119363 | -0.639534 | -0.432917 | | H | 2.144978 | 4.151789 | 0.256086 |
| C | -0.505217 | 0.922340 | 0.197309 | | H | 0.567730 | 3.803191 | -0.467181 |
| C | 0.643510 | -2.202733 | 0.824708 | | H | 0.784786 | 3.649885 | 1.273630 |
| C | 3.699048 | -3.199277 | 0.459708 | | H | -0.890668 | -0.809384 | -0.974359 |
| C | 1.277863 | 3.491536 | 0.308270 | | H | -2.963565 | -1.746156 | -1.569467 |
| C | -1.336139 | -0.033312 | -0.361532 | | H | -5.407543 | -1.860976 | -1.416628 |
| C | -2.748400 | -0.062604 | -0.227006 | | H | -5.382317 | 1.477781 | 1.320757 |
| C | -3.496668 | -1.040231 | -0.940442 | | H | -2.952111 | 1.583501 | 1.183148 |
| C | -4.859329 | -1.108815 | -0.858773 | | H | -7.428959 | 0.383896 | 0.626586 |
| C | -5.573081 | -0.194222 | -0.040463 | | H | -7.437017 | -0.946811 | -0.466049 |
| C | -4.844160 | 0.779747 | 0.687582 | | | | | |

**Cartesian coordinates of the optimized geometries: 3c**

Cartesian coordinates of **3c** optimized at **# APFD/6-311+G (2d, p)** at ground state[2]

optimized at **# APFD/6-311+G (2d, p)** at first excited state

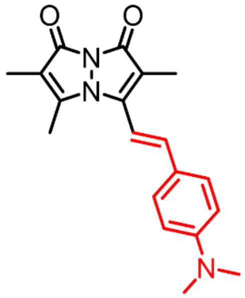

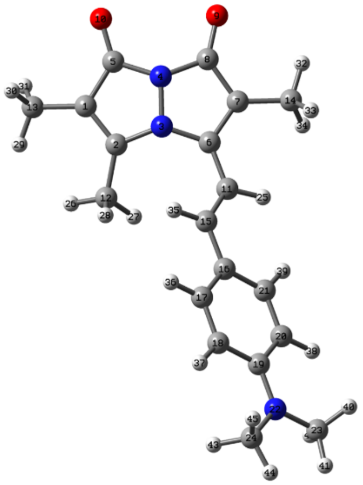

| *Atom* | *X* | *Y* | *Z* | | *Atom* | *X* | *Y* | *Z* |
|---|---|---|---|---|---|---|---|---|
| C | 3.838490 | -1.866286 | 0.248727 | | C | -6.984899 | -1.155657 | -0.687701 |
| C | 2.532485 | -1.459252 | 0.330005 | | H | -0.203427 | 1.826642 | 0.597604 |
| N | 2.437272 | -0.171688 | -0.163348 | | H | 1.711161 | -3.069076 | 1.415728 |
| N | 3.706791 | 0.273117 | -0.449023 | | H | 0.812932 | -1.553045 | 1.586764 |
| C | 4.634638 | -0.762923 | -0.212710 | | H | 0.664944 | -2.506078 | 0.111962 |
| C | 1.635964 | 0.940614 | 0.112467 | | H | 3.694033 | -3.847555 | 1.060826 |
| C | 2.451302 | 2.079154 | 0.059311 | | H | 4.785618 | -3.704332 | -0.317396 |
| C | 3.774969 | 1.687120 | -0.294673 | | H | 5.276868 | -3.084700 | 1.253708 |
| O | 4.800088 | 2.352020 | -0.481970 | | H | 2.890872 | 4.146774 | 0.121159 |
| O | 5.852130 | -0.668363 | -0.390058 | | H | 1.312750 | 3.781216 | -0.590522 |
| C | 0.229147 | 0.927592 | 0.167774 | | H | 1.530820 | 3.682142 | 1.154543 |
| C | 1.366662 | -2.182066 | 0.885217 | | H | -0.176359 | -0.843594 | -0.932726 |
| C | 4.422192 | -3.192720 | 0.580015 | | H | -2.270514 | -1.758497 | -1.508321 |
| C | 2.021623 | 3.491074 | 0.194069 | | H | -4.685957 | -1.830927 | -1.382400 |
| C | -0.609759 | -0.034886 | -0.354673 | | H | -4.618898 | 1.609401 | 1.252532 |
| C | -2.028531 | -0.035505 | -0.227271 | | H | -2.218021 | 1.655445 | 1.125012 |
| C | -2.790716 | -1.019385 | -0.907051 | | H | -6.733628 | 1.861452 | 0.433644 |
| C | -4.156652 | -1.061484 | -0.836130 | | H | -8.015042 | 0.667713 | 0.684555 |
| C | -4.875867 | -0.107636 | -0.061139 | | H | -6.686438 | 0.785117 | 1.847499 |
| C | -4.118377 | 0.872811 | 0.637912 | | H | -6.705264 | -2.156388 | -0.345769 |
| C | -2.751562 | 0.900259 | 0.559570 | | H | -8.041924 | -1.006536 | -0.489034 |
| N | -6.225464 | -0.136669 | 0.009556 | | H | -6.822997 | -1.096529 | -1.768175 |
| C | -6.949351 | 0.848974 | 0.787070 | | | | | |

**Cartesian coordinates of the optimized geometries: 3d**

Cartesian coordinates of **3d** optimized at **# APFD/6-311+G (2d, p)** at ground state

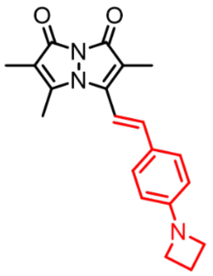

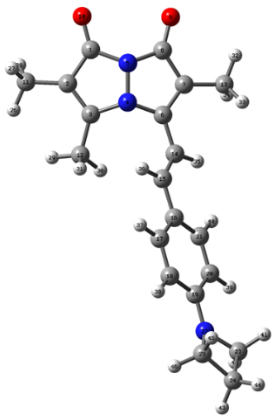

| Atom | X | Y | Z | | Atom | X | Y | Z |
|---|---|---|---|---|---|---|---|---|
| C | 4.90752 | -0.77086 | -0.27943 | | C | -7.94055 | -0.01083 | -0.16106 |
| C | 4.10671 | -1.84911 | 0.259907 | | C | -6.82928 | -0.87155 | -0.80083 |
| C | 2.811959 | -1.44121 | 0.294307 | | H | 3.957877 | -3.81272 | 1.111128 |
| N | 2.705833 | -0.17705 | -0.28876 | | H | 5.140478 | -3.66231 | -0.19155 |
| N | 3.980265 | 0.245254 | -0.60986 | | H | 5.500684 | -2.99608 | 1.396215 |
| C | 1.969062 | 0.952299 | 0.092315 | | H | 1.977269 | -3.03104 | 1.399867 |
| C | 2.782533 | 2.051681 | 0.078333 | | H | 1.022502 | -1.53854 | 1.455436 |
| C | 4.094224 | 1.641307 | -0.35154 | | H | 1.003752 | -2.54653 | 0.009654 |
| O | 5.113402 | 2.286914 | -0.52626 | | H | 3.208221 | 4.126254 | -0.01949 |
| O | 6.112136 | -0.69055 | -0.44627 | | H | 1.491831 | 3.738988 | -0.21883 |
| C | 4.699575 | -3.1482 | 0.667914 | | H | 2.234801 | 3.671418 | 1.380274 |
| C | 1.638129 | -2.17525 | 0.818291 | | H | 0.179778 | 1.73928 | 0.937642 |
| C | 2.40486 | 3.469138 | 0.318219 | | H | 0.132723 | -0.66168 | -0.96708 |
| C | 0.55498 | 0.932323 | 0.316944 | | H | -1.90641 | -1.57324 | -1.59008 |
| C | -0.29276 | 0.067252 | -0.28099 | | H | -4.34658 | -1.75223 | -1.43644 |
| C | -1.72465 | 0.011742 | -0.14604 | | H | -4.36618 | 1.403295 | 1.496792 |
| C | -2.44693 | -0.91825 | -0.91227 | | H | -1.94136 | 1.571278 | 1.337433 |
| C | -3.81935 | -1.02178 | -0.83239 | | H | -6.60731 | 1.718357 | 0.301604 |
| C | -4.54186 | -0.18421 | 0.038633 | | H | -6.94991 | 0.713606 | 1.734318 |
| C | -3.82676 | 0.753386 | 0.816 | | H | -8.63292 | -0.58151 | 0.456607 |
| C | -2.45771 | 0.841905 | 0.721769 | | H | -8.50151 | 0.619274 | -0.84845 |
| N | -5.88479 | -0.28956 | 0.150393 | | H | -6.59763 | -0.61059 | -1.84025 |
| C | -6.83545 | 0.703418 | 0.647582 | | H | -6.93907 | -1.95592 | -0.72102 |

**Cartesian coordinates of the optimized geometries: 3d**

Cartesian coordinates of **3d** optimized at **# APFD/6-311+G (2d, p)** at first excited state

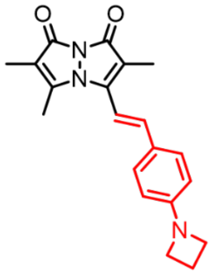

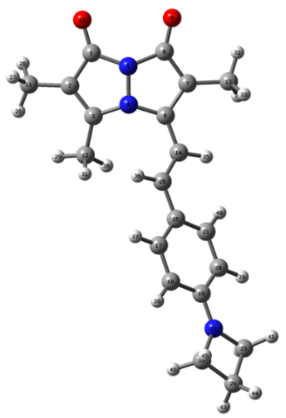

| *Atom* | *X* | *Y* | *Z* | | *Atom* | *X* | *Y* | *Z* |
|---|---|---|---|---|---|---|---|---|
| C | 4.938702 | -0.772418 | -0.219862 | | C | -7.970279 | -0.136008 | 0.070383 |
| C | 4.138211 | -1.876421 | 0.231543 | | C | -6.877250 | -0.944989 | -0.663011 |
| C | 2.833715 | -1.464492 | 0.317713 | | H | 3.986355 | -3.863533 | 1.027881 |
| N | 2.744382 | -0.171444 | -0.162232 | | H | 5.073466 | -3.715208 | -0.353352 |
| N | 4.015316 | 0.270642 | -0.444525 | | H | 5.573849 | -3.110712 | 1.220774 |
| C | 1.948338 | 0.941006 | 0.126407 | | H | 2.003149 | -3.086052 | 1.378984 |
| C | 2.766971 | 2.076307 | 0.082754 | | H | 1.120813 | -1.564468 | 1.582612 |
| C | 4.089022 | 1.682773 | -0.275954 | | H | 0.954494 | -2.488530 | 0.092049 |
| O | 5.116692 | 2.345714 | -0.457612 | | H | 3.214166 | 4.141895 | 0.163840 |
| O | 6.156568 | -0.681821 | -0.398918 | | H | 1.636116 | 3.788834 | -0.554021 |
| C | 4.716120 | -3.208723 | 0.549417 | | H | 1.850636 | 3.673663 | 1.190597 |
| C | 1.664452 | -2.186249 | 0.866590 | | H | 0.109895 | 1.819998 | 0.635818 |
| C | 2.342208 | 3.488954 | 0.229342 | | H | 0.133003 | -0.812008 | -0.957177 |
| C | 0.540223 | 0.929772 | 0.185683 | | H | -1.956025 | -1.688510 | -1.586393 |
| C | -0.299598 | -0.018571 | -0.357426 | | H | -4.403082 | -1.749884 | -1.462630 |
| C | -1.719527 | -0.022147 | -0.229382 | | H | -4.326887 | 1.560495 | 1.316951 |
| C | -2.479959 | -0.976543 | -0.956396 | | H | -1.893813 | 1.607529 | 1.199710 |
| C | -3.846011 | -1.015631 | -0.891585 | | H | -6.806118 | 1.753861 | 0.448154 |
| C | -4.545140 | -0.089129 | -0.072544 | | H | -6.743810 | 0.602566 | 1.807361 |
| C | -3.802221 | 0.858852 | 0.677905 | | H | -8.566047 | -0.715671 | 0.772388 |
| C | -2.435433 | 0.883723 | 0.601846 | | H | -8.626420 | 0.432099 | -0.585786 |
| N | -5.876999 | -0.114643 | -0.008448 | | H | -6.863074 | -0.853467 | -1.753606 |
| C | -6.841637 | 0.694408 | 0.721086 | | H | -6.801094 | -2.002766 | -0.392528 |

**Cartesian coordinates of the optimized geometries: 3e**

Cartesian coordinates of **3e** optimized at **# APFD/6-311+G (2d, p)** at ground state

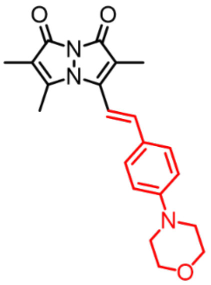


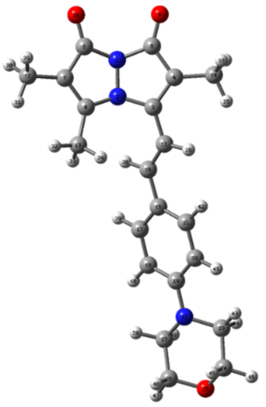

| *Atom* | *X* | *Y* | *Z* | | *Atom* | *X* | *Y* | *Z* |
|---|---|---|---|---|---|---|---|---|
| N | -4.676562 | 0.176351 | -0.579498 | | C | 7.358513 | -1.299676 | -0.233380 |
| N | -3.383537 | -0.193476 | -0.271150 | | C | 5.974150 | -1.046309 | -0.806101 |
| C | -2.679764 | 0.968522 | 0.074253 | | H | -0.884377 | 1.779831 | 0.905108 |
| C | -3.529330 | 2.037887 | 0.048376 | | H | -6.033955 | -3.073582 | 1.536123 |
| C | -4.835254 | 1.570266 | -0.348614 | | H | -5.712066 | -3.748651 | -0.056140 |
| C | -5.562578 | -0.865349 | -0.213987 | | H | -4.479680 | -3.849736 | 1.204576 |
| C | -4.715223 | -1.902155 | 0.335959 | | H | -2.526765 | -2.993290 | 1.451284 |
| C | -3.435565 | -1.448063 | 0.340851 | | H | -1.584261 | -2.477844 | 0.050056 |
| O | -5.876629 | 2.180011 | -0.514571 | | H | -1.635038 | -1.462046 | 1.488724 |
| O | -6.770838 | -0.829904 | -0.364947 | | H | -2.163487 | 3.644271 | 0.437417 |
| C | -1.257832 | 0.980427 | 0.274643 | | H | -3.501717 | 4.056997 | -0.640758 |
| C | -5.255469 | -3.212024 | 0.780945 | | H | -3.790962 | 3.885719 | 1.085626 |
| C | -2.228148 | -2.128624 | 0.860843 | | H | -0.828723 | -0.588161 | -1.037698 |
| C | -3.223293 | 3.477310 | 0.244511 | | H | 1.239496 | -1.423814 | -1.718899 |
| C | 1.030581 | 0.114619 | -0.233673 | | H | 3.654764 | -1.536059 | -1.621377 |
| C | -0.407712 | 0.141236 | -0.349344 | | H | 3.636185 | 1.519871 | 1.422134 |
| C | 1.765689 | -0.774966 | -1.024230 | | H | 1.240549 | 1.635076 | 1.289846 |
| C | 3.145057 | -0.843830 | -0.964421 | | H | 5.509167 | 1.725570 | 1.035045 |
| C | 3.873787 | -0.020793 | -0.086871 | | H | 6.114277 | 1.692602 | -0.626740 |
| C | 3.131382 | 0.877486 | 0.712011 | | H | 7.302510 | 0.278454 | 1.816127 |
| C | 1.759453 | 0.938039 | 0.639931 | | H | 7.992341 | 1.663391 | 0.934493 |
| N | 5.247702 | -0.106890 | 0.033013 | | H | 7.934641 | -1.924686 | -0.917332 |
| C | 6.015574 | 1.105272 | 0.298786 | | H | 7.268559 | -1.822261 | 0.729855 |
| C | 7.392459 | 0.757952 | 0.830765 | | H | 6.055510 | -0.670827 | -1.837810 |
| O | 8.089600 | -0.100984 | -0.053283 | | H | 5.443884 | -1.999078 | -0.831314 |

**Cartesian coordinates of the optimized geometries: 4a**

optimized at **# APFD/6-311+G (2d, p)** at ground state

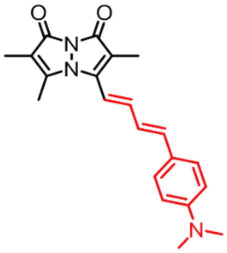

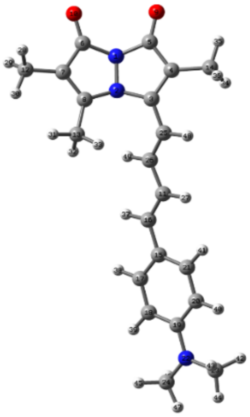

| *Atom* | *X* | *Y* | *Z* | | *Atom* | *X* | *Y* | *Z* |
|---|---|---|---|---|---|---|---|---|
| N | -4.663782 | 0.102787 | -0.618177 | | C | -0.398915 | 0.412444 | -0.117698 |
| N | -3.364054 | -0.165785 | -0.237668 | | H | 1.287874 | 1.422458 | 0.759234 |
| C | -2.770090 | 1.050621 | 0.126395 | | H | -5.905175 | -3.211049 | 1.456119 |
| C | -3.698812 | 2.052373 | 0.043412 | | H | -5.404999 | -3.893056 | -0.086372 |
| C | -4.939719 | 1.485622 | -0.417607 | | H | -4.271634 | -3.862188 | 1.267433 |
| C | -5.487473 | -0.996638 | -0.280347 | | H | -2.397117 | -2.855042 | 1.593738 |
| C | -4.596347 | -1.960552 | 0.328475 | | H | -1.434076 | -2.332295 | 0.210874 |
| C | -3.355582 | -1.412332 | 0.390911 | | H | -1.606234 | -1.268907 | 1.607009 |
| O | -6.016705 | 2.007264 | -0.651266 | | H | -2.698265 | 3.896029 | -0.411109 |
| O | -6.686360 | -1.055066 | -0.491796 | | H | -4.407655 | 4.048962 | 0.020033 |
| C | 0.993390 | 0.600778 | 0.110632 | | H | -3.200652 | 3.742483 | 1.272911 |
| C | -5.060977 | -3.301097 | 0.767244 | | H | 1.566692 | -1.016046 | -1.085660 |
| C | -2.130059 | -1.994321 | 0.982688 | | H | 3.697808 | -1.838506 | -1.569858 |
| C | -3.485601 | 3.509717 | 0.242727 | | H | 6.112414 | -1.814896 | -1.382721 |
| C | 3.356278 | -0.140497 | -0.294587 | | H | 5.834412 | 1.618666 | 1.226438 |
| C | 1.927141 | -0.209886 | -0.446590 | | H | 3.436629 | 1.572220 | 1.022018 |
| C | 4.168285 | -1.076898 | -0.953772 | | H | 7.990409 | 1.947834 | 0.509656 |
| C | 5.542537 | -1.065526 | -0.848695 | | H | 7.895730 | 0.869864 | 1.917931 |
| C | 6.200863 | -0.094650 | -0.059243 | | H | 9.281020 | 0.784789 | 0.825908 |
| C | 5.383990 | 0.852720 | 0.608154 | | H | 8.129907 | -2.067211 | -0.310062 |
| C | 4.014066 | 0.823548 | 0.489314 | | H | 8.232439 | -0.990858 | -1.718589 |
| N | 7.554845 | -0.068643 | 0.056214 | | H | 9.414361 | -0.858900 | -0.412356 |
| C | 8.205245 | 0.936017 | 0.869443 | | H | -1.104291 | 2.076148 | 0.978935 |
| C | 8.366705 | -1.047763 | -0.633045 | | H | -0.679837 | -0.409497 | -0.772365 |
| C | -1.373974 | 1.200138 | 0.395983 | | | | | |

**Cartesian coordinates of the optimized geometries: 4a**

optimized at **# APFD/6-311+G (2d, p)** at first excited state

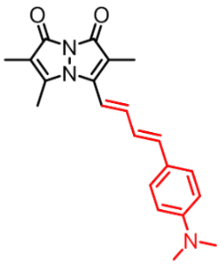


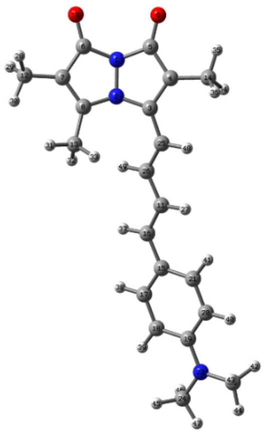

| *Atom* | *X* | *Y* | *Z* | | *Atom* | *X* | *Y* | *Z* |
|---|---|---|---|---|---|---|---|---|
| N | -4.702503 | 0.214445 | -0.527417 | | C | -0.362971 | 0.165949 | -0.143789 |
| N | -3.422173 | -0.157865 | -0.179454 | | H | 1.311432 | 1.279528 | 0.547716 |
| C | -2.704184 | 1.005557 | 0.147355 | | H | -6.179491 | -3.154640 | 1.237100 |
| C | -3.589093 | 2.087051 | 0.085988 | | H | -5.588496 | -3.804639 | -0.286733 |
| C | -4.863758 | 1.623075 | -0.340690 | | H | -4.554841 | -3.846028 | 1.142379 |
| C | -5.587140 | -0.854980 | -0.281631 | | H | -2.635034 | -2.944185 | 1.560638 |
| C | -4.757712 | -1.904412 | 0.251182 | | H | -1.600329 | -2.457543 | 0.219439 |
| C | -3.481179 | -1.429482 | 0.362673 | | H | -1.762668 | -1.404323 | 1.625787 |
| O | -5.915485 | 2.224852 | -0.569949 | | H | -2.525720 | 3.865571 | -0.483646 |
| O | -6.797324 | -0.830003 | -0.502544 | | H | -4.158554 | 4.123730 | 0.147037 |
| C | 1.012546 | 0.368222 | 0.034035 | | H | -2.829448 | 3.729496 | 1.246571 |
| C | -5.290898 | -3.245339 | 0.606354 | | H | 1.661888 | -1.422524 | -0.897077 |
| C | -2.304124 | -2.090187 | 0.970799 | | H | 3.851155 | -2.208135 | -1.235404 |
| C | -3.255221 | 3.521708 | 0.259091 | | H | 6.255174 | -2.008013 | -1.055465 |
| C | 3.390587 | -0.333882 | -0.264134 | | H | 5.753535 | 1.810346 | 0.930672 |
| C | 1.988588 | -0.511106 | -0.400589 | | H | 3.368669 | 1.592220 | 0.743341 |
| C | 4.273412 | -1.328069 | -0.760010 | | H | 7.886844 | 2.126775 | 0.154116 |
| C | 5.633702 | -1.215793 | -0.659229 | | H | 7.838798 | 1.302795 | 1.728566 |
| C | 6.227318 | -0.077901 | -0.044777 | | H | 9.241123 | 1.112122 | 0.667992 |
| C | 5.350176 | 0.925767 | 0.455347 | | H | 8.271544 | -1.940885 | 0.044463 |
| C | 3.992206 | 0.798629 | 0.347836 | | H | 8.313892 | -1.119061 | -1.531208 |
| N | 7.568288 | 0.045759 | 0.059078 | | H | 9.480246 | -0.689608 | -0.273339 |
| C | 8.159776 | 1.210877 | 0.686708 | | H | -0.951869 | 2.021052 | 0.700233 |
| C | 8.450558 | -0.983686 | -0.454197 | | H | -0.683077 | -0.728997 | -0.667405 |
| C | -1.310521 | 1.082963 | 0.283132 | | | | | |
